# Engineered Allosteric RNA Editors Enable Compact, Stimulus-Responsive Post-Transcriptional Circuits

**DOI:** 10.64898/2026.01.29.702378

**Authors:** Alexander M. Marzilli, John T. Ngo

## Abstract

Translational regulation offers a powerful biological control axis with the potential to enable programmable control over synthetic mRNAs. Here, we introduce inducible Deaminases Acting on RNA (iDARs): deaminase domains (DDs) with conditional RNA-editing activities. Using a domain-insertion strategy, we designed autoinhibited enzymes that can be converted into active RNA editors in response to triggers based on small molecules (chemiDARs), intracellular antigens (antiDARs), protease cleavage (lysiDAR), and optical excitation (optiDAR). Coupling these domains with novel stop codon containing RNA substrates enabled conditional protein translation or transcript degradation. Mutational tuning of inositol hexaphosphate (IP_6_)-binding pockets produced tightly regulated deaminases with minimal basal activity, facilitating dose-dependent readthrough translation in response to low-nanomolar drug concentration, with dynamic ranges exceeding 100-fold. By encoding iDARs alongside their substrates, we developed “self-editing” polycistronic transcripts capable of directing translation of encoded proteins in a trigger-dependent manner following delivery to cells as *in vitro* transcribed mRNAs. Overall, iDARs provide a generalizable framework for generating controllable deaminases, enabling the design of post-transcriptional circuits that link biochemical sensing to readouts based on *de novo* translation or mRNA decay.

## INTRODUCTION

Eukaryotic cells convert diverse chemical and biophysical signals into precise molecular outputs, and engineering analogous sense-and-respond systems has been a central objective in synthetic biology. Although transcriptional circuits have served as the dominant framework for achieving such control^1,2^, DNA-based approaches are inherently constrained by transcriptional latency and by fitness costs associated with the persistent maintenance of exogenous DNA sequences^3^. Post-transcriptional regulation offers a faster and more transient alternative^4,5^; however, many existing RNA-based tools exhibit basal activity, limited generalizability, or only modest inducibility, limiting their practical utility compared to DNA-based strategies^6^.

To address these limitations, researchers have developed post-transcriptional strategies^4^ for regulating mRNAs by controlling stability, splicing, or translation, and among these include approaches based on aptazymes^7–9^, riboswitches^9–11^, and engineered RNA-binding protein (RBP)–RNA assemblies^12–15^. While these systems afford a degree of modularity, their successful implementation can be highly context-dependent, and the extent to which they can be readily reconfigured to detect new molecular inputs remains unclear.

More recently, adenosine deaminase acting on RNA (ADAR)^16^ enzymes have been harnessed to control transcript recoding via RNA editing^17–22^, including strategies in which A-to-I editing is used to direct stop-to-sense codon conversions^23–26^. To regulate editing activity, current approaches primarily rely on induced-proximity mechanisms, such as conditionally recruiting active ADAR domains to transcript targets or reconstituting activity through complementation of split enzyme fragments^26–29^. While powerful, these assembly-based strategies often require precise control over the expression and stoichiometry of individual circuit components—particularly for enzyme-based systems like ADAR^30–33^—where achieving low background while maintaining high inducibility is challenging. More broadly, their reliance on multi-component assemblies limits scalability and programmability, constraining the number and diversity of inputs that can be independently sensed and integrated.

To overcome these challenges, we sought to expand the ADAR toolbox by developing inducible Deaminases Acting on RNA (iDARs): allosterically-regulated, single-chain enzymes engineered for conditional RNA editing control. By inserting autoinhibitory peptide–binder pairs into the ADAR2 deaminase domain (ADAR2-DD) scaffold, we generated chimeras that undergo allosteric activation in response to distinct cues, including small molecules (chemiDARs), intracellular proteins (antiDARs), proteolytic cleavage (lysiDARs), and optical excitation (optiDARs). Coupling these iDAR domains with high-fidelity engineered RNA substrates enables stimulus-dependent regulation of protein expression or transcript turnover.

To achieve tight regulation, we optimized individual iDARs through mutational tuning of the inositol hexaphosphate (IP_6_) binding pocket^34^, a cofactor-stabilized region whose modification minimized background and maximized inducible dynamic ranges across constructs. We further demonstrate that iDARs can be encoded within self-editing transcripts, enabling the design of polycistronic mRNAs capable of autonomously directing drug-inducible payload expression within minutes of ligand exposure. Overall, this approach establishes a generalizable framework for transducing biochemical signals into *de novo* protein synthesis or transcript degradation, thereby enabling the design of autoregulatory circuits that can be delivered to cells as *in vitro* transcribed (IVT) mRNAs.

## RESULTS

### High-fidelity substrates for ADAR-dependent protein expression

To link RNA editing to quantifiable protein outputs, we developed “turn-on” reporters for editing-dependent readthrough translation. Building on established strategies^35^, we designed constructs containing editable UAG stop codons and an MS2 hairpin that recruits MCP–ADAR2-DD between 5’ *mCherry* and 3’ *mNeonGreen* (*mNG*) coding regions (**Fig. 1a**). In this design, adenosine-to-inosine (A→I) editing converts the UAGs to UIGs, enabling readthrough translation of mNG.

**Figure 1.**
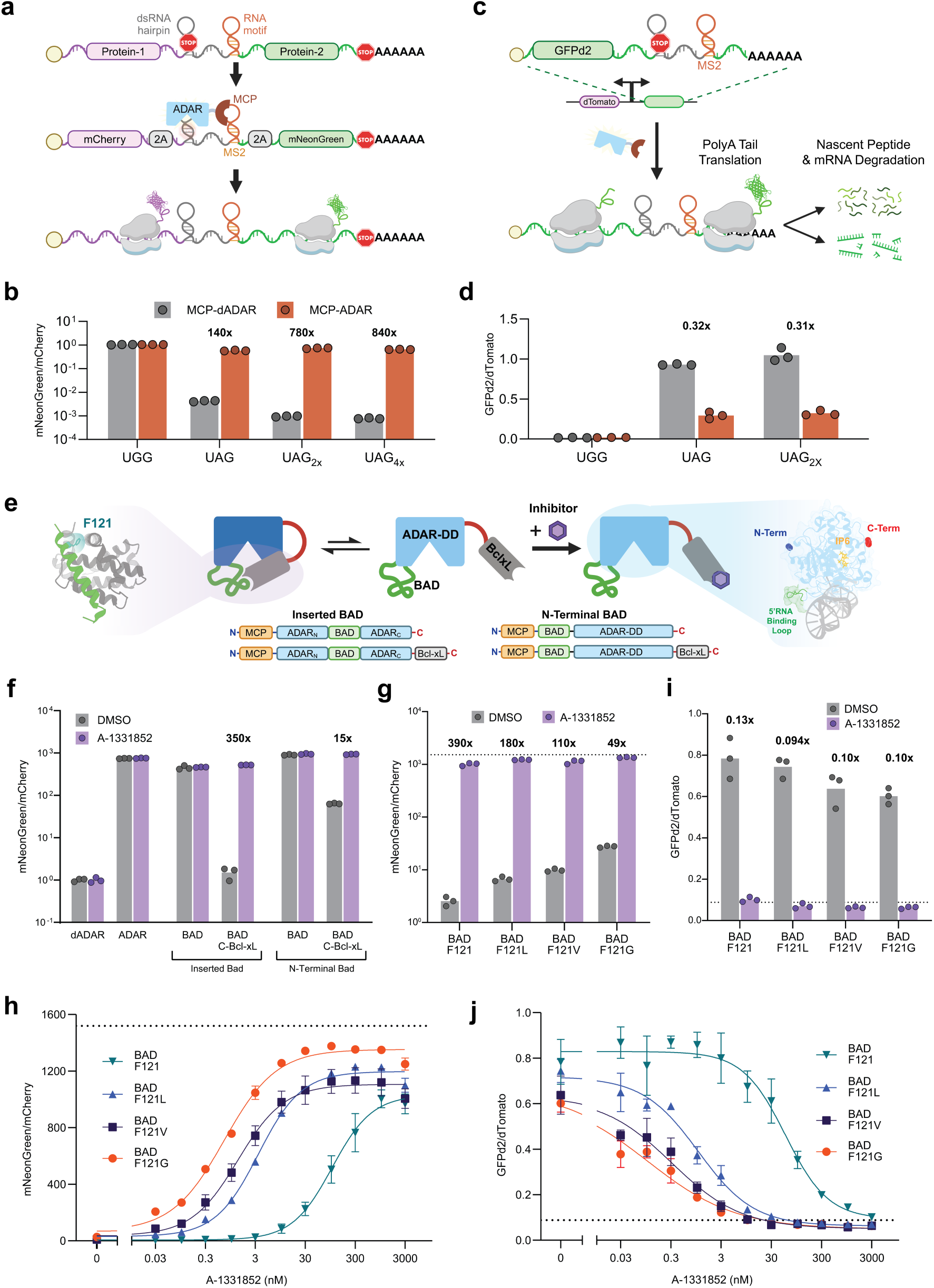
Engineered ADAR reporters and ligand-responsive deaminases enable post-transcriptional control. **a**, Schematic of the ADAR-ON reporter, in which deaminase-mediated conversion of UAG to UIG enables translational readthrough and expression of mNeonGreen (mNG). **b**, Flow cytometry analyses of HEK293FT cells expressing ADAR-ON reporters containing increasing numbers of UAG codons. Cells co-express active (MCP-ADAR, orange) or inactive (MCP-dADAR, gray) deaminases. Bars, mean; points, replicates (n = 3 independent transfections). **c**, Schematic of the ADAR-OFF reporter, in which editing of the UAG stop codon initiates readthrough of the 3’ untranslated region (UTR) and poly(A) tail, triggering nonstop decay and transcript degradation. **d**, Flow cytometry analyses of ADAR-OFF reporters bearing single or dual editable UAG codons co-transfected with MCP-ADAR (orange) or MCP-dADAR (gray). Constitutive readthrough controls in which stop codons are replaced with UGG are also shown. Bars, mean; points, replicates (n = 3 independent transfections); fold-change above. **e**, Design of chemically inducible ADARs (chemiDARs). Intramolecular binding between an inserted BAD peptide and a C-terminal Bcl-xL domain constrains the deaminase into an autoinhibited state; inhibition is relieved by ligand-mediated competitive displacement using a Bcl-xL inhibitor (A-1331852 or A-1, purple hexagon). Insets: crystal structures of BAD/Bcl-xL (PDB: 2BZW^76^) and ADAR2-DD (PDB: 5ED2^31^). **f**, Activity measurements of chemiDAR variants containing the BAD peptide inserted at the 5’ RNA binding loop (Inserted BAD) or fused to ADAR2-DD’s N-terminus (N-terminal BAD). Cells were treated with DMSO or 1 µM A-1331852; fold changes relative to dADAR shown. **g–h**, Ligand-dependent activation of chemiDAR variants bearing C-terminal Bcl-xL domains and inserted mutant BAD(F121X) peptides. Activity was evaluated in cells expressing ADAR-ON reporters. In **(g)**, cells were treated with DMSO or 1 µM A-1331852. Dotted horizontal line represents values measured from cells expressing a constitutively active MCP-ADAR control. Bars, mean; points, n = 3. In **(h)**, dose-dependent measurements were conducted using the ADAR-ON reporter and treatment with the indicated concentrations of A-1331852. Points, mean; error bars, SD; n = 3. **i–j,** ADAR-OFF activity measurements of the BAD(F121X)-containing chemiDAR variants. In **(i)**, cells were treated with DMSO or 1 µM A-1331852. Bars, mean; n = 3. In **(j)**, dose-dependent measurements were conducted using the ADAR-OFF reporter and the indicated concentrations of A-1331852. Points, mean; error bars, SD; n = 3.

Reporter performance was evaluated by co-expressing MCP-fused hyperactive ADAR2-DDs (E488Q, “MCP-ADAR”) versus catalytically inactive controls (E396A, “MCP-dADAR”)^34,36^. We quantified relative readthrough translation as the mNG/mCherry fluorescence ratio by flow cytometry (**Supplementary Fig. 1**). These values were further compared to the efficiency of constitutive readthrough constructs containing UGG sense codons in place of UAG.

We screened sequence variants bearing redundant stop codons, hairpins of varying affinity^37^ or positioning, and distinct protein recruitment motifs. ADAR-dependent mNG expression levels approached that of UGG controls independent of motif architecture or design, suggesting efficient editing across constructs (**Supplementary Fig. 2**). However, the addition of redundant stop codons further minimized background translation (**Supplementary Fig. 2**). We confirmed that editing depended on specific pairing between RNA binding domain (RBD)-DD fusions and cognate protein recruitment motifs (**Supplementary Fig. 2**) and that substrates remained insensitive to overexpressed full-length ADAR1 and ADAR2 (**Supplementary Fig. 3**).

Guided by these results, we designed an optimized “ADAR-ON” reporter comprising two identical substrates, each bearing redundant in-frame UAG pairs (two UAG_2x_, or UAG_4x_) and containing sequence features for minimizing internal initiation (**Supplementary Fig. 4**). This design yielded mNG/mCherry ratios up to 840-fold higher in MCP-ADAR cells compared to inactive MCP-dADAR controls (**Fig. 1b**). Given its favorable properties, we proceeded with this design for subsequent inducible editing assays.

### Stop codon elimination links editing to non-stop decay

Complementing “turn-ON” logic, we designed “turn-OFF” reporters linking RNA editing to non-stop mRNA decay^38,39^. Here, editable UAGs terminate a *gfpd2* reading frame upstream of a modified 3’ UTR lacking additional in-frame stop codons. Upon UAG-to-UIG conversion, ribosomal readthrough into the poly(A) tail triggers transcript turnover and nascent protein degradation^40^ (**Fig. 1c**).

To validate this design, we used a bidirectional CMV promoter to drive transcription of separate *gfpd2* reporter and control *dtomato* transcripts. Co-expression with MCP-ADAR resulted in significantly attenuated GFPd2/dTomato ratios compared to inactive MCP–dADAR controls (**Fig. 1d**). Fluorescence *in situ* hybridization^41^ confirmed that MCP-ADAR expression diminished *gfpd2* transcripts, reducing transcript abundance to comparable levels as constitutively degraded UGG controls (**Supplementary Fig. 5**).

Like the ADAR-ON reporters, the ADAR-OFF design exhibited robustness across constructs with varying stop codon redundancies and demonstrated specificity when paired with distinct RBDs (**Supplementary Fig. 6**). Collectively, these substrates provide orthogonal turn-on and turn-off outputs for conditional RNA editing platforms.

### Engineering chemically inducible adenosine deaminases

With reporters established, we next engineered ADAR2 DDs with drug-gated activities. Drawing on previous drug-gated enzymes^42,43^, we hypothesized that an intramolecular BH3/Bcl-xL interaction could constrain ADAR2-DD in an autoinhibited state relievable by ligand-induced displacement (**Fig. 1e**). Reasoning that substrate loading could be sterically hindered at the RNA-binding interface^31^ (**Fig. 1e**), we inserted a BAD-derived peptide into the splice-variable 5’ RNA binding loop^44^ and fused Bcl-xL to the N- or C-terminus. To evaluate drug sensitivity, we co-expressed the ADAR2-DD chimeras with ADAR-ON reporters in HEK293FT cells with and without Bcl-xL ligands (ABT-737 or A-1331852, “A1”)^45,46^.

Insertion of the BAD peptide alone at the loop did not impair deaminase activity (**Supplementary Fig. 7**). In contrast, C-terminal fusion with Bcl-xL led to strong autoinhibition on the BAD-containing deaminase, with ligand treatment restoring editing activity (**Fig. 1f, Supplementary Fig. 7**). By screening linker lengths we found that efficient autoinhibition could be conferred using linkers shorter than the ∼54 Å distance separating the C-terminus and insertion site in the native ADAR2-DD structure^31,34^, suggesting that the chimeric enzyme adopts a distinct conformation in its autoinhibited form (**Supplementary Fig. 7**). Surprisingly, N-terminal Bcl-xL fusions remained constitutively active independent of linker lengths, despite a shortened distance between the DD’s N-terminus and insertion site (∼31 Å**; Supplementary Fig. 7**)^31,34^.

Given the known role of the C-terminus in IP_6_-mediated folding^34^ and the reported weak to null activity of truncated splice variants^47^, our results suggest that autoinhibition primarily arises due to conformational constraints imposed on the enzyme’s C-terminal region by its fusion to Bcl-xL. Consistent with this hypothesis, constructs bearing C-terminal Bcl-xL fusions together with N-terminally fused, or alternatively inserted BAD peptides within loops distal to the RNA-interface, yielded autoinhibited deaminases with drug-inducible activity (**Fig. 1f, Supplementary Fig. 8**). Extending this architecture to the conserved ADAR1-DD similarly produced deaminases with ligand-controllable activity (**Supplementary Fig. 8**). Together, these findings establish the C-terminal fusion strategy as an effective and generalizable scaffold for engineered allostery, which we term chemically-inducible ADARs (chemiDARs).

### Tuning of chemiDAR sensitivity by peptide-protein mutagenesis

Next we tested whether chemiDARs with enhanced drug sensitivity could be designed by altering the intramolecular interaction between BAD and Bcl-xL binding units. To test this we mutated the F121 residue in the BAD peptide as inserted at the original 5’ RNA binding loop, replacing the residue with progressively destabilizing substitutions (F121, F121L, F121V, F121G)^43^, anticipating that the resulting chemiDARs would have heightened ligand sensitivity.

Dose-response measurements using ABT-737 and A1 revealed that drug sensitivities increased alongside weakened peptide affinities as expected (**Fig. 1g-h, Supplementary Fig. 9**). For A1, EC_50_ values decreased from ∼100 nM (for wild-type BAD) to 3.5 nM for F121L, 1.3 nM for F121V, and 0.71 nM for F121G (**Fig. 1h**). Sensitivity increases were accompanied by concomitant elevations in basal activities, with mNG/mCherry ratios increasing from 2.6-fold above dADAR controls for wildtype BAD to 6.7-fold (F121L), 9.8-fold (F121V), and 28-fold (F121G) (**Fig. 1g**).

These trends persisted in OFF reporter assays, in which A1 treatment reduced GFPd2 levels by up to 10-fold, with mutant-specific sensitivities aligning with those measured using the ON reporter system (**Fig. 1i-j**).

### Modular protein architecture enables diverse chemiDARs

To probe modularity, we tested whether the chemiDAR scaffold could accommodate alternative autoinhibitory peptide-protein pairs. We first tested exchanging the BAD/Bcl-xL domains for homologous domains such as the MS1/Mcl-1 pair (**Fig. 2a**)^48^. The resulting chimeras were effectively autoinhibited and responded to the Mcl-1 ligand S63845 (**Fig. 2b**)^49^. As observed with BAD/Bcl-xL chemiDARs, destabilizing mutations at MS1 residue I17 shifted EC_50_ values from >1 μM for the wild-type and I17V peptide variants to 180 nM and 6.1 nM for the I17A and I17G MS1 mutants, respectively (**Fig. 2c**).

**Figure 2.**
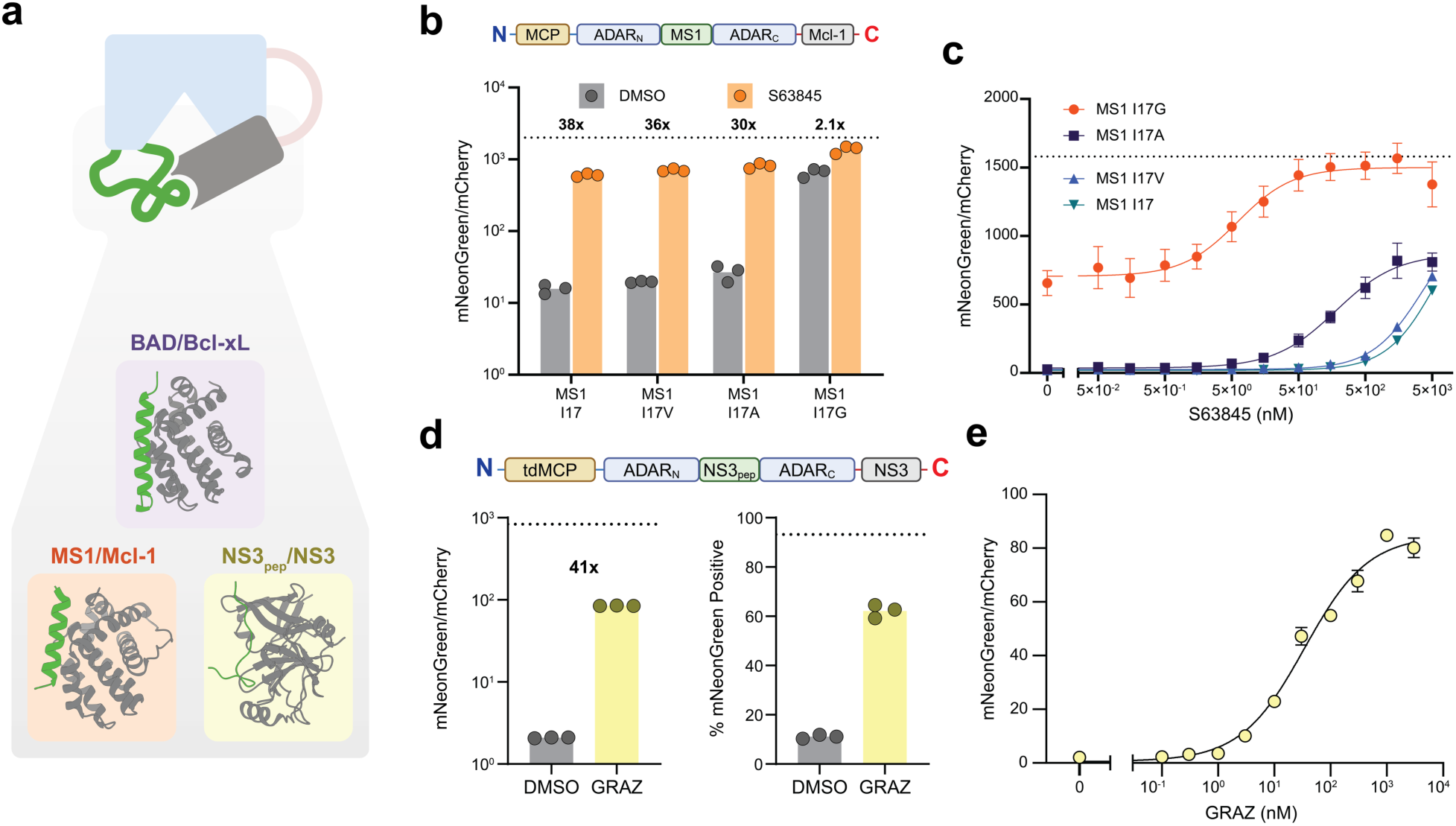
Modular autoinhibitory domains facilitate construction of chemiDARs with distinct drug sensitivities. **a,** Distinct peptide-protein pairs were incorporated into the iDAR scaffold as autoinhibitory domains in place of BAD and Bcl-xL. Structures: BAD/Bcl-xL (PDB: 2BZW^76^), MS1/Mcl-1 (PDB: 5W8F^77^), and HCV NS3_pep_/NS3 protease (PDB: 4A1V^78^). **b,** Schematic of the Mcl-1-based chemiDAR sequence (top) and ADAR-ON activity measurements in cells co-expressing chemiDARs with the indicated (I17X) MS1 peptide mutations (bottom). Cells were treated with DMSO (gray) or 5 µM S63845 (orange). Bars, mean; points, n = 3 independent transfections. **c,** Dose-dependent ADAR-ON measurements using the indicated MS1/Mcl-1 chemiDAR variants and S63845 concentrations. Points, mean; error bars, SD; n = 3; dotted line, mean of three constitutive MCP-ADAR positive control replicates (maximum activation). **d,** Schematic of the NS3-based chemiDAR sequence (top) and ADAR-ON activity measurements in cells DMSO (gray) or 1 µM grazoprevir (GRAZ; yellow). Results are shown as median mNG/mCherry ratios (left) and as percentage of mNG^+^ cells (right). Bars, mean; points, n = 3 independent transfections. **e,** Dose-dependent ADAR-ON measurements for the NS3-based chemiDAR response to the indicated concentrations of GRAZ. Points, mean; error bars, SD; n = 3. Dotted horizontal lines in **(b-d)** represent mean values measured using MCP-ADAR as a positive (max activation; n=3).

To probe the binding requirements for autoinhibition, we tested alternative BH3 sequences with varying binding affinities (K_D_ values ranging from 87–1100 nM)^50^. Bcl-xL- and Mcl-1-based chemiDARs incorporating BIK-, BID-, or mBMF-derived peptides all supported robust autoinhibition and drug-inducible editing (**Supplementary Fig. 10**), together demonstrating that modest-affinity pairs are also suitable for functional chemiDAR generation.

We further generalized the chemiDAR architecture beyond BH3 motifs by C-terminally-fusing the hepatitis C virus NS3 protease domain and inserting a corresponding high-affinity inhibitory peptide (NS3_pep_)^51,52^ (**Fig. 2a**). This design yielded an inducible deaminase with sensitivity to the FDA-approved antiviral drug grazoprevir^53^ (EC_50_ ∼35 nM; **Fig. 2d-e**). Together, these results show that distinct autoinhibitory modules can be leveraged to engineer deaminases that are sensitive to chemically distinct ligands.

### IP_6_-pocket mutagenesis reduces background and expands iDAR designs

Although the tested chemiDARs facilitated ligand-inducible editing, some variants exhibited high levels of basal editing. Because the IP_6_ co-factor stabilizes the active conformation of ADAR2-DD^34^, we hypothesized that mutating residues involved in IP_6_ coordination could bias the enzyme toward autoinhibition and thereby reduce leaky activity (**Fig. 3a**).

**Figure 3.**
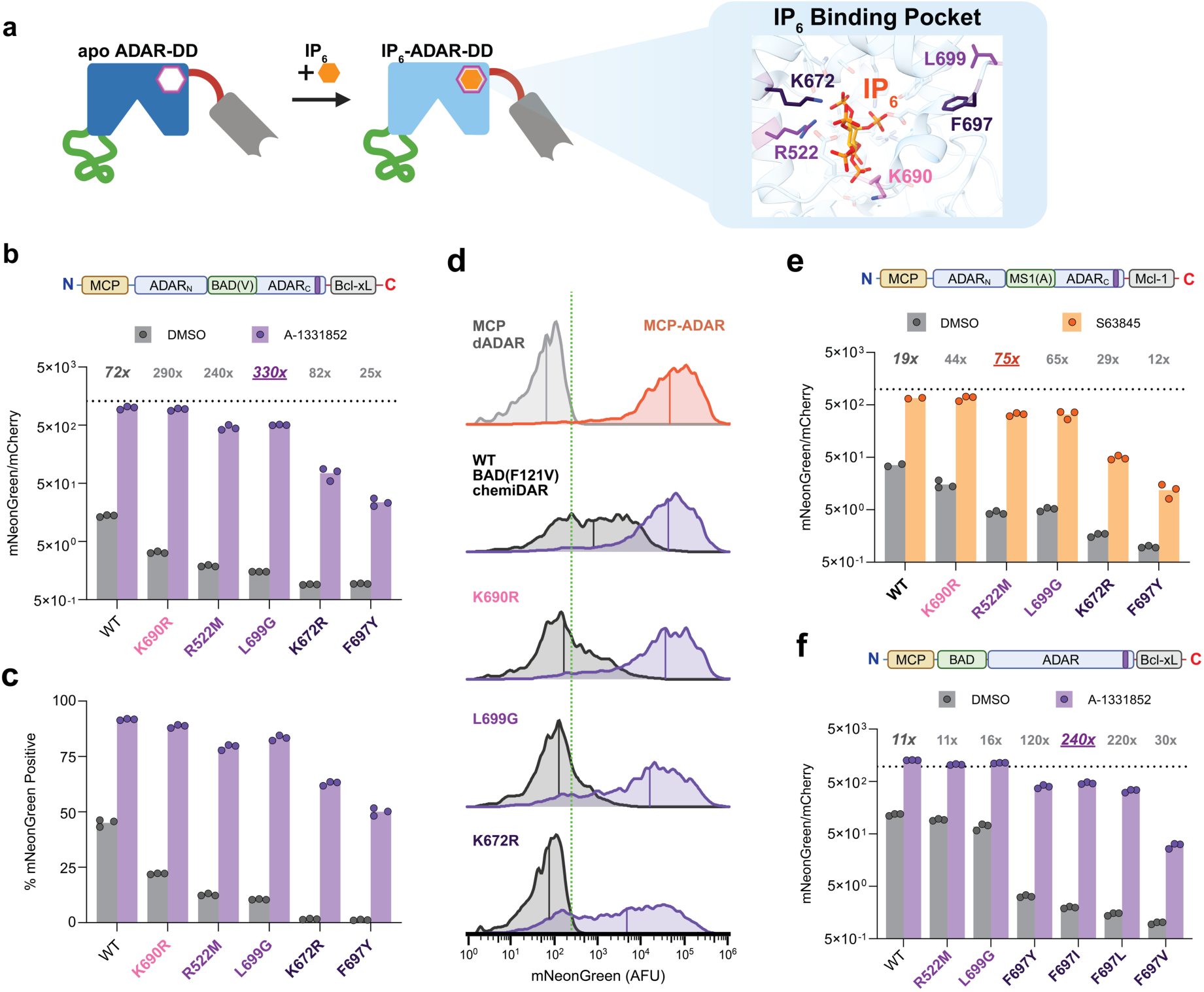
Tuning the ADAR2 IP_6_-binding pocket reduces background and improves inducible control. **a,** ADAR2-DD requires the binding of an IP_6_ co-factor (orange) to become catalytically active. Selected residues stabilizing the C-terminal IP_6_-binding pocket shown (inset: R522, K672, K690, F697, L699; PDB: 5ED2^31^). **b–d,** Activities of BAD(F121V)/Bcl-xL chemiDAR variants bearing the indicated IP_6_-pocket mutations. Measurements were conducted using the ADAR-ON reporter in HEK293FT following treatment with DMSO (gray) or 1 µM A-1331852 (purple). Values in **(b)** represent mNG/mCherry ratios. Data in **(c)** represent percentage of mNG^+^ cells. Bars, mean; points, *n* = 3 independent transfections. Representative histograms shown in **(d)** for results in **(b)** and **(c)**. Distributions of mNG intensities are shown. Vertical green dotted line represents threshold value for defining cells as mNG^+^; median mNG values for each trace are shown as solid lines (AFU). **e,** Activities of MS1(I17A)/Mcl-1 chemiDAR variants bearing the IP_6_-pocket mutations. Measurements were conducted using the ADAR-ON reporter in HEK293FT following treatment with DMSO (gray) or 1 µM S63845 (orange). Values represent mNG/mCherry ratios. Bars, mean; points, *n* = 3 independent transfections (other than n=2 for WT). **f,** Activity measurement of chemiDAR variants based on ADAR2-DD with an N-terminally fused BAD(F121) and C-terminal Bcl-xL. Sequences bearing the indicated IP_6_-pocket mutations were evaluated using the ADAR-ON reporter with fluorescence quantification following treatment with DMSO (gray) or 1 µM A-1331852 (purple). Values represent mNG/mCherry ratios. Bars, mean; points, *n* = 3 independent transfections. Dotted horizontal lines in (b-d) represent mean values measured using MCP-ADAR as a positive (max activation; n=3).

We introduced 22 mutations at residues involved in IP_6_ coordination (R400, R522, S531, Y658, K672, K690) or in C-terminal stability (V688, F697, L699). A prior screen informed our selection of mutations to R400, R522 and S531^33^. In the leaky BAD(F121V)/Bcl-xL chemiDAR, 17 of the tested mutants retained drug responsiveness while exhibiting reduced basal activity, with effects that were additive (**Supplementary Fig. 11**). Highly perturbative substitutions, including K672R and F697Y, nearly abolished background but also diminished maximal activation, whereas other mutations, such as K690R, produced more modest reductions in basal editing (**Fig. 3b-d**). A subset of variants, including R522M and L699G, exhibited lowered basal activities (reduced to near-dADAR levels) and robust drug-inducible editing, leading to dynamic ranges exceeding 300-fold (**Fig. 3b**).

This mutational strategy proved portable to other chemiDARs, consistently reducing basal editing. For example, C-terminal mutations reduced background in the MS1(I17A) chemiDAR, improving yields from 19- to 75-fold (**Fig. 3e, Supplementary Fig. 12**). Mutagenesis also rescued the performance of designs based on N-terminal BAD fusions, with strong mutations (e.g., F697Y) suppressing background and increasing induction yields from 11- to 240-fold (**Fig. 3f**, **Supplementary Fig 13**). Collectively, these results demonstrate that targeted mutagenesis of the IP_6_-binding pocket reduces background editing and enables more stringent and selective chemiDAR control.

### Compact and stringent self-editing circuits using chemiDARs

To minimize construct complexity, we next tested whether chemiDARs and their substrates could be encoded together within single-gene constructs. We designed “self-editing” sequences in which a constitutively translated chemiDAR-mCherry fusion precedes UAG-containing cassettes that conditionally drive expression of downstream proteins (**Fig. 4a–b**).

**Figure 4.**
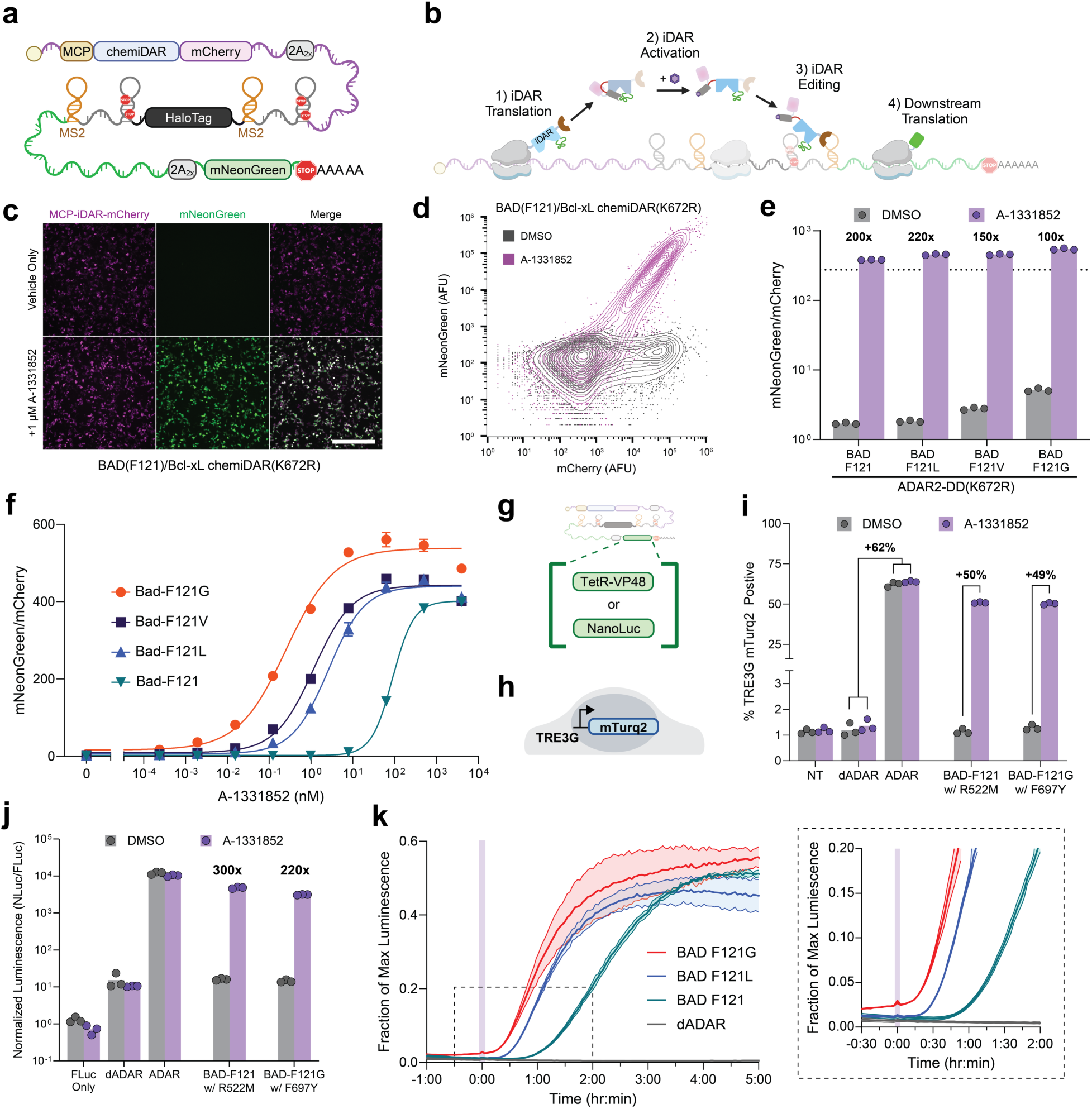
Self-editing iDAR transcripts enable rapid and sensitive ligand-dependent gene expression. **a,** Schematic depicting the sequence organization of a self-editing polycistronic chemiDAR construct. **b,** MCP-chemiDAR-mCherry is constitutively expressed via cap-dependent translation. Ligand-induced chemiDAR activation leads to editing of the UAG substrates, facilitating readthrough translation of proteins encoded by downstream reading frames. **c,** Fluorescence images of cells expressing a self-editing transcript encoding an MCP-chemiDAR-mCherry variant based on BAD(F121)/Bcl-xL and containing the K672R IP_6_ pocket mutation. mNG expression is detected in cells treated with 1 µM A-1331852. Scale bar, 500 µm. **d,** Representative flow cytometry contour plots showing mCherry vs. mNG intensities. HEK293FT cells expressing the construct from **(c)** were analyzed following treatment with DMSO (gray) or 1 µM A-1331852 (purple). **e,** Quantified mNG/mCherry ratios from cells expressing the construct from **(c–d)** along with variants carrying (F121X) BAD mutations in addition to the K672R IP_6_-pocket mutation. Cells were analyzed by flow cytometry following treatment with DMSO (gray) or 1 µM A-1331852 (purple). Bars, mean; points, biological replicates (*n* = 3). **f,** Dose-response measurements of cells expressing the constructs from **(e)** following treatment with the indicated concentrations of A-1331852. Points, mean; error bars, SD; *n* = 3 independent transfections. **g,** Schematics depicting the design of self-editing constructs encoding alternative payloads, in which the synthetic transcription factor TetR-VP48 or NLuc-based reporter is encoded in place of mNG. **h,** Schematic of HEK293FT line containing TRE3G-mTurq2 reporter. **i,** Flow cytometry measurement of mTurq2 expression in *TRE3G-mTurq2* reporter cells, as induced via editing-mediated TetR-VP48 expression. Self-editing circuits encoding MCP-chemiDAR-mCherry variants based on the indicated BAD(F121)/Bcl-xL variants and containing the specified IP6-pocket mutations were expressed in HEK293FT cells. mTurq2 levels were quantified following overnight treatment with DMSO or 1 µM A-1331852. Cells containing self-editing constructs based on MCP-ADAR-mCherry and MCP-dADAR-mCherry were analyzed as controls. NT represents non-transfected reporter cells. Values are shown as percentage of mTurq2+ cells, as gated based on mCherry expression. Bars, mean; points, *n* = 3 independent transfections. **j,** Normalized NLuc activity measurements from HEK293FT cells expressing self-editing constructs in which NLuc is encoded. End-point measurements were conducted using cells treated with DMSO or 1 µM A-1331852. FLuc encoding plasmids were used to normalize for variability in transfections, with the FLuc only condition being singly transfected controls with no NLuc expression potential. Bars, mean; points, *n* = 3 independent transfections. **k,** Real-time NLuc activity measurements from cells expressing variants of the self-editing construct from (i) bearing the indicated (F121X) BAD mutations (in addition to the existing K672R IP_6_-pocket mutation). Transfected HEK293FT cells were grown overnight prior real-time luminescence analysis. Cells were treated with 1 µM A-1331852 at t=0 (purple bar). NLuc activity is detected within 15-30 min and plateauing at ∼5 h following drug treatment. Inset, magnified view of the area indicated by the dashed lines. Traces, mean ± SD of n=3 technical replicates for A-1331852 treated; values are normalized to positive control self-editing construct based on MCP-ADAR and compared to a negative control based on MCP-dADAR (black trace).

When tested in HEK293-FT cells, single-component BAD(F121)/Bcl-xL circuits supported constitutive chemiDAR-mCherry expression and drug-inducible mNG translation (**Supplementary Fig. 14**). A combinatorial screen of four BAD(F121X) variants with five IP_6_-pocket substitutions showed that pocket mutations consistently modulated background and drug-induced mNG levels (**Supplementary Fig. 14**). Notably, introducing the K672R mutation reduced background editing across BAD(F121X) variants while maintaining enhanced A1 sensitivities (**Fig. 4c–f**). Similar trends involving IP_6_-pocket mutations were observed for MS1/Mcl-1 and NS3pep/NS3 single-vector constructs, confirming the benefit of combining background-reducing IP_6_-pocket mutations with sensitivity-enhancing intramolecular binding substitutions (**Supplementary Fig. 15**).

The self-editing architecture further supported sensitive outputs beyond fluorescent protein translation (**Fig. 4g**). Specifically, circuits driving the TetR-VP48 transcription factor activated transcription in a stable reporter line containing an integrated *TRE3G-mTurq2* reporter gene (**Fig. 4h)**, with chemiDAR induction shifting the fraction of mTurq2-positive cells from ∼1% to ∼51% of the population (**Fig. 4i**). Similarly, circuits encoding NanoLuciferase (NLuc) yielded up to 300-fold induction with uninduced cells exhibiting comparable luminescence as those containing catalytically inactive dADAR controls (**Fig. 4j**).

### Rapid self-editing dynamics of chemiDAR logic

A key advantage of post-transcriptional regulation is the ability to bypass transcriptional latency^4,5^. To quantify this, we measured the activation kinetics of the self-editing NLuc- encoding circuit in HEK293FT cells. Real-time luminescence measurements revealed rapid responses upon A1 addition, with NLuc activity becoming detectable within 15-30 minutes and stabilizing within ∼5 hours (**Fig. 4k**). Notably, variants with weaker peptide-protein interactions exhibited accelerated onsets, likely due to increased unbinding kinetics as was previously demonstrated^43^. These results confirm that self-editing chemiDARs enable rapid response times, consistent with the expected dynamics of a post-transcriptionally acting system.

### Antigen-responsive iDARs (antiDARs) enable protein-sensitive circuits

We next sought to enable protein-responsive autonomous circuits by adapting the iDAR scaffold to detect intracellular ligands. We thus engineered “antigen-inducible” ADARs (antiDARs) in which an intramolecular epitope-binder interaction autoinhibits the ADAR2-DD, while competitive binding to a target antigen activates it (**Fig. 5a-b**).

**Figure 5.**
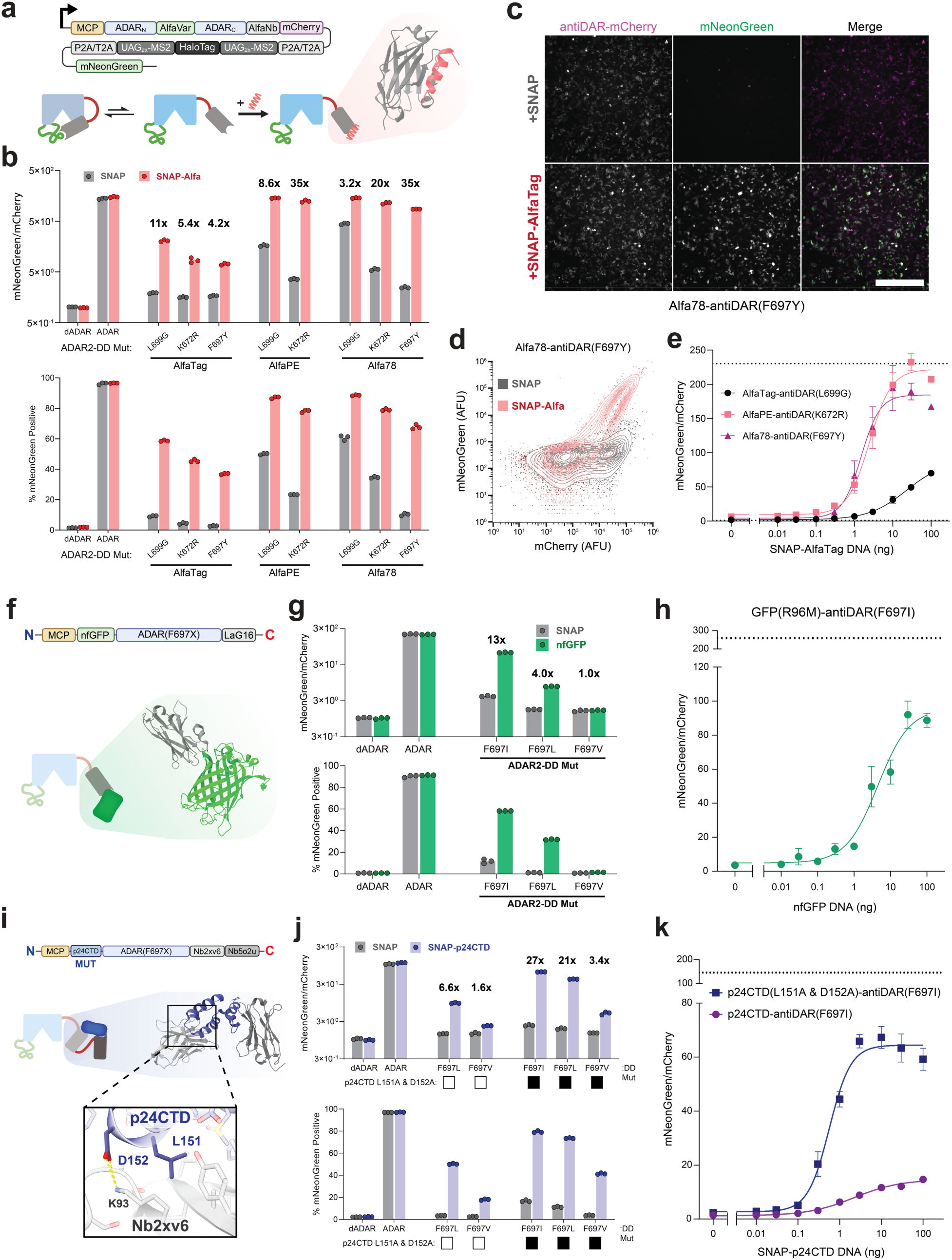
Antibody- and epitope-derived autoinhibitory domains for protein-responsive antiDAR circuits. **a,** Sequence organization of self-editing constructs encoding an MCP-antiDAR-mCherry based on the AlfaTag-AlfaNb antigen-nanobody pair (top) and schematic depicting antiDAR activation via competitive displacement by AlfaTag (bottom). Inset: crystal structure of AlfaTag (red) bound to AlfaNb (gray, PDB: 6I2G^54^). **b,** Flow cytometry measurements of HEK293FT cells expressing self-editing constructs encoding antiDARs based on AlfaNb in combination with the indicated AlfaTag derivatives and indicated ADAR2-DD mutations. Measurements were performed by co-expressing a SNAP-AlfaTag fusion (red) or an unfused SNAP domain (gray) as an antigen-lacking control, with control constructs based on MCP-ADAR-mCherry and MCP-dADAR-mCherry. Displayed values represent mNG/mCherry ratios (top) and percentages of mNG⁺ cells (bottom). Bars, mean; points, *n* = 3 independent transfections. **c,** Representative fluorescence microscopy images of transfected HEK293FT cells expressing a self-editing AlfaTag/AlfaNb-antiDAR(F697Y) construct. Emissions from MCP-antiDAR-mCherry and editing-induced mNG expression are shown. Cells co-expressing the unfused SNAP domain (top) and the SNAP-AlfaTag fusion (bottom) are shown. Scale bar, 500 µm. **d,** Representative flow-cytometry contour plots of cells expressing self-editing Alfa78/AlfaNb-antiDAR(F697Y) constructs. Plots for cells co-expressing SNAP-AlfaTag (pink) or the SNAP control (gray) are shown. **e,** Dose-dependent activation of the indicated self-editing constructs from **(b)** in cells expressing increasing amounts of SNAP-AlfaTag. Points, mean; error bars, SD; *n* = 3 independent transfections. **f,** Sequence organization of the GFP-sensitive antiDAR containing autoinhibitory domains based on an N-terminal non-fluorescent GFP mutant (nfGFP - R96M), and a C-terminal anti-GFP nanobody (LaG16 - top). Competitive binding to nfGFP leads to antiDAR activation (bottom). Inset: GFP (green) bound to LaG16 (gray) (PDB: 6LR7^79^). **g,** Flow cytometry measurements of self-editing nfGFP/LaG16-antiDARs constructs. Variants bearing the indicated ADAR2-DD mutation (F697X) were tested in cells co-expressing nfGFP or a SNAP domain control. Values represent mNG/mCherry ratios (top) and percentage of mNG⁺ cells (bottom). Bars, mean; *n* = 3 independent transfections. **h,** Dose-dependent activation of nfGFP/LaG16-antiDAR(F697I) variant from **(g)** in cells co-transfected with increasing amounts plasmid encoding GFP(R96M). Points, mean; error bars, SD; *n* = 3 independent transfections. **i,** Sequence organization of self-editing constructs encoding an MCP-antiDAR containing autoinhibitory units based on the p24CTD antigen and corresponding tandem nanobody fusions (Nb2XV6 and Nb5O2U; top). Inset: crystal structures of p24CTD (blue) bound to Nb2XV6 (gray; PDB: 2XV6^60^) and Nb5O2U (dark gray; PDB: 5O2U^59^); enlarged Nb2XV6-p24CTD interface with critical residues labeled. **j,** Flow cytometry measurements of self-editing p24CTD-antiDARs constructs in cells co-expressing SNAP-p24CTD (blue) or unfused SNAP as a control (gray). Top, mNG/mCherry ratios; bottom, percentage of mNG⁺ cells. Bars, mean; *n* = 3 independent transfections. **k,** Dose-dependent activation of the p24CTD- and p24CTD(L151A & D152A)-containing antiDARs in cells containing increasing amounts of the SNAP-p24CTD antigen fusion. Points, mean; error bars, SD; *n* = 3 independent transfections.

As an initial model, we tested the AlfaTag/AlfaNb pair^54^, inserting the AlfaTag peptide into ADAR2-DD at the RNA-binding loop and fusing AlfaNb to the C-terminus (**Fig. 5a**). We confirmed the resulting antiDAR was autoinhibited, and measurements made under transient antigen co-expression verified its sensitivity to AlfaTag-mediated activation (**Supplementary Fig. 16**). Substitution of the inserted AlfaTag with weakened-affinity variants (AlfaPE and Alfa78^55^) generated antiDARs with enhanced sensitivity to AlfaTag-mediated activation, consistent with our chemiDAR results (**Supplementary Fig. 16**). To evaluate whether antigen avidity could further enhance sensitivity, we fused the antiDAR C-terminus to a second Nb that recognizes GFP (VHHGFP4)^56^. Tests against a non-fluorescent GFP(R96M)^57^ (nfGFP)-AlfaTag fusion antigen verified the enhanced sensitivity of the dual Nb construct (**Supplementary Fig. 16**), thus establishing avidity-enhanced displacement as an additional strategy for boosting antiDAR performance.

In self-editing circuits, weaker epitope-nanobody interactions supported higher maximal activation but required ADAR2-DD mutations to suppress background (**Fig. 5b**). The Alfa78(F697Y) variant resolved this trade-off, yielding robust activity in flow cytometry and live-cell imaging assays (**Fig. 5c-d**). Titration assays confirmed that the weaker interacting pairs (AlfaPE and Alfa78) enabled activation at lower antigen thresholds relative to the high-affinity AlfaTag (**Fig. 5e**).

To assess generality beyond AlfaTag, we tested additional antigen-antibody pairs. GFP-responsive designs utilizing an N-terminal antigen (nfGFP) and C-terminal LaG16 nanobody^58^ fusion supported selective, dose-dependent activation (**Fig. 5f–h, Supplementary Fig. 17**). We further demonstrated applicability to disease-associated targets by developing antiDARs against the HIV capsid subunit p24 (**Fig. 5i, Supplementary Fig. 18**). High-avidity p24CTD-antiDARs were constructed using p24 in combination with C-terminally fused dual Nbs targeting distinct p24 epitopes^59,60^ (**Fig. 5i**). ADAR-ON measurements in antigen-expressing cells confirmed the antiDAR’s sensitivity to soluble p24CTD, and adding p24-Nb interface mutations substantially enhanced antiDAR sensitivity without increasing leakiness (**Fig. 5j–k, Supplementary Fig. 18**). Together, these results establish antiDARs as a modular platform for designing protein-sensitive RNA editing enzymes, extending iDAR control to inputs based on peptides, proteins, and pathogen-encoded antigens.

### Protease and light-inducible iDARs expand input modalities

To diversify the set of biochemical inputs that could be used to direct editing, we engineered iDARs activated by the cleavage of autoinhibitory domains (**Fig. 6a**), paralleling prior dissociation-based activation strategies^29,61,62^. To test the feasibility of proteolysis-induced ADARs (“lysiDARs”), we inserted TEV protease (TEVp) cleavage sites between ADAR2-DD and C-terminal autoinhibitory domains, hypothesizing that cleavage would relieve autoinhibitory strain (**Fig. 6a-b**). Screening designs identified N-terminal BAD and C-terminal Bcl-xL fusions as the most effective lysiDARs (**Supplementary Fig. 19**). In self-editing circuits, lysiDAR co-expression with TEVp yielded ∼170-fold induction in NLuc luminescence assays, matching the induction levels achieved via drug-induced activation of the same iDAR construct (**Fig. 6b–c, Supplementary Fig. 20**). Further tests confirmed the scaling of editing levels against increasing TEVp doses (**Fig. 6d**) and the sensitivity of the lysiDAR to activation via rapamycin-mediated split-TEVp reconstitution (**Supplementary Fig. 20**)^63,64^.

**Figure 6.**
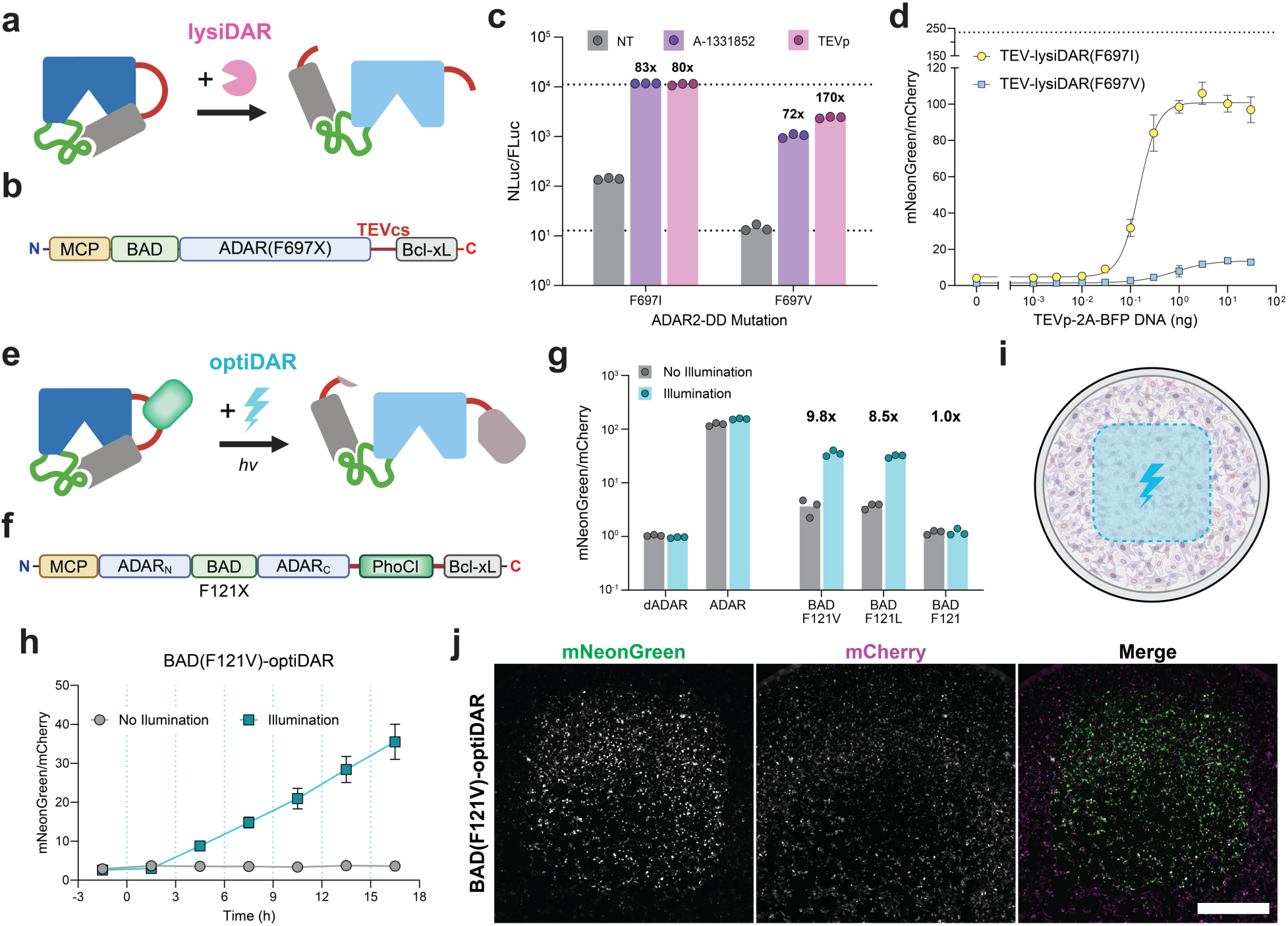
Protease- and light-activated iDARs enabled by cleavable linkages. **a,** Schematic depicting the design and activation of a protease-activated iDAR (lysiDAR). A protease (pink sector) cleaves a substrate sequence embedded within a labile linker (red), thereby facilitating antiDAR activation. **b,** Sequence organization of lysiDAR proteins containing BAD-Bcl-xL autoinhibitory domains at each terminus. A TEV cut site (TEVcs) is used as a linker to connect ADAR-DD and the C-terminally fused Bcl-xL, producing a construct sensitive to TEV protease-mediated activation or drug-mediated displacement. **c,** Activity measurement of the indicated lysiDARs as self-editing constructs encoding NLuc. Normalized NLuc activities were quantified from transfected HEK293FT cells under basal conditions (gray), following treatment with 1 µM A-1331852 (purple), or under co-transfection with TEV protease encoding plasmids (pink). A co-transfection marker based on an FLuc encoding plasmid was used for luminescence normalization. Dotted lines, constitutive MCP-ADAR (positive control; top) and MCP-dADAR (negative control; bottom). Bars, mean; points, *n* = 3 biological replicates. **d,** Flow cytometry measurement of editing-dependent mNG levels in HEK293FT cells co-transfected with self-editing circuits encoding the indicated lysiDARs-mCherry fusions in combination with the specified amounts of a plasmid encoding TEVp-2A-BFP. Values represent mNG/mCherry ratios. Points, mean; error bars, SD; *n* = 3. **e,** Schematic of the optiDAR design containing the photocleavable protein domain PhoCl (pale green). Upon light excitation (cyan bolt), PhoCl undergoes photolytic chromophore cleavage, leading to complex dissociation (separated gray segments) and optiDAR activation. **f,** Sequence organization of PhoCl-enabled optiDARs in which PhoCl is inserted in between ADAR2-DD and the C-terminally fused Bcl-xL. **g,** End-point fluorescence measurements of editing-induced mNG levels in HEK293FT cells co-expressing the indicated optiDAR variants in combination with the ADAR-ON reporter. Values represent mNG/mCherry ratios, as quantified from cell images following violet light illumination (cyan bars) or from light protected cells as a control (gray bars). Bars, mean; points, *n* = 3. **h,** Activation kinetics of a BAD-F121V-based optiDAR in HEK293FT cells co-expressing the ADAR-ON reporter. Cells were subjected to repeated pulse illuminations at 3 h intervals (dotted-cyan line). Values represent mNG/mCherry ratios, as quantified from cell images captured at the indicated times between pulse illuminations. Mean ± SD; *n* = 3. **i,** Schematic depicting spatially-restricted light-mediated OptiDAR activation. Only cells within the indicated region (blue box) are subjected to light-mediated OptiDAR activation. **j,** Fluorescence images depicting HEK293FT expressing the optiDAR variant from (**h**) in combination with an ADAR-ON reporter encoding mCherry and an editing-dependent mNG. Images were acquired at 18 h following optiDAR activation via fluorescence microscopy. Representative image showing mCherry (magenta) and spatially-restricted, editing-induced expression of mNG (green). Image representative of n = 3. Scale bar, 1000 µm.

For optically-induced ADARs (optiDARs), we mirrored this lysis-based activation by using the photocleavable protein PhoCl^65^ to induce light-dependent dissociation (**Fig. 6e**). Violet light stimulation of cells co-transfected with ADAR-ON and optiDAR constructs resulted in spatially-defined mNG expression, limited to cells residing within illuminated wells regions (**Fig. 6f–j, Supplementary Fig. 21**). Overall, lysiDAR and optiDAR broaden the set of inputs that can be used to control iDAR activity, and their successful development further supports the generalizability of our autoinhibition-installation and -release strategies.

### Self-editing iDAR circuits function as *in vitro* transcribed mRNA

Because iDARs operate post-transcriptionally, self-editing circuits should remain functional when delivered as *in vitro* transcribed (IVT) mRNA (**Fig. 7a**). To test this, we transfected HEK293FT cells with ARCA-capped and poly(A)-tailed self-editing transcripts encoding BAD/Bcl-xL-based chemiDARs and quantified editing-mediated NLuc activities following A1 treatment. BAD/Bcl-xL variants retained ligand-dependent activation and sensitivity as IVT mRNA, with the most sensitive variant, BAD(F121G)/Bcl-xL ADAR2-DD(F697Y), exhibiting an EC_50_ of ∼12 nM (**Fig. 7b-c**).

**Figure 7.**
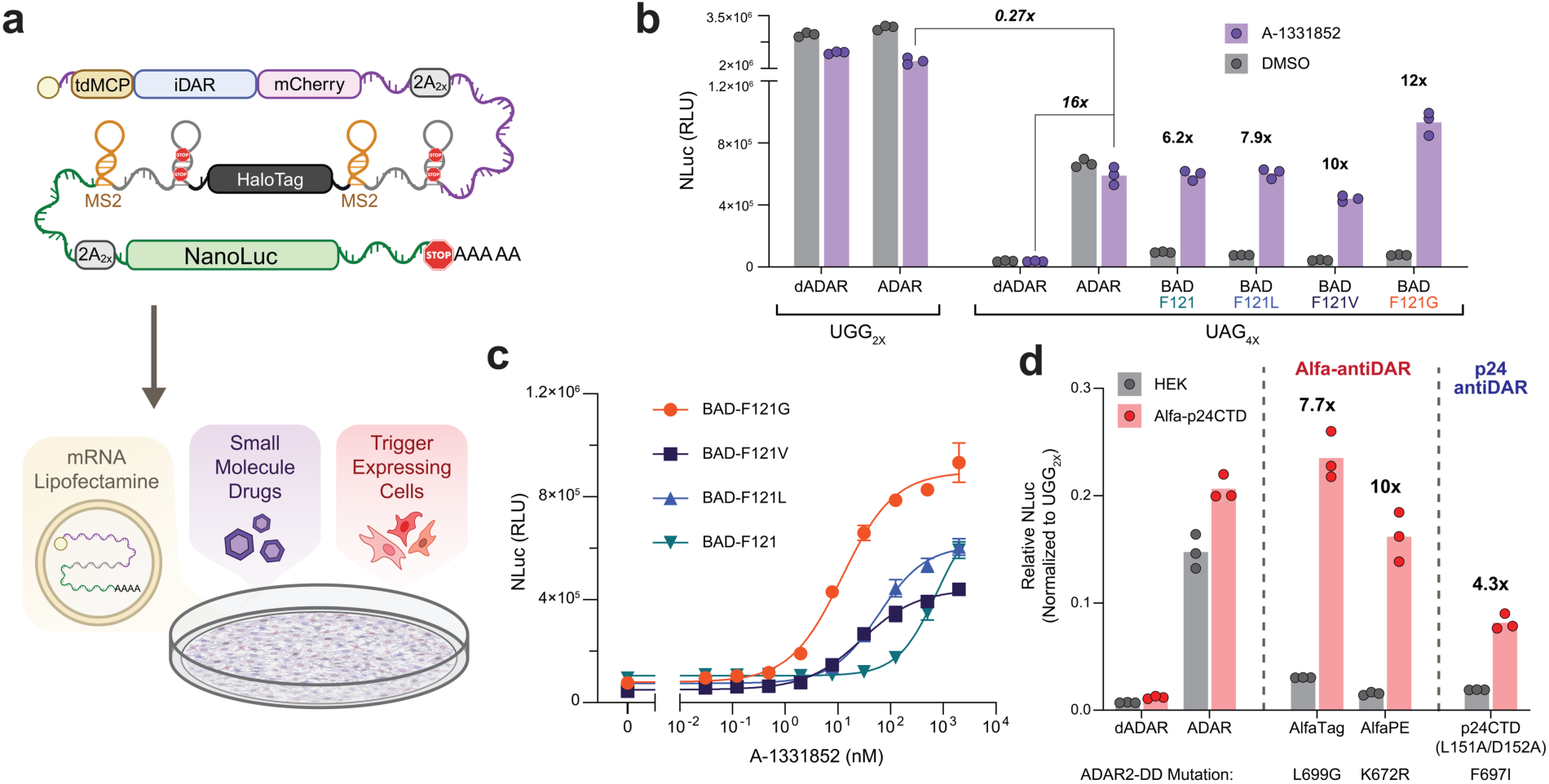
Self-editing iDAR circuits function as in vitro transcribed mRNA. **a,** Schematic depicting the sequence organization of a self-editing iDAR mRNA (top). Transcripts are delivered to cells as IVT mRNA, enabling ligand- or antigen-dependent translation of a NanoLuciferase (NLuc)-based reporter enzyme. **b,** Ligand-mediated activation of the BAD(F121X)/Bcl-xL chemiDAR(F697Y) variants. HEK293FT cells were transfected with self-editing IVT mRNAs and treated with DMSO (gray) or 1 µM A-1331852 (purple). Cells were assayed by luminescence detection at 24 h post-transfection. Bars, mean; points, *n* = 3 independent transfections. Fold-changes between selected conditions shown. **c,** Dose-response of cells transfected with the IVT mRNAs from **(b)** following treatment with the indicated concentrations of A-1331852. Points, mean; error bars, SD; *n* = 3 independent transfections. **d,** Antigen-mediated activation of self-editing IVT mRNAs following delivery to HEK293FT cells expressing a p24CTD-AlfaTag fusion (red) or unmodified (antigen-lacking) control cells. AntiDAR variants and corresponding ADAR2-DD mutations are shown for each tested circuit, including AlfaTag-antiDAR(L699G), AlfaPE-antiDAR(K672R), and p24CTD(L151A/D152A)-antiDAR(F697I). NLuc activities were measured via luminescent detection with normalization to UGG-containing positive controls. Bars, mean; points, *n* = 3 independent transfections.

To next test protein-responsive circuits, we transfected cells stably expressing AlfaTag-p24CTD fusions with antiDAR transcripts. Both AlfaTag- and p24CTD-directed antiDARs produced significantly higher NLuc signals in antigen-expressing cells than in non-expressing control cells (**Fig. 7d**), confirming that the circuits maintain their ligand specificity as IVT mRNAs. These results demonstrate that iDAR circuits function effectively as self-contained, post-transcriptional regulators in an mRNA format.

## DISCUSSION

This work introduces iDARs as a modular and generalizable platform for engineering allosterically regulated RNA editors. By embedding autoinhibitory protein-peptide interactions within the ADAR2-DD scaffold, we achieved conditional control over RNA editing in response to diverse triggers, including small molecules, intracellular antigens, proteolytic activity, and light. When coupled with high-fidelity engineered substrates containing editable UAG stop codons, iDARs enable programmable control over protein translation and mRNA stability. By encoding these domains together with their substrates, we designed self-editing mRNA circuits capable of undergoing rapid induction and delivery as IVT mRNAs, thereby establishing a framework for fast and transient post-transcriptional control with potential applications in synthetic biology, cell engineering, and biological investigations.

A central contribution of this work is the demonstration that ADAR2-DD tolerates diverse regulatory insertions while remaining active, with the incorporation of *cis*-interacting domains providing a general mechanism for achieving autoinhibition. Tests of multiple fusion orientations revealed that autoinhibition depends on the placement of one binding unit at the C-terminus. This requirement suggests that intramolecular binding constraints disrupt the orientation of ADAR2-DD’s native C-terminal residues, which line the IP_6_-binding pocket^34^ and are required for efficient editing activity^47^. Notably, this architecture proved compatible across multiple tested binding partner insertion sites, ADAR paralogs, and protein-peptide pairs, together underscoring the platform’s versatility.

A fundamental challenge in engineering allosteric enzymes is balancing repressibility with inducibility^66^. We developed a rational framework for tuning iDAR sensitivity, minimizing background editing, and optimizing circuit dynamic range through combined mutagenesis of the inserted autoinhibitory units and native ADAR2-DD residues involved in IP_6_ coordination. Weakening intramolecular peptide-protein interactions predictably increased ligand sensitivity but also elevated basal activity. This effect was counterbalanced by introducing IP_6_-pocket mutations, which served to favor autoinhibited iDAR conformations by destabilizing their active configurations. Together, this dual-mutagenesis strategy enabled construction of chemiDARs with basal activities approaching those of inactive (dADAR) controls while retaining robust inducibility, achieving dynamic ranges exceeding 300-fold in some contexts. Importantly, this approach proved transferable across designs, providing a general strategy to improve the usability of individual iDAR constructs. The results further suggest that modulation of cofactor-binding sites may represent a viable strategy for enhancing the engineerability of other cofactor-bound enzymes beyond that of ADAR2-DD.

iDARs exhibit several technical distinctions from existing transcriptional and post-transcriptional control strategies. Because actuation occurs in the cytosol after RNA processing, iDARs enable rapid gene activation, with detectable protein outputs emerging within 15–30 minutes of stimulus exposure, substantially faster than transcription-based sensors while also being directly compatible with IVT-based methods^2,6^. Compared to purely RNA-based regulatory approaches, iDARs represent a modular scaffold for developing novel sensors that can be readily outfitted with diverse protein-based binding agents^4^. In contrast to systems based on induced heterodimerization^26–29^, the single-polypeptide iDAR design eliminates concentration-and stoichiometry-dependent variability while maintaining low basal activity and achieving high editing yields. Together with the mutagenic tuning strategies demonstrated here, these features provide a general framework for engineering new iDAR sensors with predictable and favorable performance characteristics.

To ensure effective implementation and to facilitate the design of new iDAR sequences, several practical aspects should be considered. First, the compatibility of specific inputs with the target cellular context must be evaluated, such as cell-type–specific tolerances for certain chemiDAR inducers, as certain cell lines are known to possess intrinsic sensitivity to A1 and ABT-737^67^. Further, achieving effective activation requires careful consideration of relevant inducer concentrations, as factors like low antigen expression may impose barriers to achieving efficient antiDAR activation. Second, when designing novel iDARs, constructs should be benchmarked against catalytically dead (RBD-dADAR) and constitutively active (RBD-ADAR) controls to assess background and induced editing activity levels. Identifying tightly regulated and highly inducible enzymes may require screening distinct insertion geometries and ADAR2-DD mutations, and the extent of required optimization may vary depending on payload selection, expression modality, or delivery mechanism.

The modularity and performance of the iDAR platform suggest utility beyond simple gene regulation, providing a versatile scaffold for discovery- and design-driven applications in biology and engineering. Because iDARs operate post-transcriptionally, they are well-suited for probing translation, including spatially restricted translation in polarized cells and neurons^68^, where optiDARs could be used to activate translation within defined subcellular regions. The development of iDARs that respond to cell-specific or dynamic intracellular cues, including markers of cell-cycle or transient signaling states, could further enable precise and programmable modulation of specific cellular processes^69–72^. Beyond regulating transgene expression, iDAR domains should integrate readily with existing ADAR-based technologies, such as guide-directed and localization-dependent editing platforms, to provide a conditional control layer for achieving inducible or cell-identity-dependent RNA editing^18,73–75^.

In summary, this work establishes general frameworks for designing allosterically controllable RNA-editing enzymes and for implementing them in logic circuits based on UAG stop-codon editing. By enabling systems capable of sensing and responding to user-defined stimuli at the post-transcriptional level, the iDAR platform represents an important step toward the design of autonomous, programable, and fast-acting input–output relationships within cells.

## DATA AVAILABILITY

The datasets generated during and/or analyzed during the current study are available from the corresponding author upon reasonable request.

## ACKNOWLEDGEMENTS

We thank D. Abdel-Meguid for preliminary work on ADAR reporters and R. Keating for preliminary work on lysiDARs. We are grateful to J. Tran and C. Kuffner for helpful discussions and advice regarding protein engineering. Research reported in this publication was supported by the National Institute of General Medical Sciences (NIGMS) of the National Institutes of Health (NIH) under award number R35GM128859 (to J.T.N.). The content is solely the responsibility of the authors and does not necessarily represent the official views of the NIH. A.M.M. was the recipient of a National Science Foundation Graduate Research Fellowship. The funders had no role in study design, data collection and analysis, decision to publish or preparation of the manuscript.

## COMPETING INTERESTS

A.M.M. and J.T.N. are co-inventors on a patent application filed by the Trustees of Boston University related to the design of inducible adenosine deaminases and post-transcriptional circuits (U.S. Patent Application, 18/392,928). A.M.M. and J.T.N. are co-founders, hold stock and board positions in Motive Tx.

## CONTRIBUTIONS

All authors contributed to the design and execution of experiments, analysis of the results, and preparation and editing of the manuscript.

## Notes

### Competing Interest Statement

A.M.M. and J.T.N. are co-inventors on a patent application filed by the Trustees of Boston University related to this work (U.S. Patent Application 18/392,928) and are co-founders of Motive Tx.

