## Supplementary Information for "Engineered Allosteric RNA Editors Enable Compact, Stimulus-Responsive Post-Transcriptional Circuits"

**CONTENTS:**

I. Materials and Methods
II. Supplementary Figures 1-21
III. Supplementary Notes 1-2
IV. Additional References

### MATERIALS AND METHODS

#### DNA Constructs

DNA constructs were generated by Gibson assembly using the HiFi Assembly Master Mix (New England Biolabs, NEB) or by ligation of cleaved fragments using T4 DNA ligase (New England Biolabs). DNA fragments were generated via polymerase chain reaction (PCR) amplification using Phusion High-Fidelity PCR Master Mix (NEB) or ordered as custom synthesized sequences via Integrated DNA Technologies (IDT), Twist Biosciences, or Azenta. All construct sequences were verified by Sanger sequencing before use (Quintara Biosciences).

The MS2_adRNA-MCP_ADAR2 DD (E488Q)_NES plasmid was a gift from Prashant Mali (Addgene plasmid # 124707), pDY1173_ADAR1p150 was a gift from Omar Abudayyeh and Jonathan Gootenberg (Addgene plasmid # 193192), and the ADARB1-HiBiT Fusion Vector was a gift from Promega Corporation (Addgene plasmid # 236892).

Sequence information for the utilized genetic parts and generated reporter constructs, along with details regarding their experimental use, are available upon request and will be published upon acceptance of the manuscript as **Supplementary Tables 1-5**.

#### Mammalian Cell Culture

HEK293FT cells (Thermo Fisher, R70007) were maintained in a humidified incubator at 37° C with 5% $CO_2$. Cells were cultured in media based on high-glucose Dulbecco's Modified Eagle Medium (DMEM; Cytiva, SH30285) supplemented with 10% (v/v) fetal bovine serum (FBS; typically, either Cytiva SH30396.03 or Corning 35010CV), 1x nonessential amino acids, 1x Glutamax (LifeTechnologies), and 1x penicillin-streptomycin solution. For experiments involving cells containing doxycycline-inducible reporter genes, media containing 10% (v/v) Tet-System Approved FBS (Takara, 631106) was used in place of regular FBS.

#### DNA Transfections

DNA transfections were carried out using Lipofectamine 3000 Reagent (Thermo Fisher, L3000015) according to the manufacturer's instructions. Unless otherwise noted, cells were transfected in 96-well plates using 75,000 freshly trypsinized HEK293FT cells suspended in 200 µL of growth media per well.

Transfection mixtures were prepared in Opti-MEM (Thermo Fisher, 31985062) using a total of 100 ng of DNA per well; when necessary, DNA mixtures were supplemented with empty vector or salmon sperm DNA as "filler" prior to transfection complex preparation. The amounts of specific DNA constructs used were varied according to individual experimental applications as described in the sections below.

***Dual transfection of ADAR-ON reporters and iDAR/ADAR variants:*** Transfection complexes were prepared using 30 ng of ADAR-ON reporter plasmid in combination with 20 ng of plasmid encoding the RBD-iDAR/ADAR variants or their corresponding inactive dADAR versions as specified in the figure captions. For ADAR-lacking controls, 20 ng of empty vector (pcDNA3) was used in place of an ADAR- or dADAR-encoding plasmid. Individual DNA mixtures were supplemented with 50 ng of empty vector (pcDNA3) as filler. For deaminase titration experiments, 10 ng of ADAR-ON reporter plasmid was combined with varying amounts of MCP-ADAR-encoding plasmid (0.01-100 ng, as indicated in the captions) and supplemented with a corresponding amount of empty vector to reach a total DNA amount of 110 ng per well.

***Dual transfection of ADAR-OFF reporters and iDAR/ADAR variants:*** GFPd2-encoding ADAR-OFF reporter constructs were encoded together with a separate dTomato-encoding transcription using a plasmid containing a bidirectional CMV promoter (originally based on pGuide-it-tdTomato, Takara, 632604). Transfection mixtures were prepared using 10 ng of the indicated dTomato/ADAR-OFF reporter variants, 25 ng of plasmid encoding the specified RBD-ADAR constructs, and 65 ng of empty vector (pcDNA3) as filler DNA. For deaminase titration experiments, 10 ng of dTomato/ADAR-OFF reporter plasmid was combined with varying amounts of MCP-ADAR-encoding plasmid (0.01-100 ng, as indicated in the captions) and supplemented with empty vector to a final total DNA concentration of 110 ng per well.

***Triple transfection of ADAR-ON reporters, protein-responsive iDARs (antiDARs), and antigen-expressing vectors:*** Transfection complexes were prepared using 30 ng of plasmid encoding the specified ADAR-ON reporter, 20 ng of plasmid encoding the indicated MCP-antiDAR-tagBFP variants, 10 ng of plasmid encoding the specified antigen proteins and 40 ng of empty vector as filler DNA.

***Triple transfection of ADAR-ON reporters, protease-responsive iDARs (lysiDARs), and protease-expressing vectors:*** Transfection complexes were prepared using 30 ng ADAR-ON reporter plasmid, 20 ng of plasmid encoding the indicated MCP-lysiDAR-tagBFP variants, 3 ng of a pcDNA3-based plasmid encoding TEVp and 47 ng of empty pcDNA3 as filler DNA.

***Single and dual transfection of self-editing iDAR circuits:*** Transfection complexes were prepared using 100 ng of plasmid encoding the self-editing iDAR circuits indicated in the figure captions. For endpoint NLuc measurements, transfection mixtures were supplemented with an additional 10 ng of Firefly luciferase (Fluc)-encoding plasmid (pcDNA3-Fluc) for luminescence normalization.

For antiDAR-based circuits, mixtures contained an additional 10 ng of plasmid encoding the specified antigens. For lysiDAR-based circuits, mixtures contained an either an additional 3 ng of plasmid encoding full-length TEVp, or an additional 30 ng of DNA mixture containing plasmids encoding rapamycin-inducible split-TEVp components.

For titration experiments where antigen or protease levels were varied, transfection mixtures were prepared using 100 ng of plasmid encoding the specified self-editing iDAR circuits and varying amounts of antigen- or protease-encoding plasmid (0.01-100 and 0.001-30 ng respectively, as indicated in the captions). Plasmid mixtures were supplemented with empty vector to reach a total amount of 200 ng per transfection.

***Preparation of cells for mRNA imaging by single-molecule FISH:*** Transfection complexes were prepared using 1 ng of dTomato/ADAR-OFF reporter plasmid, 20 ng of plasmid encoding the indicated MCP-ADAR2-DD-tagBFP variants and 54 ng of empty vector as filler DNA. Transfections were carried out in 18-well chambered cover glass plates (Cellvis, C18SB-1.5H) precoated with fibronectin using a solution of 10 µg/mL fibronectin in phosphate buffered saline (PBS) for 1 hour at room temperature.

Transfections were initiated by combining the DNA complexes with 50,000 HEK293FT cells suspended in growth media followed by gentle mixing before seeding into individual microwells.

**Flow Cytometry and Gating Procedures**

Flow cytometry was performed using an Attune-NxT Flow Cytometer (Thermo Fisher). Transfected cells were analyzed at approximately 48 hours post-transfection unless otherwise indicated. Cells cultured in 96-well plates were dissociated using 50 µL of pre-warmed 0.25% Trypsin-EDTA per well with incubation at 37° C for ≤3 min. Trypsinization reactions were quenched by adding 200 µL of pre-warmed growth medium; the generated cell suspensions were then utilized for flow cytometry. Data were analyzed using the FlowJo v10 software.

Representative gating procedures for discriminating relevant cell populations are provided in **Supplementary Fig. 1**. Live cells were gated based on forward and side scatter (FSC-A vs. SSC-A) and singlets were determined by side scatter (FSC-A vs. FSC-H). For ADAR-ON/OFF reporter assays, cells were gated based on the top 0.5% of non-transfected HEK293FT controls as analyzed under mCherry/dTomato detection settings. For tagBFP-fused iDAR constructs, cells were gated based on the top 1% of non-transfected HEK293FT cells as analyzed under tagBFP detection settings. mNG positive cells were defined as those displaying fluorescent intensities exceeding the top 0.5% of non-transfected HEK293FT controls. Co-transfected cells were identified on the basis of dual mCherry/dTomato (reporter) and tagBFP (iDAR) emissions. For self-editing circuits encoding iDAR-mCherry fusions, transfected populations were gated based on the top 0.5% of non-transfected HEK293FT as analyzed under mCherry/dTomato detection settings. For experiments involving *TRE3G-mTurquoise2* reporter cells, mTurquoise2-positive cells were gated based on the top 1% of non-transfected reporter cells, as analyzed under mTurquoise2 detection settings.

mNG/mCherry and GFPd2/dTomato fluorescence ratios were calculated using measurements collected from cells dually positive for mCherry/dTomato and tagBFP. Binning analyses (**Supplementary Fig. 1b**) indicated that reporter-specific background signal was distinguishable from basal cellular fluorescence only at higher expression levels (>$10^4$ AFU) and were more indicative of true fold-changes. Consequently, quantitative analyses were restricted to this high-expression population to ensure that editing-dependent signal could be accurately distinguished from cellular autofluorescence and noise.

**Fluorescence *in situ* hybridization (FISH) labeling and analysis**

FISH labeling was carried out using a previously described protocol based on the Hybridization Chain Reaction (HCR)[1] utilizing buffers and reagents from  HCR RNA-FISH Bundle (Molecular Instruments). Briefly, transfected HEK293FT cells were prepared in triplicate as described above. At 24 h post-transfection, cells were fixed using a 4% formaldehyde solution (w/v) diluted in PBS from a 16% (w/v) methanol-free formaldehyde stock (Thermo Fisher, 28906). The fixation reaction proceeded for 10 min at room temperature prior to rinsing with PBS. Cells were permeabilized using prechilled 70% (v/v) ethanol in water solution with treatment at -20°C overnight. Next day, hybridization was performed using the split B1-FISH probe set targeting '*d2eGFP'* (Molecular Instruments) followed by amplification using B1-AlexaFluor647 amplifier probes (Molecular Instruments). Labeled cells were washed according to the manufacturer's protocol and subsequently imaged as submerged in PBS containing 0.5% Tween-20 (v/v).

Widefield images were taken with a Zeiss Axio Observer Z1 microscope equipped with an HXP 120-V halogen lamp as the excitation source. Images were recorded using a Prime95B sCMOS camera (Teledyne Photometrics) and analyzed in ImageJ/FIJI software. Images were captured through a 20x/0.8 numerical aperture air objective lens. GFPd2 was visualized under 'eGFP' filter settings (Zeiss Filter Set 38; BP 470/40, FT 495, BP 525/50); dTomato under 'Cy3' filter settings (Zeiss Filter Set 43; BP 550/25, FT 570, BP 605/70); ADAR-tagBFP under 'EBFP2' filter settings (Chroma 49021; ET405/20x, T425lpxr, ET460/50m); and AlexaFluor647-labeled single-mRNAs under 'Cy5 Narrow Excitation' filter settings (Chroma 49009; ET640/30x, T660lpxr, ET690/50m). To minimize bias during data recording, captured well regions and Z-planes were selected based on solely on dTomato emissions.

Raw image files (.czi) were processed and quantified using a custom macro in ImageJ/Fiji. To correct for background fluorescence, the modal gray value of each channel was calculated and globally subtracted (this value was confirmed to approximate the background signal measured in non-transfected cell regions). To identify cells co-expressing both the reporter and the ADAR construct, binary masks were generated for dTomato (reporter) and tagBFP (ADAR) channels using empirical intensity thresholds. A final segmentation mask was created via the Boolean intersection (AND operation) of the dTomato and tagBFP masks, and individual cellular Regions of Interest (ROIs) were defined using the 'Analyze Particles' function using a threshold of > 20 pixels). For ratiometric quantification, 32-bit floating-point ratio images were generated by pixel-wise division of signal channel values (for GFP or AlexaFluor647) by corresponding normalization channel (dTomato) intensities. The median pixel intensity of the resulting ratio images was measured within each cellular ROI. Calculated values were exported to GraphPad Prism for statistical analysis. Outliers were identified using the ROUT (Q = 1%) method in Prism; any ROI identified as an outlier in either the GFP/dTomato or AlexaFluor647/dTomato datasets were eliminated from datasets to ensure consistent analyses. The custom macro script is provided in **Supplementary Note 1**.

#### Drug compounds and chemiDAR activation

The following small molecule ligands were obtained from MedChemExpress: ABT-737 (HY-50907), A-1331852 ('A1'; HY-19741), S63845 (HY-100741), and grazoprevir (HY-15298). Stocks of each compound were prepared by dissolution in dimethyl sulfoxide (DMSO) at 5-10 mM concentration and stored as aliquots at -20° C. For chemiDAR activation, ligand stocks were diluted into pre-warmed growth media and added to microwells immediately after seeding transfection/cell suspension mixture. For real-time kinetic assays, ligand-containing media were added to wells following baseline signal equilibration under ligand-free conditions (see section describing luminescence measurements). Unless otherwise indicated in the figure legends, A1, S63845 and grazoprevir were used at a final concentration of 1 µM. Vehicle control samples were prepared by using volume-matched amounts of DMSO. For dose-response experiments, serial dilutions were prepared using fresh growth media and matched final DMSO concentrations. Care was taken to ensure that treatment media contained DMSO at 0.1% (v/v) or less for all conditions.

#### Generation and maintenance of stable cell lines

A clonal HEK293FT reporter line bearing an integrated *TRE3G-mTurquoise2* gene construct was isolated by limiting dilution. Cells were maintained under selection with 0.5 µg/mL puromycin and were transferred to puromycin-lacking growth media prior to transfection.

To generate inducible antigen-expressing cell lines, a sequence encoding EGFP(R96M)-AlfaTag-p24CTD was cloned into a doxycycline-inducible lentiviral transfer vector in-frame with a downstream sequence encoding T2A-TagBFP as a fluorescence marker (**Supplementary Table 5)**. Lentiviral particles were produced using a second-generation system by co-transfecting HEK293FT cells with the transfer plasmid in combination with packaging (psPAX2) and envelope (pVSV-G) plasmids using Lipofectamine 3000. Cells were grown in 6-well TC treated plates to 90% confluence prior to transfection with 750 ng of transfer plasmid, 1200 ng each of psPAX2 and pVSV-G plasmids. Transfection media were replaced at 6 h post-transfection. Viral supernatants were harvested at 48 h post-transfection and cell debris was eliminated by filtration through sterile PES syringe filters containing 0.45 µm pores. Filtered supernatants were used to transduce HEK293FT cells immediately or stored as aliquots at -80° C. At 48 h post-transduction, cells were subjected to selection in growth media containing Hygromycin B at 75 µg/mL. Stably transduced pools were maintained for 4 passages (~10 days) with regular media replenishment. Induced expression of the encoded constructs was confirmed via tagBFP detection following 24 h of growth in media containing doxycycline at 1 µg/mL.

**Luminescence assays**

Endpoint luminescence measurements for self-editing plasmid experiments were performed using the Nano-Glo Dual-Luciferase Reporter Assay System (Promega, E1980) according to the manufacturer's instructions. Briefly, cells in white 96-well plates (Corning, 353296) were analyzed at 48 h post-transfection by exchange into pre-warmed Opti-MEM using 40 µL per well. FLuc activities were measured by adding 40 µL of the ONE-Glo EX Reagent to each well followed by incubation at room temperature for 5 min with gentle orbital shaking. FLuc luminescence levels were then recorded on a Cytation 5 Cell Imaging Multimode Plate Reader (BioTek) using a 1 sec integration time. FLuc activity was quenched prior to measuring NanoLuc (NLuc) activities by adding 40 µL of the NanoDLR Stop & Go Reagent. Samples were incubated for 5 min before recording NLuc luminescence values using a 1 s integration time. Reported values represent normalized NLuc/FLuc ratios.

For real-time kinetic measurements, cells were transfected in HEPES buffered Opti-MEM containing 5% (v/v) FBS. At 24 h post-transfection, cells were exchanged into 225 µL of pre-warmed Opti-MEM containing a 1:100 dilution of Nano-Glo Vivazine Live Cell Substrate (Promega, N2590). Plates were equilibrated to 37° C under 5% $CO_2$ within the Cytation 5 instrument until signal stabilization was reached (~1.5 h), at which point cells were treated with either DMSO (control) or A-1 at a final concentration of 1 µM. Treatments were carried out by adding 25 µL Opti-MEM containing DMSO or A-1 at 10x working concentration for final volumes of 250 µL per well. Luminescence levels were then monitored continuously for 16 h. Measured values were normalized to the mean signal of constitutively active MCP-ADAR controls (n=3) at each time point to get the fraction of maximal activity.

For cells transfected with IVT mRNAs, NLuc luminescence values were recorded ~24 h post-transfection by exchanging cells into 1x Universal Lysis Buffer (NanoLight Technology, 333-50; diluted from 5x stock in PBS) supplemented with 10 µM fluorofurimazine (Selleck Chemicals, E1620) as the luminescent substrate. The substrate-containing lysis buffer was added to cells at a volume of 100 µL per well. Lysis proceeded for 5 min at room temperature before transfer of lysates into a luminescence compatible white 96-well microplate. Luminescence values were then recorded via the Cytation 5 plate reader using a 1 s integration time. Aliquots of the fluorofurimazine substrate were stored at -20°C as 10 mM stocks in DMSO.

**OptiDAR photoactivation, live-cell microscopy and analysis**

HEK293FT cells were transfected as described above in fibronectin-coated, glass-bottom, black-walled 96-well plates (Cellvis, P96-1-N). At 24h post-transfection, cells were exchanged into pre-warmed HEPES-buffered Opti-MEM supplemented with 10% (v/v) FBS. Cells were then imaged and subjected to on-microscope PhoCl activation using a Zeiss AxioObserver inverted widefield epifluorescence microscope equipped with an HXP 120V metal-halide lamp as the illumination source. Cells were kept in a humidified environmental chamber maintained at 37° C during photoactivation and subsequent imaging.

For imaging and photoactivation, a 3×3 tile region (approx. 2 mm × 2 mm) was defined for each well based on mCherry$^+$ transfection density. The experiment consisted of iterative cycles of image acquisition (mCherry and mNG channels) followed by photoactivation for the subset of 'Illuminated' cells. Images were captured through a 20x/0.8 numerical aperture air objective lens. mNG emissions were recorded under 'phYFP' filter settings (Chroma 49003; ET500/×20, T515lp, ET535/30m) and mCherry emission under 'mCherry' filter settings (Chroma 49008; ET560/×40, T585lpxr, ET630/75m). Exposure times were set to optimize signal dynamic range of fluorescence accumulation over the non-steady-state time course.

Photocleavage of PhoCl was carried out using 10 sec illumination per field of view (9 per well) using light from an HXP 120V lamp as applied to cells through an ET405/20x excitation filter present within an 'EBFP2' filter set (Chroma 49021; ET405/20x, T425lpxr, ET460/50m). These conditions were empirically verified to facilitate efficient PhoCl photocleavage, as determined based on immediate green-to-red conversion in singly transfected reference cells expressing only the PhoCl-containing optiDAR (without a reporter sequence). The experimental protocol consisted of alternating imaging and activation steps: samples were imaged at t = 0 h, followed by photoactivation at t =1.5 h later. Subsequent rounds of imaging and photoactivation occurred at 3 h intervals (imaging at t = 3, 6, 9, 12, 15, and 18 h; activation at t = 4.5, 7.5, 10.5, 13.5, and 16.5 h) for a total of 6 cycles lasting 18 h in duration. Post-acquisition, full-well montages were acquired to map spatial activation patterns within each well.

Raw tiled images from the time-lapse experiment were stitched using the ZEN Black Edition software (Zeiss). Stitched images were then processed in ImageJ/Fiji using a custom macro. Background fluorescence was corrected by global subtraction of the modal gray value of each channel (this value was confirmed to approximate the background signal measured in non-transfected cell regions). Transfected cells were segmented using a binary mask generated based on mCherry emissions with an empirical threshold set to >50 AFU. Masks were further refined via hole filling and watershed segmentation to separate adjacent cells, followed by conversion into Regions of Interest (ROIs) using the 'Analyze Particles' function (particle size > 100 pixels; circularity 0.1–1.0). For ratiometric quantification, 32-bit floating-point ratio images were generated by pixel-wise division of reporter mNG intensities by the normalization signal (mCherry) for each time point. Median pixel intensities from ratio images were then measured within each cellular ROI. Data were exported to Microsoft Excel and means for each condition and timepoint were determined, and values were then statistically analyzed in GraphPad Prism. The custom macro script is provided in **Supplementary Note 2**.

#### *in vitro* transcription (IVT) and mRNA Transfection

To create templates for IVT, custom DNA synthesis was used to generate a plasmid bearing the human *β-globin* 5' UTR and *AES/mtRNR1* 3' UTR (similar to those utilized in the BNT162b2 vaccine template) followed by a unique *PmeI* site for linearization and polyA addition. PCR-generated fragments corresponding to tdMCP-iDAR-mCherry-UAG$_{4x}$-NLuc circuits were inserted into this custom vector via Gibson assembly. The resulting vector was verified by Sanger sequencing and utilized as a template for IVT.

Prior to transcription, plasmid templates were linearized by restriction digests with PmeI and MfeI enzymes (NEB). Overdigestions were performed by incubation at 37° C for 2 hours to ensure complete template linearization. Linearized DNAs were purified using the DNA Clean & Concentrator-5 kit (Zymo Research, D4014) and eluted using RNAse Free Water (Invitrogen, 10977015). DNA concentrations were determined by extinction using a NanoDrop One spectrophotometer (Thermo Fisher).

IVT reactions were carried out using the HiScribe T7 ARCA mRNA Kit (with tailing; NEB, E2060). Capped, non-modified mRNAs were synthesized using Anti-Reverse Cap Analog (ARCA) incorporation, followed by enzymatic polyadenylation using *E. coli* Poly(A) Polymerase according to the HiScribe T7 ARCA mRNA Kit protocol. Synthesized mRNAs were purified using the Monarch RNA Cleanup Kit (50 µg scale; NEB, T2040L) and eluted in RNA Storage Solution (Thermo Fisher, AM7001). RNA concentrations were determined by extinction coefficient using a NanoDrop One spectrophotometer. Generated mRNAs were stored as aliquots at -20° C for short-term use (<1 week) and kept at -80° C for long-term storage.

HEK293FT cells were transfected with purified mRNAs using Lipofectamine MessengerMAX (Thermo Fisher, LMRNA008) according to the manufacturer's suspension transfection protocol. Typically, 100 ng of iDAR mRNA was used to transfect 75,000 HEK293FT cells per well at the 96-well plate scale. Transfected cells were assayed for NLuc expression by luminescent detection at 24 h post-transfection (see corresponding section of luminescence assays).

#### Data collection and analysis software

Flow cytometry data were acquired using the Attune NxT Software (Thermo Fisher) and analyzed using FlowJo v10 (BD Life Sciences). Microscopy images were captured using ZEN Black Edition 2.3 (Zeiss) and analyzed using ImageJ/Fiji v2.16.0. Luminescence measurements were recorded using the Gen5 Microplate Reader and Imager Software (BioTek). Statistical analyses and graphing were performed using GraphPad Prism v10.4.2.

#### Statistics and reproducibility

Statistical analyses were performed using GraphPad Prism (version 10.4.2). Unless otherwise noted, data are presented as mean ± standard deviation (s.d.) of n=3 independently transfected samples processed in parallel. Each sample analyzed by flow cytometry contained over 1000 cells after gating. Unless otherwise noted, the median values derived from flow cytometry samples were used. All microscopy images are representative of results from at least three independent replicates. Specific statistical tests performed are described in the corresponding figure legends.

### SUPPLEMENTAL FIGURES

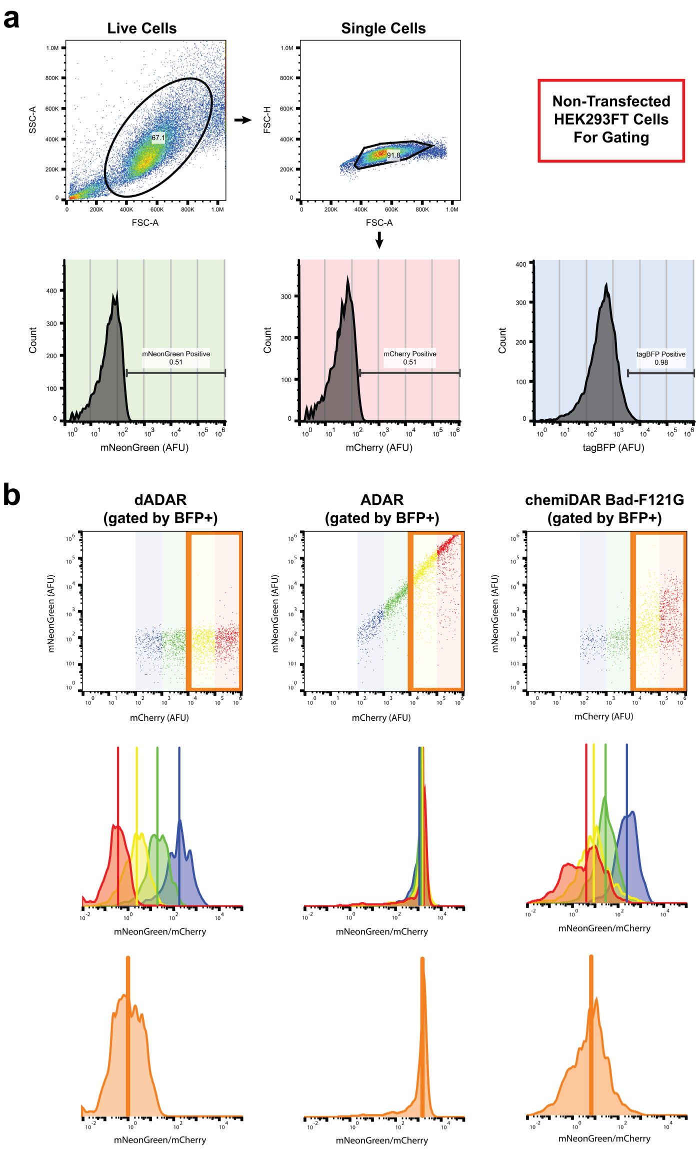


***Supplementary Figure 1 – Flow cytometry gating strategy and quantification of reporter translation efficiency***

**a**, Representative gating workflow using non-transfected HEK293FT cells as negative controls. Live cells were selected based on FSC-A vs SSC-A, followed by singlet gating on FSC-A vs FSC-H. Fluorescent protein-positive gates were defined relative to background fluorescence of non-transfected HEK293FT cells: the top 0.5% for mCherry and mNeonGreen (mNG), and the top 1% for tagBFP. For co-transfection experiments, tagBFP$^+$ cells were used to identify transfected populations, whereas for self-editing/all-in-one constructs, mCherry$^+$ cells were used. Percent mNG$^+$ cells were determined relative to the mNG fluorescence gate.

**b,** Representative analysis of how mNG/mCherry ratios were determined. HEK293FT cells were co-transfected with an ADAR-responsive reporter and MCP-ADAR-tagBFP construct, then gated on tagBFP$^+$ cells. (Top) Scatter plots of mNG vs mCherry expression, with mCherry fluorescence subdivided into logarithmic bins: $10^2$–$10^3$ (blue), $10^3$–$10^4$ (green), $10^4$–$10^5$ (yellow), and $10^5$–$10^6$ (red). The high-expression region (orange box, $10^4$–$10^6$ AFU) was selected for downstream analysis. (Middle) Histograms showing the distribution of mNG/mCherry ratios within each bin; vertical lines mark the median of each distribution. (Bottom) Pooled histogram of high-expressing cells ($10^4$–$10^6$ AFU), with the median indicated by a vertical line. This gating strategy permitted clearer discrimination of background activation and ADAR-dependent reporter signal.

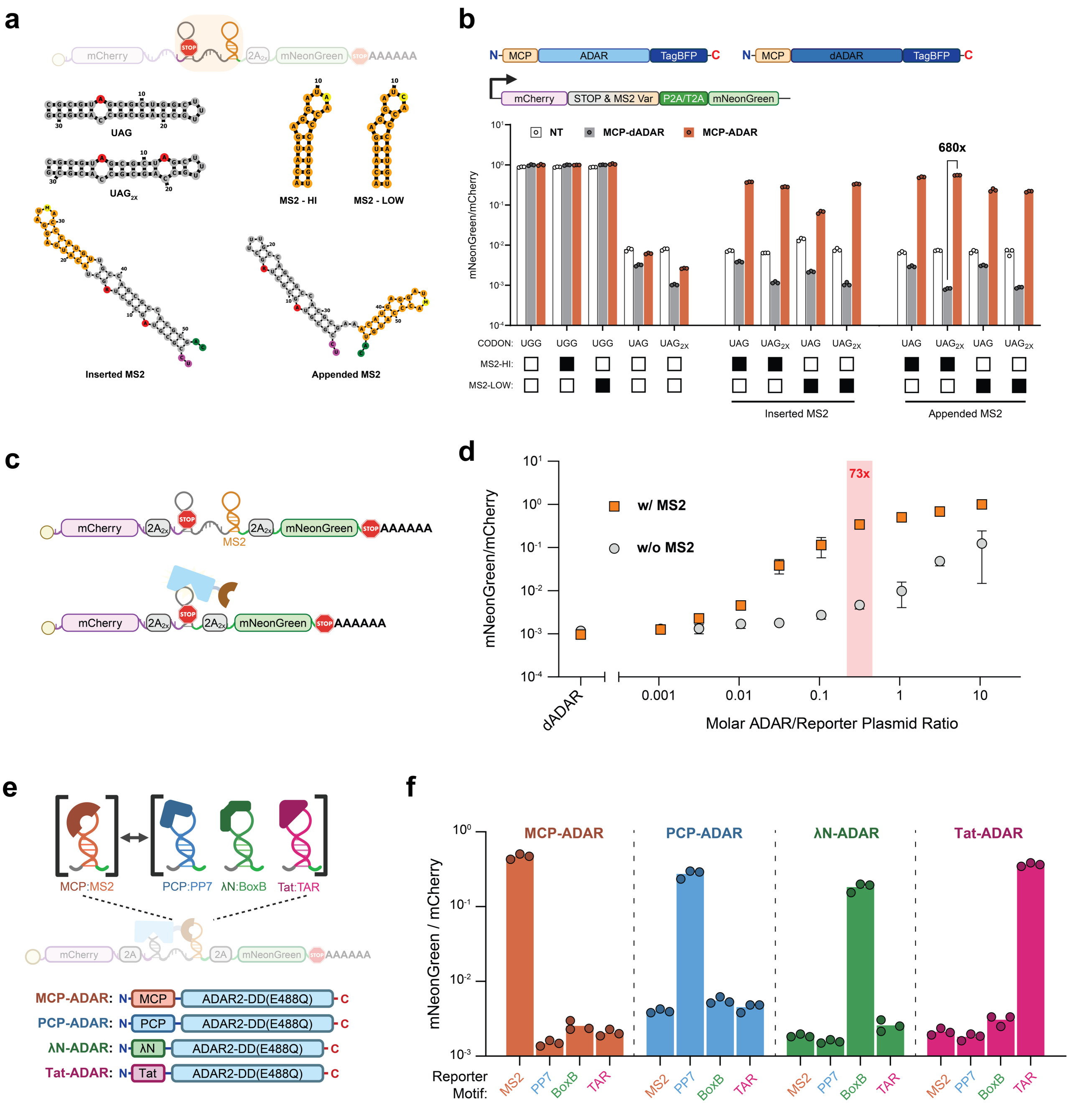


***Supplementary Figure 2 – Optimization of reporter design and assessment of RNA motif specificity***

**a**, Schematic of reporter RNA substrates (gray) showing variant ADAR target sites and RNA motif positioning (orange). Predicted secondary structures of substrates containing one (top) or two (bottom) UAG stop codons are shown, with editable adenosines in red. Two MS2 operator variants with substitutions at the -5 position (yellow) are shown: MS2-HI (C variant, high affinity, $K_D$ ~1 nM) and MS2-LOW (A variant, lower affinity, $K_D$ ~400-600 nM). The relative positioning of MS2 motifs (Inserted vs Appended) and substrates is shown. RNA secondary structures were predicted by RNAfold[2].

**b**, Construct maps of active MCP-ADAR and inactive MCP-dADAR(E396A) together with the reporter variants used in this assay. HEK293FT cells co-transfected with the indicated reporters and either control DNA (white), MCP-dADAR (gray), or MCP-ADAR (orange). Median mNeonGreen/mCherry fluorescence ratios were quantified 48 h post-transfection by flow cytometry. Bars, mean; points, *n* = 3 independent transfections.

**c**, Illustration of cis-acting on-target editing (top), in which MCP-ADAR edits UAG codons within MS2-containing reporters, compared with non-specific editing (bottom) that may occur in the absence of an RNA motif. This comparison serves as a proxy for off-target activity.

**d**, Dose-response of MCP-ADAR activity. HEK293FT cells were co-transfected with 10 ng of reporter RNA (with or without MS2 motifs) and increasing amounts of MCP-ADAR plasmid. MCP-dADAR served as a negative control. Cells were analyzed 48 h post-transfection by flow cytometry. The optimal ADAR:reporter DNA ratio producing maximal signal discrimination is highlighted in red. Data represent median mNeonGreen/mCherry fluorescence; bars, mean; points, n = 3 independent transfections.

**e**, Schematic of orthogonal RNA-binding protein (RBP) and RNA motif pairs tested: MCP/MS2 (orange), PCP/PP7 (blue), λN/BoxB (green), and Tat/TAR (pink).

**f**, Specificity of RBP-motif interactions. HEK293FT cells were co-transfected with ADAR-ON reporters containing one of the four RNA motifs and ADAR constructs fused to the corresponding or mismatched RBDs. Cells were analyzed by flow cytometry 48 h post-transfection. Bars represent median mNeonGreen/mCherry fluorescence ratios per replicate, normalized to a positive control reporter containing UGG sense codons in place of editable UAGs.

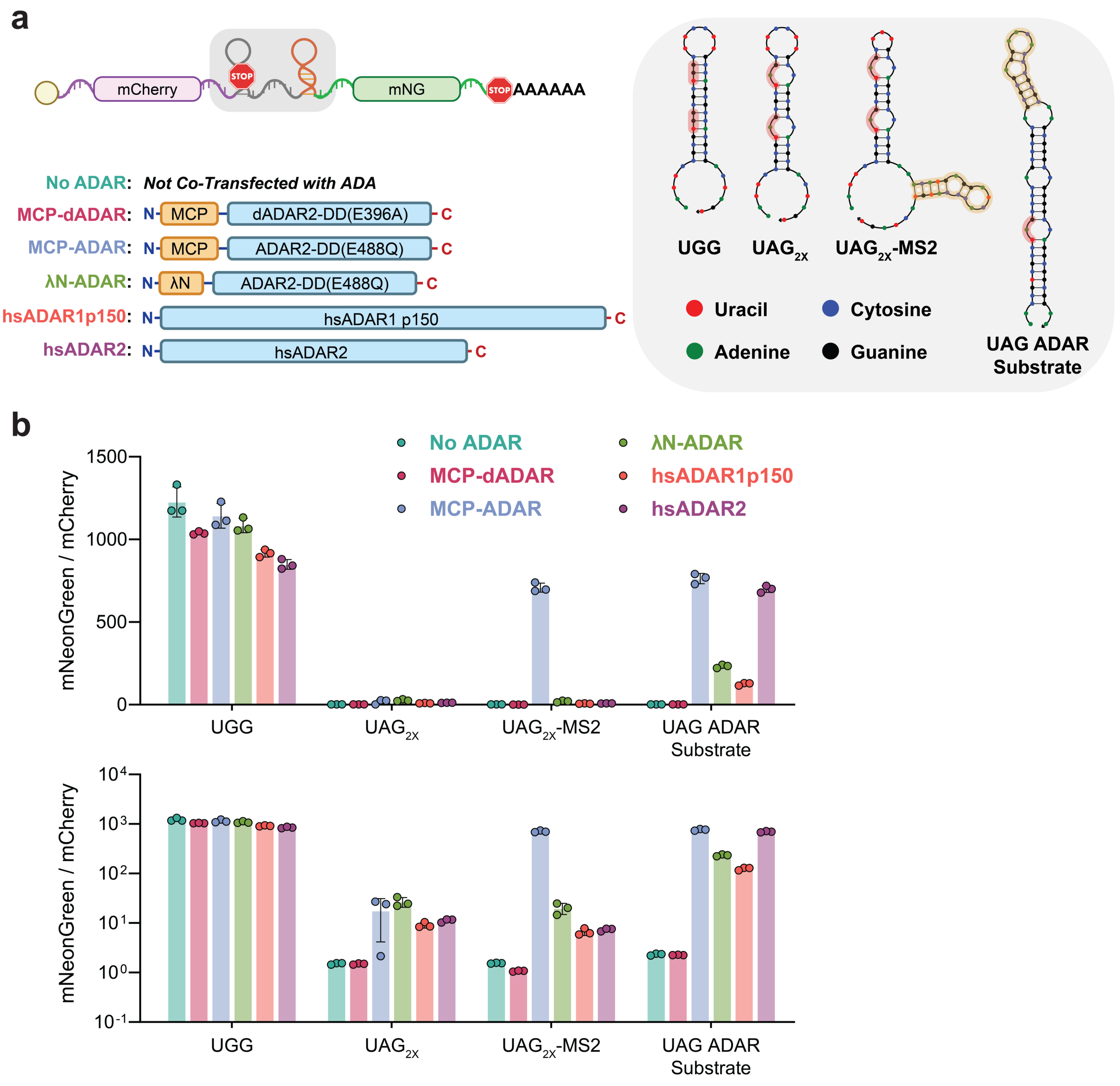


***Supplementary Figure 3 – ADAR-ON reporters are inefficient substrates for full-Length human ADAR1-p150 and ADAR2***

**a**, Schematic of reporter and ADAR-DD containing constructs used to assess off-target editing by recombinantly overexpressed full-length human ADAR1-p150 (hsADAR1) and ADAR2 (hsADAR2) proteins. Reporters contained the previously tested UGG-containing control sequence, editable substrates based on $UAG_{2x}$ or $UAG_{2x}$-MS2, as well as an hsADAR editing positive-control substrate based on a sequence predicted to be efficiently edited by full-length hsADAR1 and hsADAR2 (UAG ADAR Substrate). Predicted RNA secondary structures were generated by NUPACK[3] (right).

**b**, Activation of the reporter constructs when co-transfected with various ADAR-DD containing proteins. HEK293FT cells co-transfected with the indicated reporters and either salmon sperm DNA ('No ADAR' – seafoam green), MCP-dADAR (red), MCP-ADAR (blue), λN-ADAR (green), full-length hsADAR1-p150 (orange) or hsADAR2 (purple). Median mNeonGreen/mCherry fluorescence ratios were quantified 48 h post-transfection by flow cytometry. Top and bottom represent the same data shown on a linear (top) or log (bottom) scale. Bars, mean; error-bars, SD; points, *n* = 3 independent transfections.

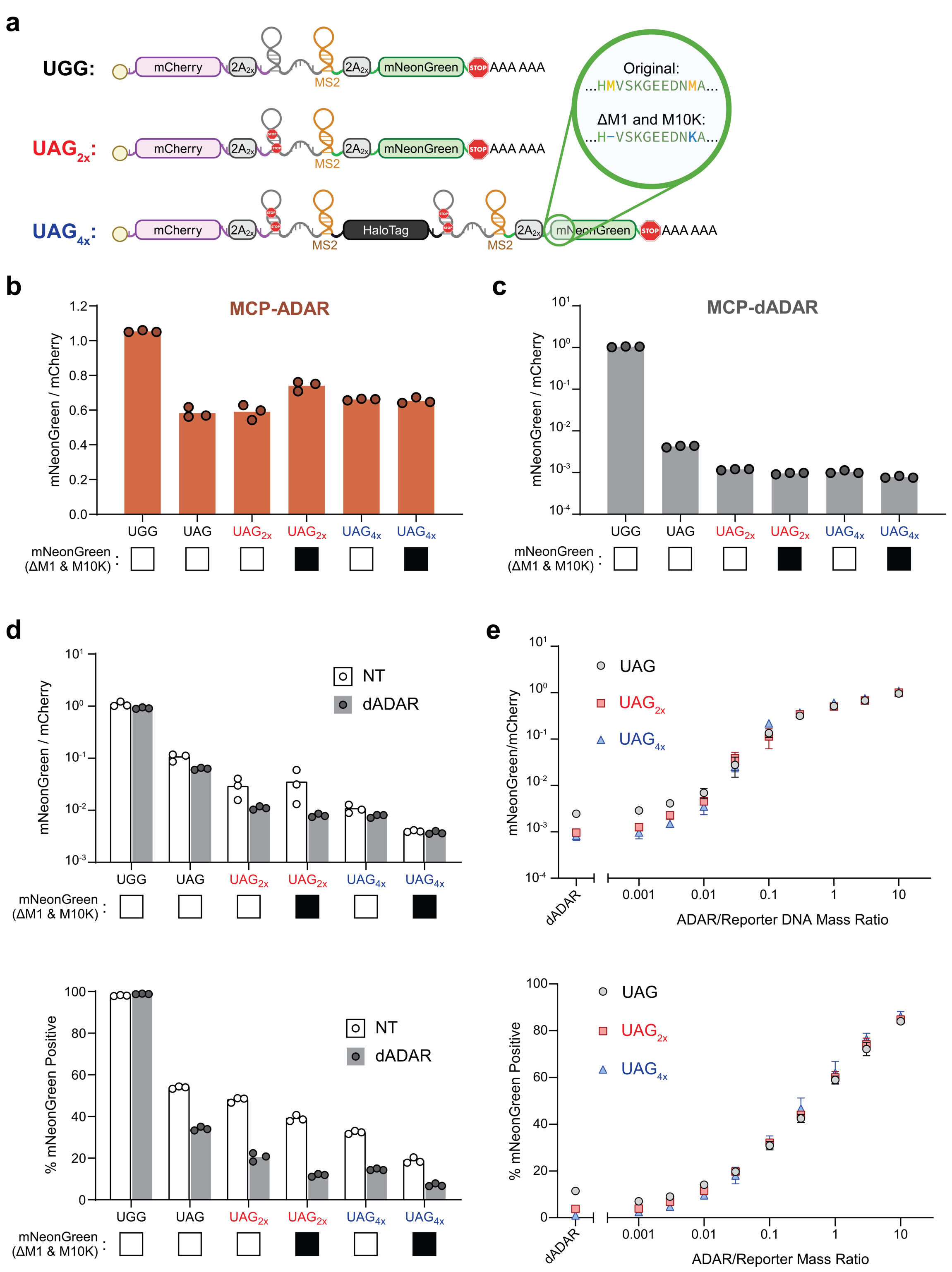


***Supplementary Figure 4 – Increasing UAG redundancy and removing internal methionines reduces background and maintains ADAR-dependent activation***

**a**, Reporter designs used to evaluate strategies for minimizing background readthrough translation. Reporters contained one, two, or four editable and in-frame UAG codons (UAG, $UAG_{2x}$, $UAG_{4x}$) with or without deletion of the first methionine (ΔM1) codon and substitution of a downstream in-frame methionine codon with a lysine codon (M10K) within the mNeonGreen (mNG) coding sequence.

**b**, Reporter activation in the presence of catalytically active MCP-ADAR. Median mNeonGreen/mCherry fluorescence ratios are shown for each construct. Bars, mean; points, *n* = 3 independent transfections.
**c,** Reporter background when co-transfected with catalytically inactive MCP-dADAR. Bars, mean; points, *n* = 3 independent transfections.
**d,** Comparison of background expression in non-transfected (NT, white) and MCP-dADAR co-transfected (gray) HEK293FT cells. Reporters containing four stop codons and lacking internal methionine codons ($UAG_{4X}$, ΔM1/M10K) exhibited the lowest background activity. Cells were analyzed 48 h post-transfection by flow cytometry. Detector voltages were adjusted to detect low-level background, resulting in saturation of the UGG (control) signal. Top: median mNeonGreen/mCherry fluorescence per replicate, normalized to a stop-codon-free reporter control. Bottom: percentage of $mNG^+$ cells per condition, defined relative to the top 0.5 % fluorescence in non-transfected HEK293FT controls. Bars, mean; points, *n* = 3 independent transfections.
**e,** Dose-response of ADAR-dependent reporter activation. HEK293FT cells were co-transfected with 10 ng of reporter plasmid (UAG, $UAG_{2x}$, $UAG_{4x}$) and increasing amounts of MCP-ADAR plasmid. Cells were analyzed by flow cytometry 48 h post-transfection, and activation was quantified as median mNeonGreen/mCherry ratio (top) or percent $mNG^+$ cells (bottom). Positive cells were defined as those exceeding the top 0.5 % fluorescence in non-transfected controls. Points, mean; error bars, SD; *n* = 3 independent transfections.

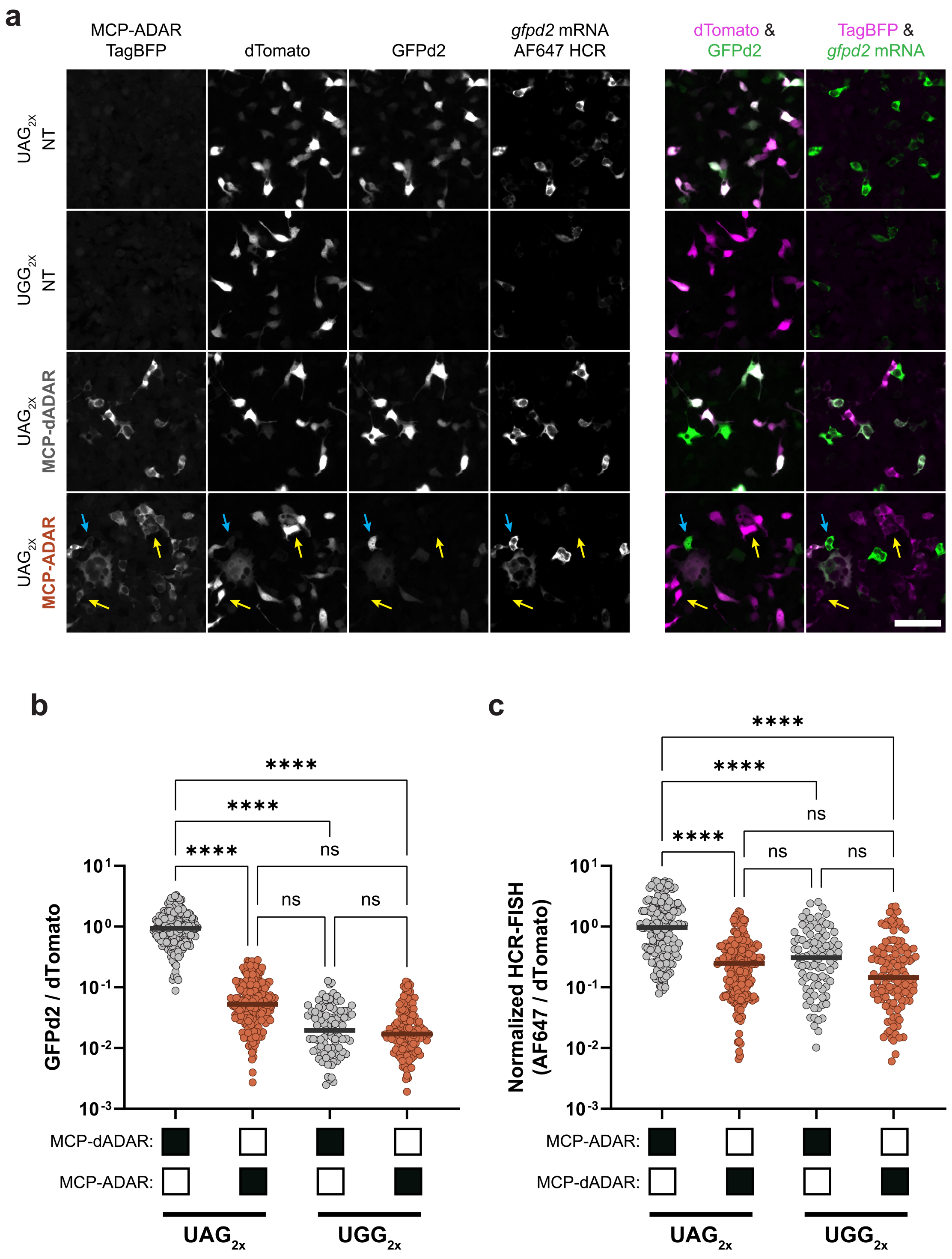


***Supplementary Figure 5 – Non-stop decay OFF-reporter editing reduces reporter protein and transcript abundance***

**a**, Representative fluorescence micrographs of HEK293FT cells co-transfected with OFF-reporters and the indicated ADAR constructs. At 24-hours post-transfection, cells were fixed and processed for HCR-FISH using probes against *gfpd2* mRNA (AF647). Cells containing constitutive readthrough $UGG_{2x}$ reporters had reduced GFPd2 emissions and *gfpd2* mRNA levels compared to those with $UAG_{2x}$ reporters. For cells containing the $UAG_{2x}$ reporter, reduced GFPd2 emissions and *gfpd2* transcripts were observed selectively in MCP-ADAR positive cells (yellow arrows indicate $tagBFP^{+}$ cells; blue arrows indicate $tagBFP^{-}$). Images are representative of 3 replicates processed in parallel. Scale bar, 100 µm.

**b**, Quantification of GFPd2 emission intensities via analysis of microscopy images using ImageJ via a custom macro. Reporter/ADAR-expressing cells were identified by dTomato and tagBFP co-expression and GFPd2/dTomato ratios were measured per cell. Cell counts per condition: $UAG_{2x}$/MCP-dADAR = 180; $UAG_{2x}$/MCP-ADAR = 220; $UGG_{2x}$/MCP-dADAR = 100; $UGG_{2x}$/MCP-ADAR = 128. Images were acquired from 3 replicates processed in parallel. Bars, median; points, individual cell measurements. Significance was assessed by one-way ANOVA (****, $P < 0.0001$; ns, not significant).
**c**, Quantification of *gfpd2* mRNA abundance using HCR-FISH (AF647) normalized to dTomato fluorescence in the same set of cells analyzed in panel **b**. Cell counts and statistical tests as in **b**. Bars, median; points, individual cell measurements. Significance was assessed by one-way ANOVA (****, $P < 0.0001$; ns, not significant).

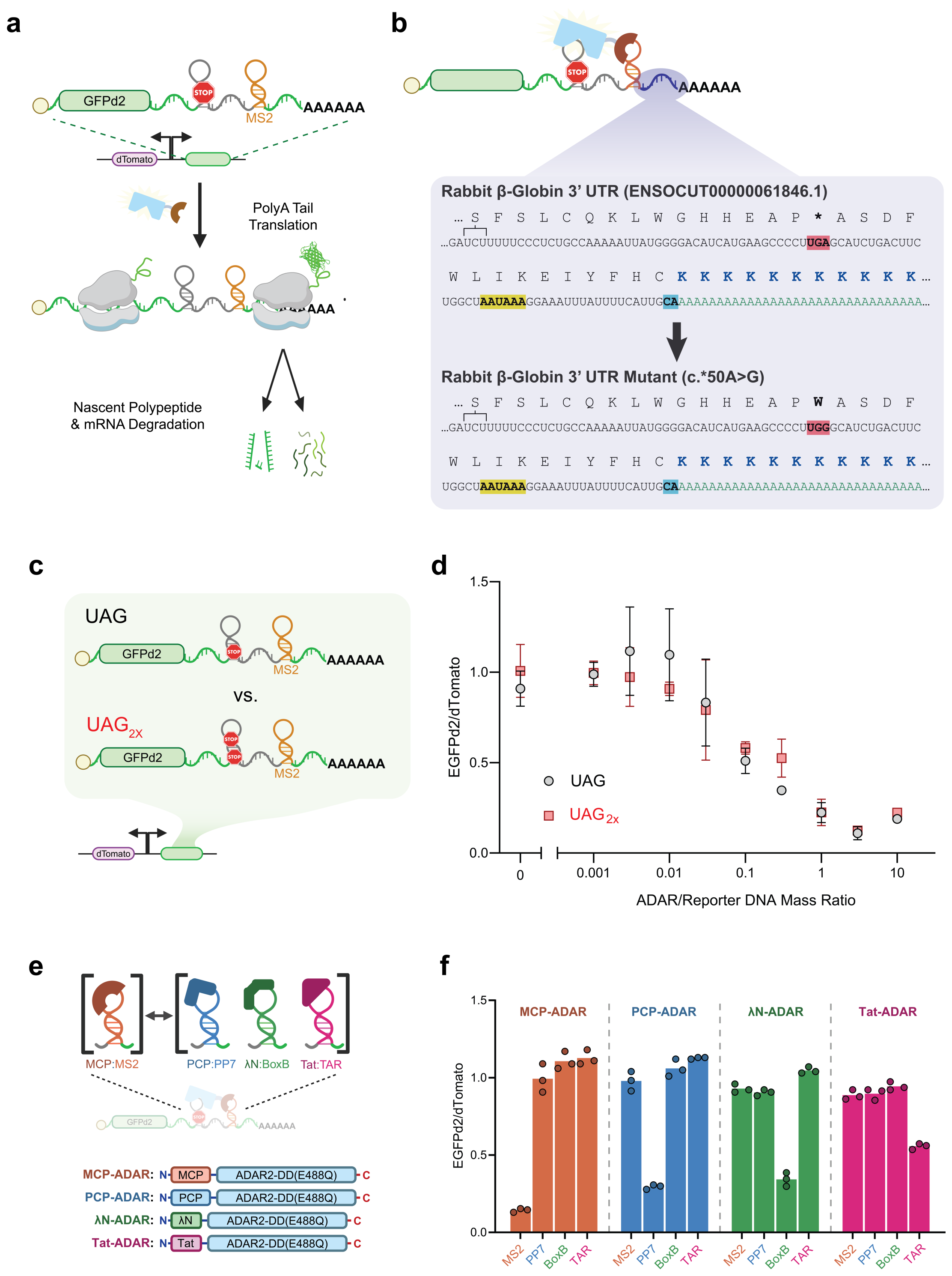


***Supplementary Figure 6 – Design of the ADAR-OFF reporter and assessment of RNA motif specificity***

**a**, Schematic of the ADAR-OFF reporter, in which editing leads to nonstop mRNA decay. Editing of the UAG stop codon to UIG facilitates ribosomal translation into the (stop codon-lacking) 3' UTR and poly(A) tail, thereby triggering transcript degradation via the nonstop decay pathway.

**b**, Mutation of a single nucleotide within the rabbit *β-globin* 3' UTR (ENSOCUT00000061846.1) results in a sequence lacking additional in-frame stop codons, thereby facilitating ribosomal translation into the poly(A) tail region upon transcript editing. An original UGA codon that is mutated to UGG in the ADAR-OFF reporter is highlighted in red, the polyadenylation signal in yellow, and the poly(A) cleavage site in cyan.

**c,** Comparison of ADAR-OFF reporters containing one (UAG) or two ($UAG_{2x}$) editable stop codons.
**d,** HEK293FT cells were co-transfected with a fixed amount (10 ng) of either UAG or $UAG_{2x}$ ADAR-OFF reporters and increasing amounts of MCP-ADAR plasmid. At 48 h post-transfection, cells were analyzed by flow cytometry. Data represent median GFPd2/dTomato ratios per replicate; points, mean; error bars, SD; *n* = 3 independent transfections.
**e,** Schematic of ADAR-OFF reporters in which the MS2/MCP interaction (orange) was replaced with alternative RNA/RBD pairs, including PP7/PCP (blue), λN/BoxB (green), and Tat/TAR (pink). Construct maps of the ADAR constructs shown below.
**f,** HEK293FT cells were co-transfected with ADAR-OFF reporters with different RNA motifs and corresponding RBD-ADAR fusions. Cells were analyzed by flow cytometry at 48 h post-transfection. Median GFPd2/dTomato fluorescence ratios are shown. Only cells bearing cognate RBD-motif pairings facilitated editing-mediated OFF reporter degradation. Bars, mean; points, replicates; n=3 independent transfections.

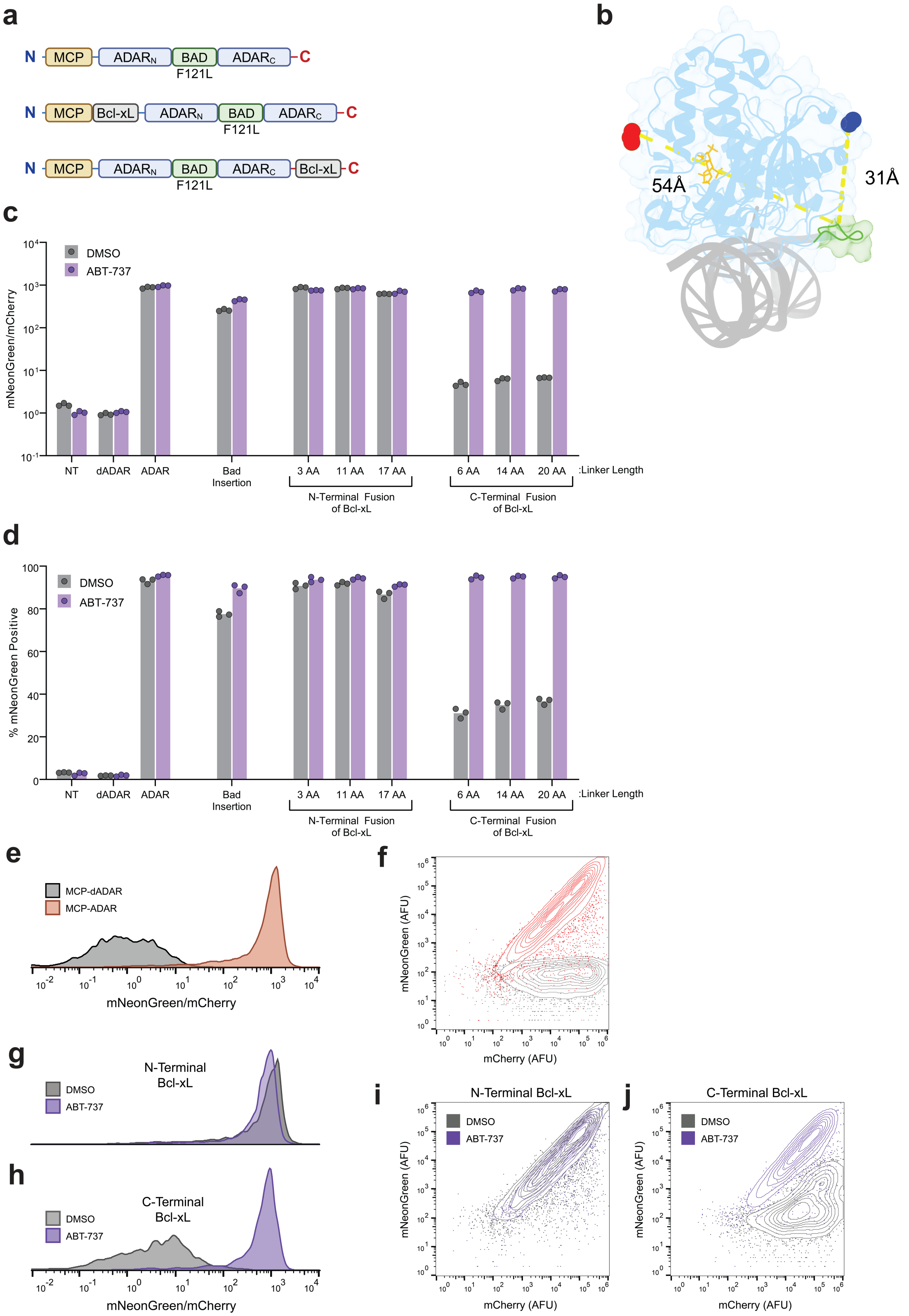


***Supplementary Figure 7 – Reversible autoinhibition of the ADAR2 deaminase domain via BAD(F121L) insertion and Bcl-xL fusion***

**a**, Construct maps of ADAR2 deaminase domain (ADAR2-DD) variants engineered for autoinhibition. Constructs include insertion of a BAD(F121L) peptide into the 5’ RNA-binding loop (between A468 and D469) and fusion of Bcl-xL to either the N- or C-terminus of ADAR2-DD using linkers of varying dimensions as indicated.

**b**, Crystal structure of the ADAR2-DD bound to dsRNA (PDB: 5ED2[4]) with the 5' RNA-binding loop highlighted (green). Distances from this loop to the N-terminus (dark blue, 31 Å) and C-terminus (red, 54 Å) are indicated.
**c,** Median mNeonGreen/mCherry fluorescence ratios measured 48 h after co-transfection of HEK293FT cells with the indicated ADAR constructs, the reporter plasmid and treatment with DMSO (gray) or 3 μM ABT-737 (purple). Bars, mean of replicates; points, replicates; n=3 independent transfections.
**d,** Percentage of mNeonGreen-positive cells from the same populations as in panel c, gated using the top 0.5% of non-transfected cells. Bars, mean of replicates; points, replicates; n=3 independent transfections.
**e–f,** Representative histogram (**e**) and contour plot (**f**) of control samples expressing inactive MCP-dADAR (gray) or active MCP-ADAR (orange), illustrating distinct inactive versus active distributions.
**g–h,** Representative histograms of N-terminal (**g**, 17-aa linker) and C-terminal (**h**, 6-aa linker) Bcl-xL fusions ± ABT-737 treatment.
**i–j,** Representative contour plots corresponding to panels g-h, showing population-level increases in mNeonGreen expression upon 3 μM ABT-737 treatment.

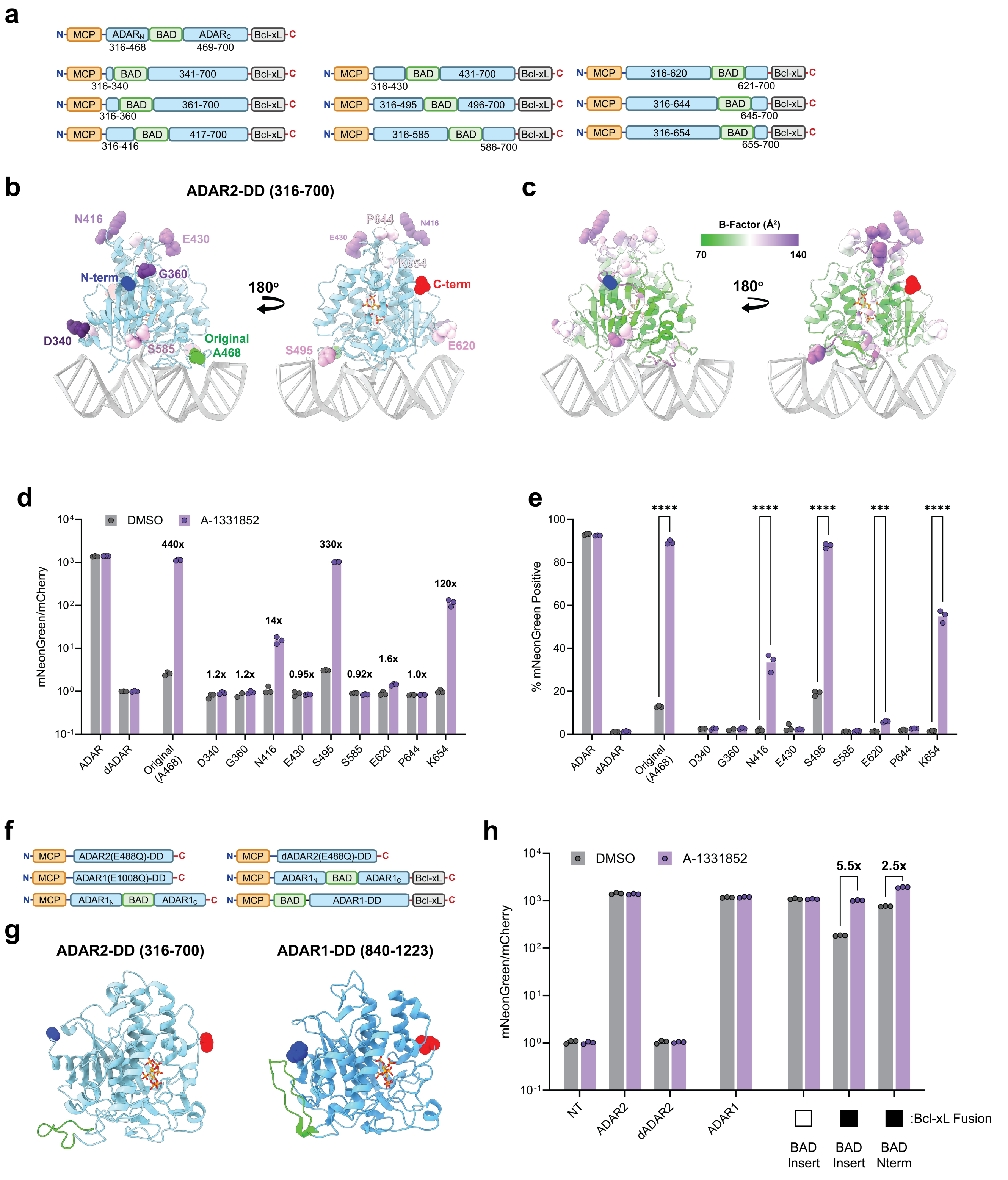


***Supplementary Figure 8 – Alternative ADAR domains and insertion sites are compatible with the iDAR strategy***

**a**, Construct maps showing alternative BAD(F121) insertion sites tested within ADAR2-DD. Each variant is named according to the residue immediately N-terminal to the inserted peptide (e.g., N416 indicates insertion between N416 and S417).

**b**, Structure of ADAR2-DD bound to dsRNA (PDB: 5ED2[4]) highlighting the N terminus (blue), C terminus (red), the original insertion site (A468, green), and all additional insertion sites (purple). Residues shown as spheres.
**c**, The same structure colored according to crystallographic B factor values ($Å^2$), indicating local backbone flexibility during structure determination. All tested insertion sites lie within solvent-exposed loops with relatively high B factors (higher flexibility), suggesting higher relative structural tolerance for peptide insertion.
**d**, Functional screening of ADAR2-DD chemiDAR variants containing alternative BAD insertion sites. HEK293FT cells were co-transfected with reporter plasmid and each variant, then treated with DMSO (gray) or 1 μM A-1331852 (purple). Median mNG/mCherry fluorescence ratios were quantified 48 h post-transfection. Variants N416, S495, and K654 exhibited drug-dependent activation exceeding 10x, while the original A468 site produced the largest dynamic range (>400x). Bars, mean; points, replicates; error bars, SD; n=3 independent transfections.
**e**, Corresponding percentage of mNeonGreen$^+$ cells for the same populations shown in **d**. Variants with strong inducibility in **d** also showed significant increases in the fraction of fluorescent cells upon drug treatment. Bars, mean; points, replicates; error bars, s.d.; n = 3. Significance determined by two-way ANOVA with multiple comparisons: ***, P < 0.001; ****, P < 0.0001.
**f**, Constructs used to test the generality of inducible ADAR (iDAR) design across ADAR paralogs. The ADAR1 deaminase domain (ADAR1-DD, residues 840-1223) was substituted for ADAR2-DD (residues 316-700) within the same MCP-BAD/Bcl-xL scaffold. The analogous site for insertion was used.
**g**, Structures of ADAR2-DD and ADAR1-DD (PDB: 5ED2[4] and 9B84[5], respectively). N termini (blue), C termini (red), $IP_6$ (orange sticks), and the 5' RNA-binding loop (green) are indicated.
**h**, Median mNeonGreen/mCherry fluorescence ratios measured 48 h after co-transfection of HEK293FT cells with the indicated ADAR constructs, the reporter plasmid and treatment with DMSO (gray) or 1 μM A-1331852 (purple). ADAR1-DD based constructs edit to comparable degrees as ADAR2-DD but have higher background and therefore lower fold changes. Bars, mean; points, replicates; n=3 independent transfections.

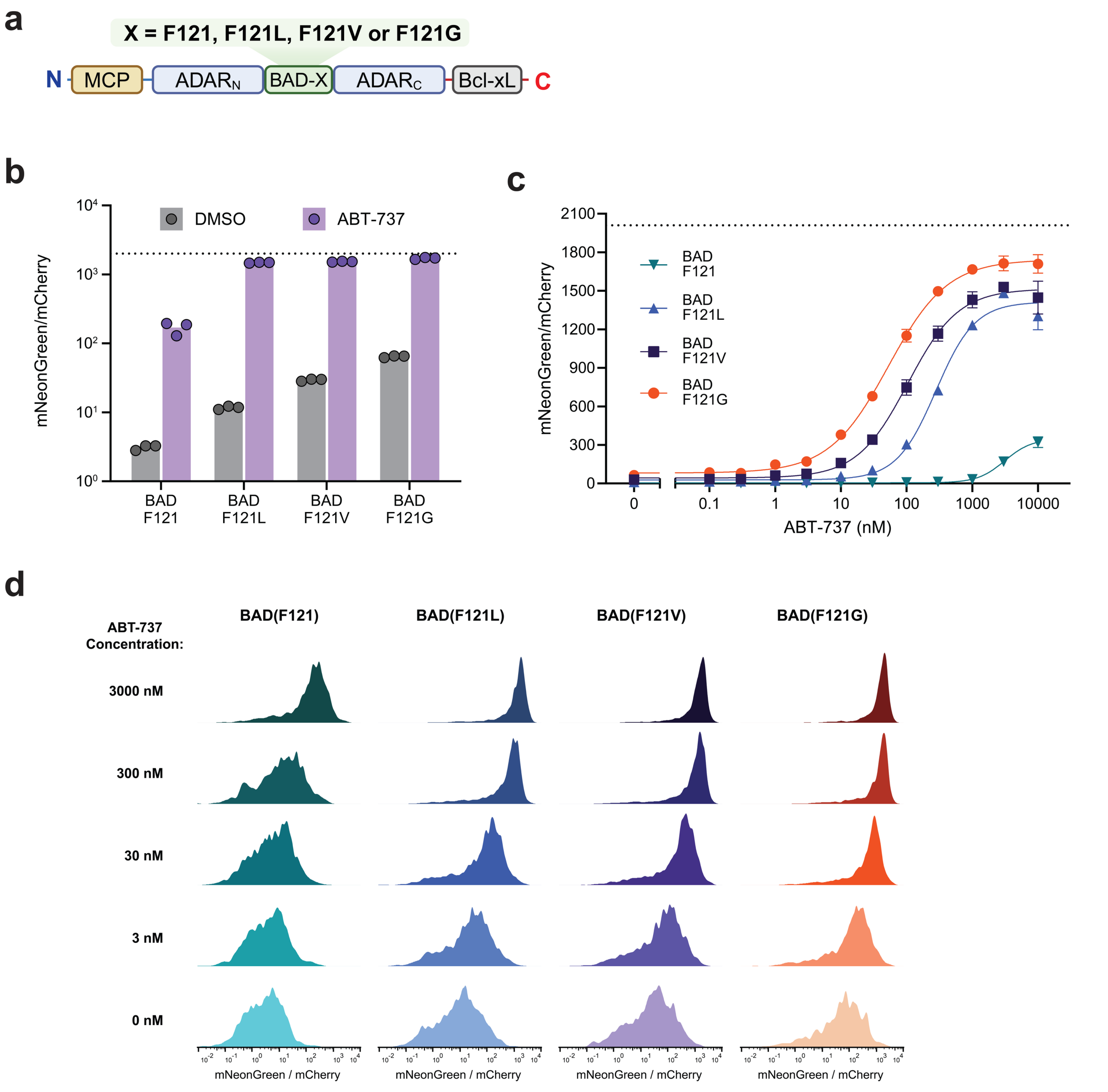


***Supplementary Figure 9 – BAD/Bcl-xL chemiDAR variants exhibit tunable, dose-dependent activation by ABT-737***

**a,** Domain architecture of chemiDAR variants containing BAD/Bcl-xL autoinhibitory domains. The F121 residue of BAD was substituted with F121L, F121V, or F121G to modulate BAD/Bcl-xL affinity.

**b,** Flow cytometry of HEK293FT cells co-transfected with ADAR-ON reporters and the indicated chemiDAR variants, treated with DMSO (gray) or 3 μM ABT-737 (purple). The dotted line indicates the mean response from MCP-ADAR positive controls, providing an estimate of maximal activation potential (n=3). Bars, mean; points, replicates; error bars, SD; n=3 independent transfections.

**c,** Dose-dependent activity of the specified BAD/Bcl-xL chemiDAR variants in response to the indicated ABT-737 concentrations. The dotted line represents mean response from MCP-ADAR positive controls (n=3). Points, mean; error bars, SD; n=3 independent transfections.

**d,** Representative histograms showing mNeonGreen/mCherry distributions for each variant across ABT-737 concentrations.

**a**

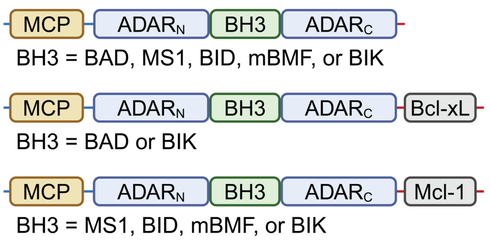


| Bcl Family | BH3 | $K_D$ (nM) | Source |
|---|---|---|---|
| Bcl-xL | BAD | 0.27 | Kong et al. |
| Bcl-xL | BIK | 102 | Kong et al. |
| Mcl-1 | MS1 | 1.9 | Foight et al. |
| Mcl-1 | BID | 87 | Kong et al. |
| Mcl-1 | mBMF | 340 | Kong et al. |
| Mcl-1 | BIK | 1100 | Kong et al. |

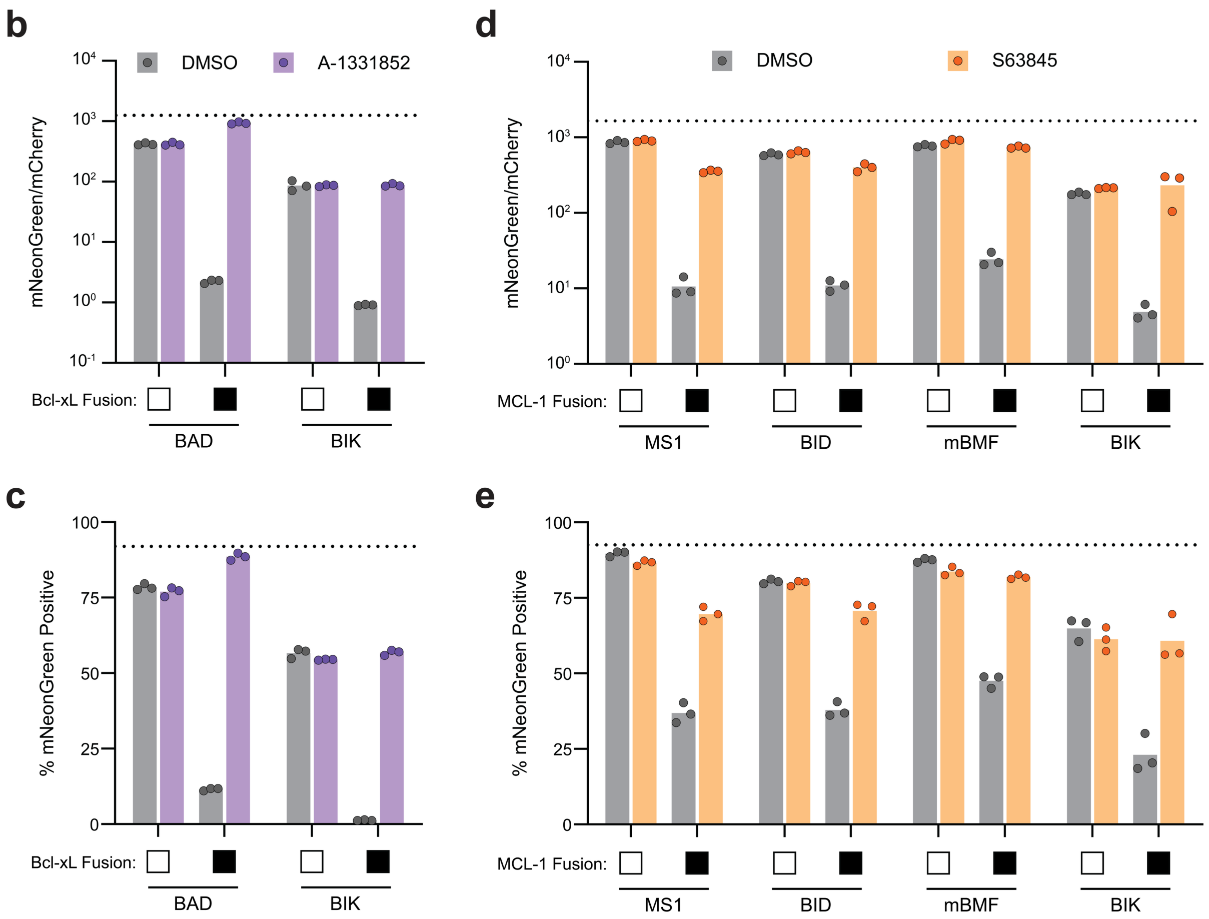


***Supplementary Figure 10 – Alternative BH3 domains with reduced affinity for Bcl-xL or Mcl-1 enable ligand-reversible autoinhibition***

**a,** Domain architectures of chemiDAR variants containing alternative BH3 peptides (BAD, MS1, BID, mBMF, or BIK) inserted into ADAR2-DD either alone or in combination with C-terminally fused Bcl-xL or Mcl-1 domains. Table (right) lists reported dissociation constants ($K_D$) for each BH3/Bcl pair, indicating relative binding affinities[6,7].

**b–c,** Activity of chemiDAR variants containing BAD or BIK BH3 peptides and Bcl-xL as the C-terminal drug-binding domain. HEK293FT cells were co-transfected with ADAR-ON reporters and the indicated chemiDAR constructs and treated for 48 h with DMSO (gray) or 1 µM A-1331852 (purple). Median mNeonGreen/mCherry fluorescence ratios (b) and percentage of mNeonGreen$^+$ cells (c) were quantified by flow cytometry.

**d–e,** Activity of chemiDAR variants containing MS1, BID, mBMF, or BIK BH3 peptides fused to Mcl-1. HEK293FT cells were co-transfected as in b-c and treated with DMSO (gray) or 3 µM S63845 (orange). Median mNeonGreen/mCherry fluorescence ratios (d) and percentage of mNeonGreen$^+$ cells (e) were quantified by flow cytometry. Dotted lines indicate the mean values from MCP-ADAR positive controls (n=3). Bars, mean; points, replicates; n=3 independent transfections.

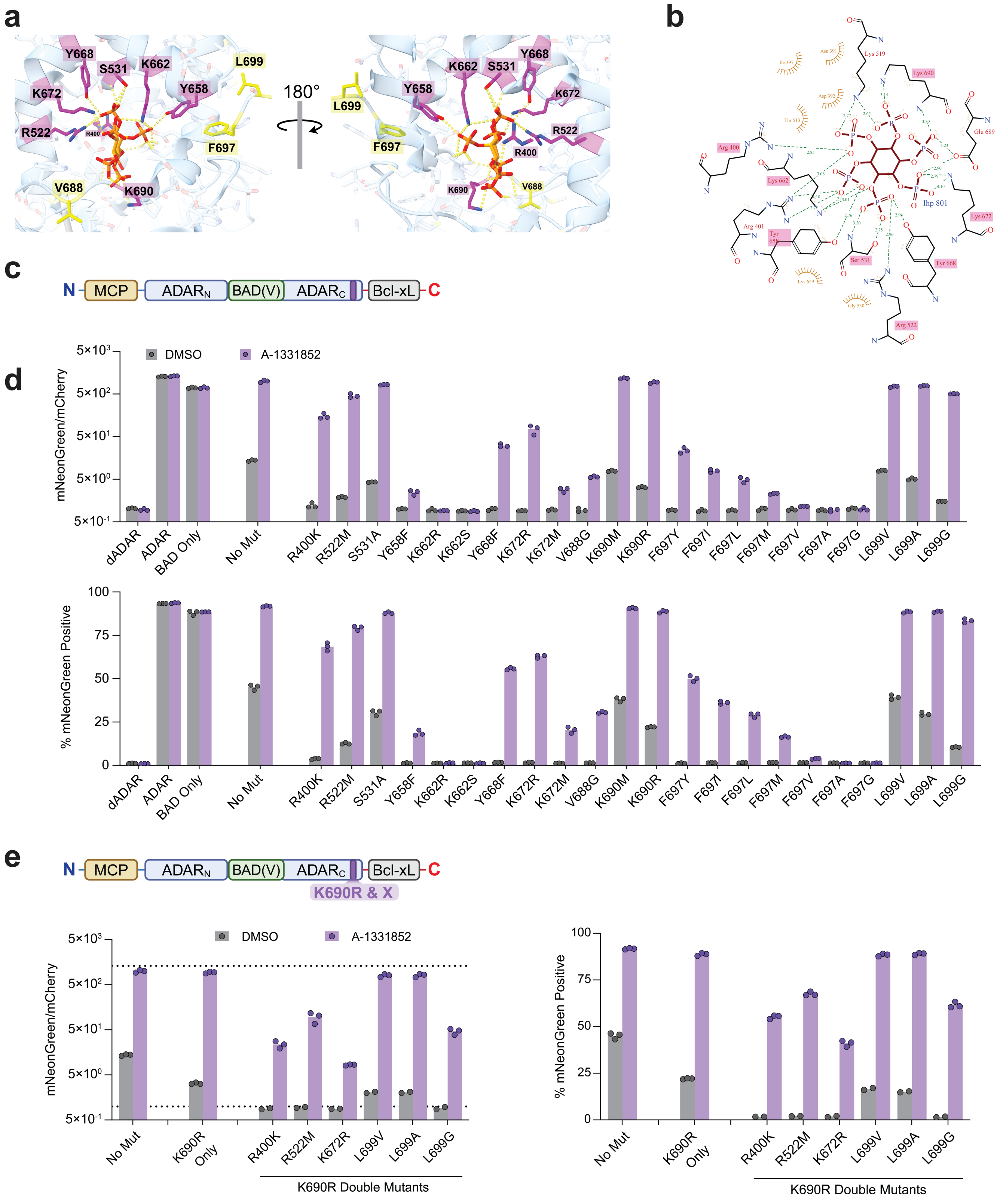


***Supplementary Figure 11 – Mutational analysis of the ADAR2 $IP_6$-binding pocket identifies variants with reduced background activity***

**a,** Structural views of the $IP_6$-binding pocket within the ADAR2 deaminase domain (PDB: 5ED2[4]). Polar sidechains directly coordinating $IP_6$ (R400, R522, S531, Y658, K662, Y668, K672, K690) are shown in magenta and hydrophobic contacts (V688, F697, L699) are shown in yellow.

**b,** LigPlot[8] diagram of the same $IP_6$-binding pocket, illustrating hydrogen-bonding and hydrophobic interactions.
**c,** Domain architecture of the BAD(F121V) chemiDAR variant used for mutational screening. Individual point mutations were introduced into the ADAR2-DD. TagBFP served as a transfection marker.
**d,** Flow cytometry of HEK293FT cells co-transfected with BAD(F121V) chemiDAR variants carrying the indicated ADAR2-DD mutations and an ADAR-ON reporter, followed by treatment with DMSO (gray) or 1 μM A-1331852 (purple) for 48 h. Median mNeonGreen/mCherry (top) and percentage of mNeonGreen$^+$ cells (bottom) are shown. Bars, mean; points, replicates; n=3 independent transfections.
**e,** Activity of BAD(F121V) chemiDAR variants containing K690R and the indicated additional $IP_6$-pocket mutations, analyzed as in panel d. Median mNeonGreen/mCherry (left) and percentage of mNeonGreen$^+$ cells (right) are shown. Dotted lines, constitutive ADAR (positive control) and dADAR (negative control). Fold changes relative to DMSO-treated conditions are indicated above bars. Bars, mean; points, replicates; n=3 independent transfections.

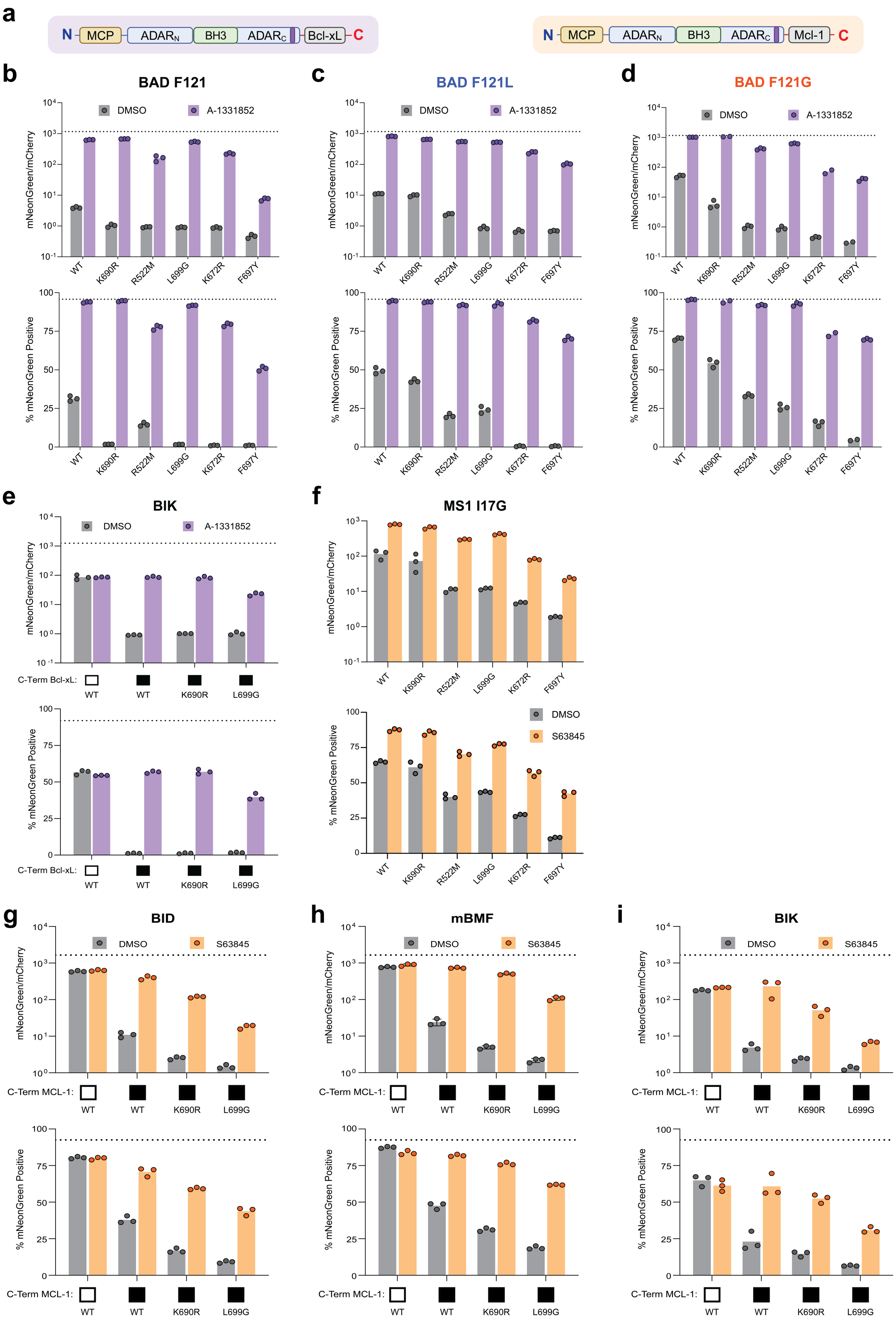


***Supplementary Figure 12 – Conserved effects of ADAR2-DD mutations across chemiDARs with diverse BH3 domains***

**a,** Domain architectures of chemiDAR constructs incorporating distinct BH3 peptides fused to Bcl-xL or Mcl-1 as autoinhibitory partners.

**b–e,** Activity of Bcl-xL-based chemiDARs containing BAD(F121) (b), BAD(F121L) (c), BAD(F121G) (d), or BIK (e) BH3 peptides. HEK293FT cells were co-transfected with the indicated variants and ADAR-ON reporters, then treated for 48 h with DMSO (gray) or 1 μM A-1331852 (purple).
**f–i,** Activity of Mcl-1-based chemiDARs containing MS1(I17G) (f), BID (g), mBMF (h), or BIK (i) BH3 peptides. Cells were treated for 48 h with DMSO (gray) or 2 μM S63845 (orange).
For each construct, reporter activity is shown as median mNeonGreen/mCherry (top) and percentage of mNeonGreen$^{+}$ cells (bottom). Fold-changes relative to DMSO-treated conditions are indicated above relevant bars. The dotted line indicates the mean response of MCP-ADAR positive controls (n=3). Bars, mean; points, replicates; n = 3 independent transfections.

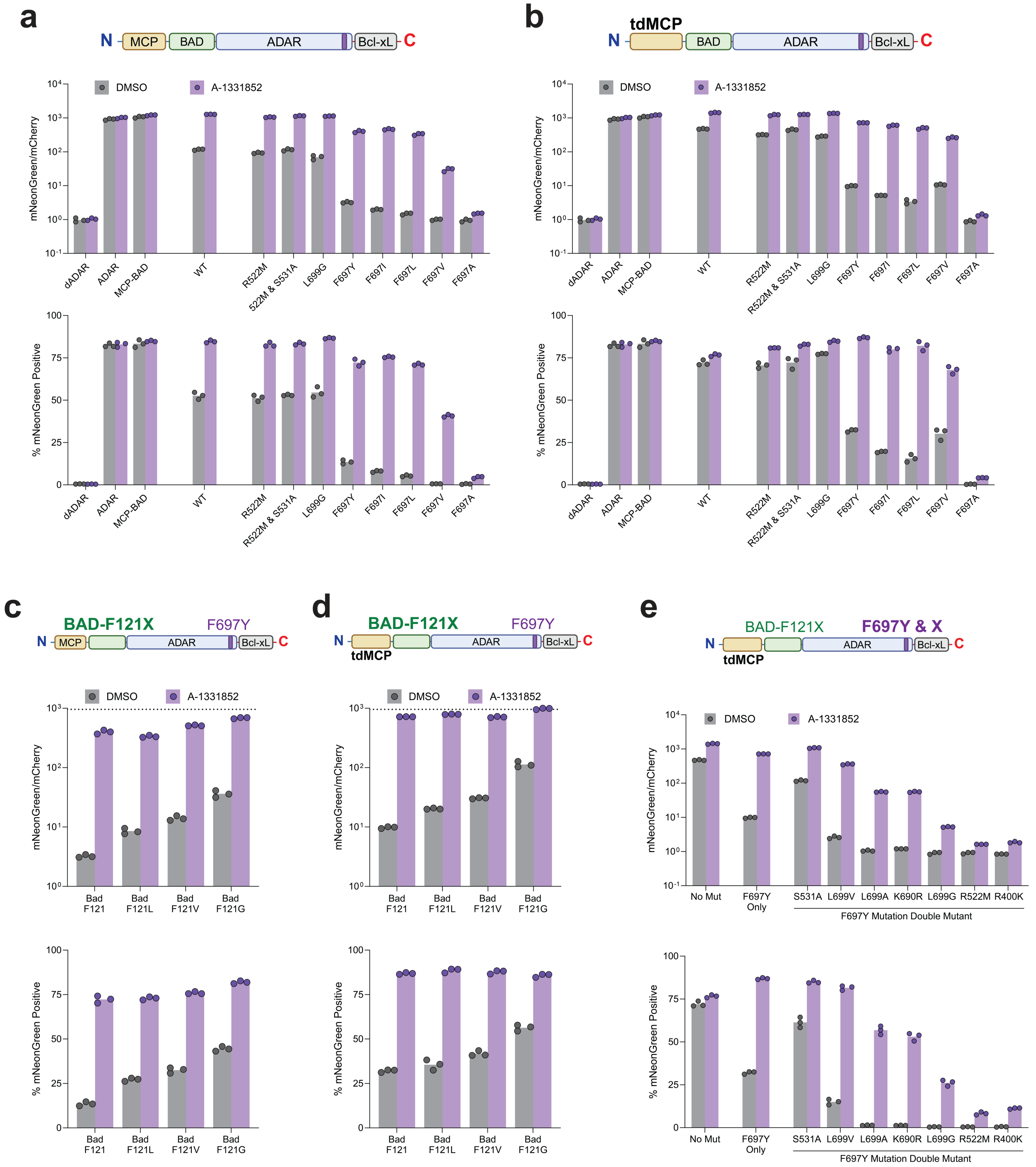


***Supplementary Figure 13 – Additional testing of IP₆-binding pocket mutations across distinct iDAR architectures***

**a–e,** Flow cytometry analysis of HEK293FT cells co-transfected with iDAR constructs containing BAD/Bcl-xL autoinhibitory domains and the indicated ADAR2-DD $IP_6$-binding pocket mutations. Cells were treated for 48 h with DMSO (gray) or 1 μM A-1331852 (purple) prior to analysis by flow cytometry. Reporter activation is shown as median mNeonGreen/mCherry fluorescence (top) and percentage of $mNeonGreen^+$ cells (bottom), with positive gates defined as the top 0.5% of fluorescence in non-transfected controls. Bars, mean; points, replicates; n = 3 independent transfections.

**a,** Dimeric MCP-based constructs containing BAD(F121) and the indicated $IP_6$-pocket mutations.

**b,** Monomeric tdMCP-based constructs containing BAD(F121) and the indicated $IP_6$-pocket mutations.

**c,** Dimeric MCP constructs incorporating the F697Y mutation together with BAD(F121X) variants (X = F121, F121L, F121V, or F121G). The dotted line indicates the mean values from MCP-ADAR positive controls (n=3).
**d,** Monomeric tandem dimer (td)MCP constructs incorporating the F697Y $IP_6$-pocket mutation and BAD(F121X) variants. The dotted line indicates the mean values from MCP-ADAR positive controls (n=3).
**e,** Monomeric tdMCP constructs containing the F697Y $IP_6$-pocket mutation in combination with the indicated added ADAR2-DD mutations (indicated below bars).

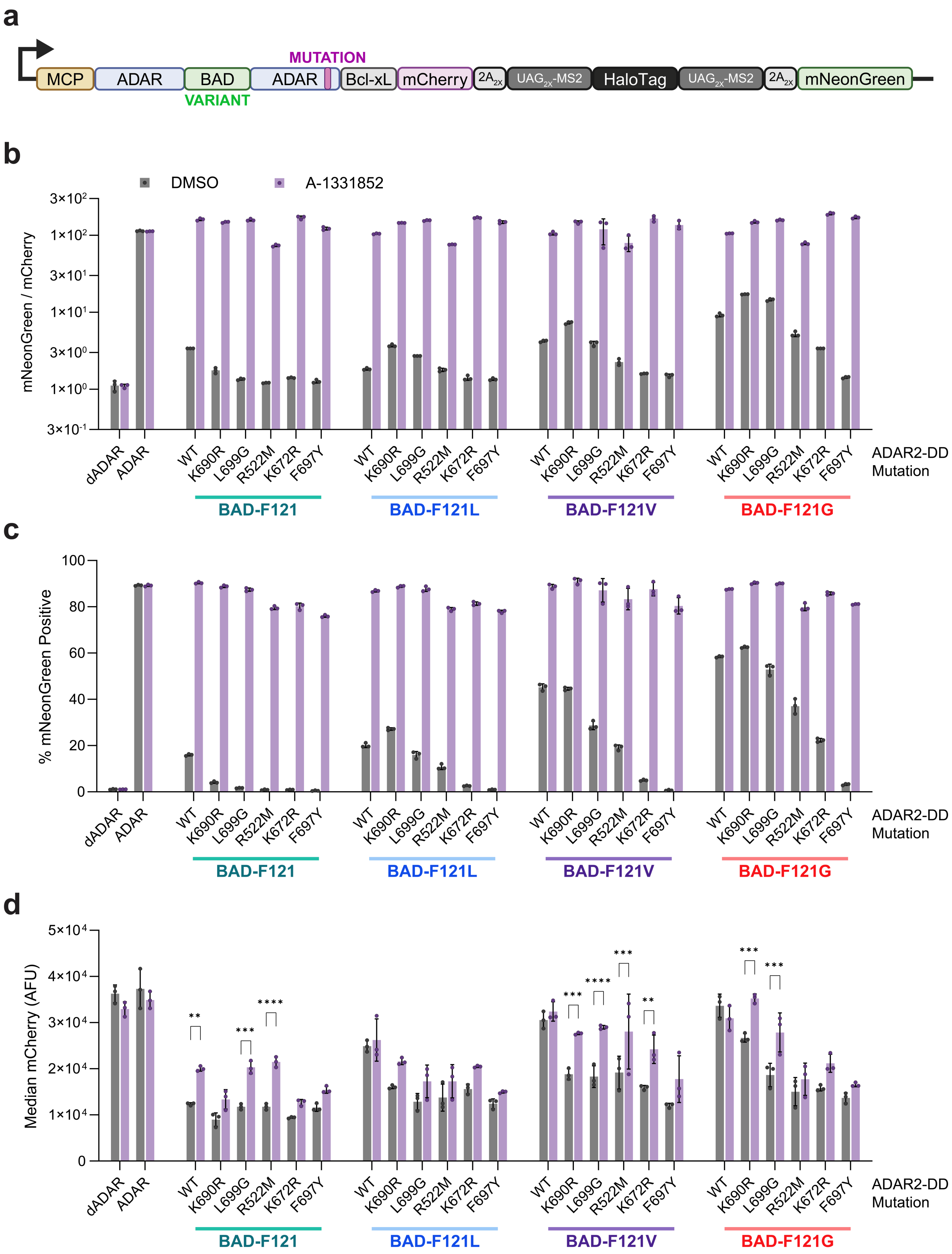


***Supplementary Figure 14 – Background activity, induction efficiency, and expression levels across self-editing iDAR circuit variants***

**a,** Schematic of the self-editing iDAR transcript architecture, in which the chemiDAR enzyme is encoded together with a corresponding editing-dependent fluorescent reporters sequence on a single transcript, with intervening 2A peptides as indicated

**b–d,** Flow cytometry analysis of HEK293FT cells co-transfected with self-editing BAD(F121X)/Bcl-xL chemiDAR circuit variants (X = F121, F121L, F121V, or F121G) containing the indicated ADAR2-DD mutations. Cells were treated for 48 h with DMSO (gray) or 1 μM A-1331852 (purple) prior to analysis by flow cytometry.
**b,** Median mNeonGreen/mCherry reporter fluorescence ratio.
**c,** Percentage of mNeonGreen$^+$ cells, with positive gates defined as the top 0.5% of fluorescence in non-transfected controls.
**d,** Median mCherry fluorescence (arbitrary fluorescence units, AFU), reporting on chemiDAR expression levels. Bars, mean; points, replicates; error bars, SD; n = 3 independent transfections. Statistical significance was determined by two-way ANOVA (*$p < 0.05$; **$p < 0.01$; ***$p < 0.001$; ****$p < 0.0001$).

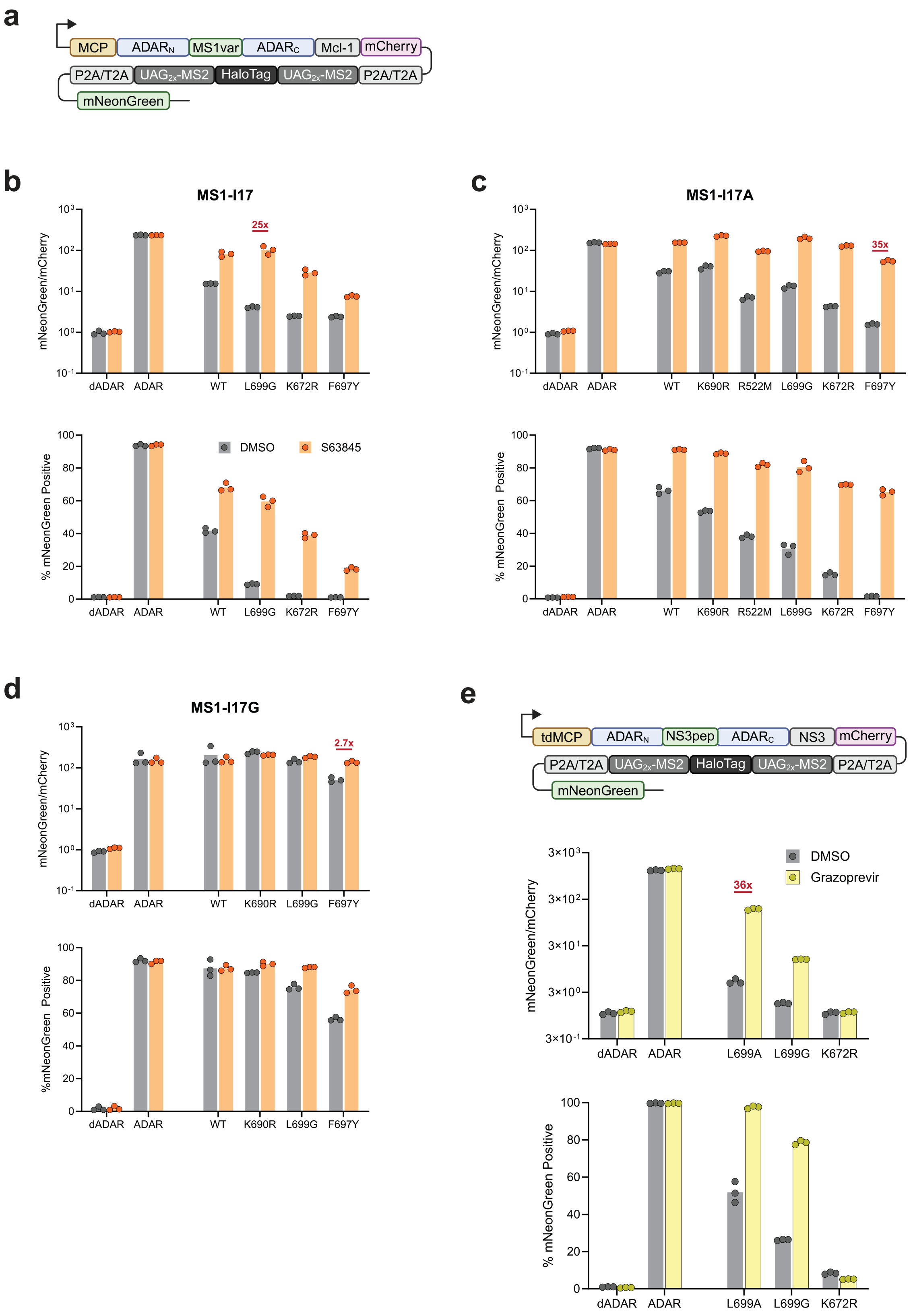


***Supplementary Figure 15 – Activities of self-editing circuits encoding chemiDAR based on MS1/Mcl-1 and NS3$_{pep}$/NS3 autoinhibitory units are enhanced by IP$_6$-pocket mutations***

**a,** Schematic of MS1/Mcl-1 self-editing circuit architecture, in which chemiDAR and fluorescent reporters are encoded on a single transcript separated by 2A peptides.

**b–d,** Activity of self-editing circuits incorporating MS1(I17X)/Mcl-1 autoinhibitory domains, measured by flow cytometry 48 h after transfection of HEK293FT cells and treatment with DMSO (gray) or 2 μM S63845 (orange). Shown are BAD(F121) variants containing the indicated ADAR2-DD mutations: **b,** MS1(I17); **c,** MS1(I17A); **d,** MS1(I17G).
**e,** Schematic and activity measurement of tdMCP-chemiDAR fusions based on $NS3_{pep}$/NS3 autoinhibitory domains, analyzed by flow cytometry following treatment with 1 μM grazoprevir (yellow) or DMSO (gray).
For all panels, reporter activity is shown as median mNeonGreen/mCherry fluorescence (top) and percentage of $mNeonGreen^{+}$ cells (bottom). Bars, mean; points, replicates; n = 3 independent transfections. Fold-change values relative to DMSO-treated controls are indicated in red where applicable.

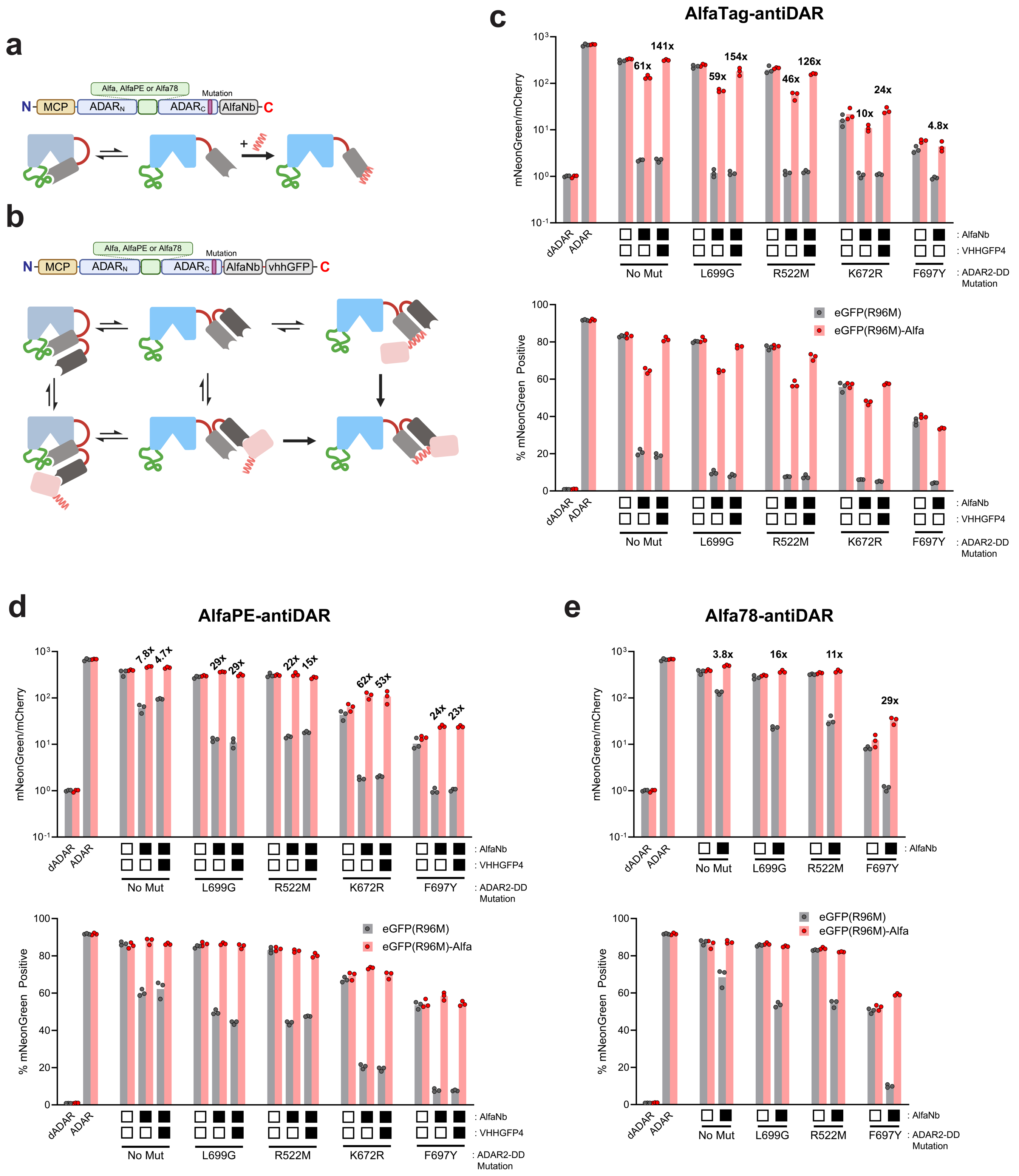


***Supplementary Figure 16 – Deaminase-domain mutations, epitope-nanobody affinity, and antibody fragment fusion modulate activity of AlfaTag-inducible antiDARs***

**a,** Schematic of the AlfaTag-antiDAR design incorporating the AlfaTag/AlfaNb pair, in which the AlfaTag epitope is inserted into the RNA-binding loop and AlfaNb is fused to the ADAR2-DD C-terminus.

**b,** Avidity-enhanced antiDAR design containing a second Nb (VHHGFP4) fused to the antiDAR C-terminus. The added Nb serves to localize the nfGFP-AlfaTag fusion antigen to the chemiDAR, thereby positioning the AlfaTag

within proximity (and with high effective local concentration) to facilitate antiDAR activation.
**c–e,** Flow cytometry analysis of HEK293FT cells co-transfected with constructs encoding nfGFP or the nfGFP-AlfaTag fusion antigen in combination with antiDAR variants containing the specified ADAR2-DD mutations. In (d-e), the inserted AlfaTag within the chemiDAR is substituted with the indicated peptide variants (AlfaPE and Alfa78); **d,** intermediate-affinity AlfaPE; **e,** low-affinity Alfa78.
Reporter activity was measured 48 h after transfection and is shown as median mNeonGreen/mCherry fluorescence (top) and the percentage of mNeonGreen$^+$ cells (bottom). Fold-change values relative to control based on co-expression with untagged nfGFP are indicated above bars. Bars, mean; points, replicates; n = 3 independent transfections.

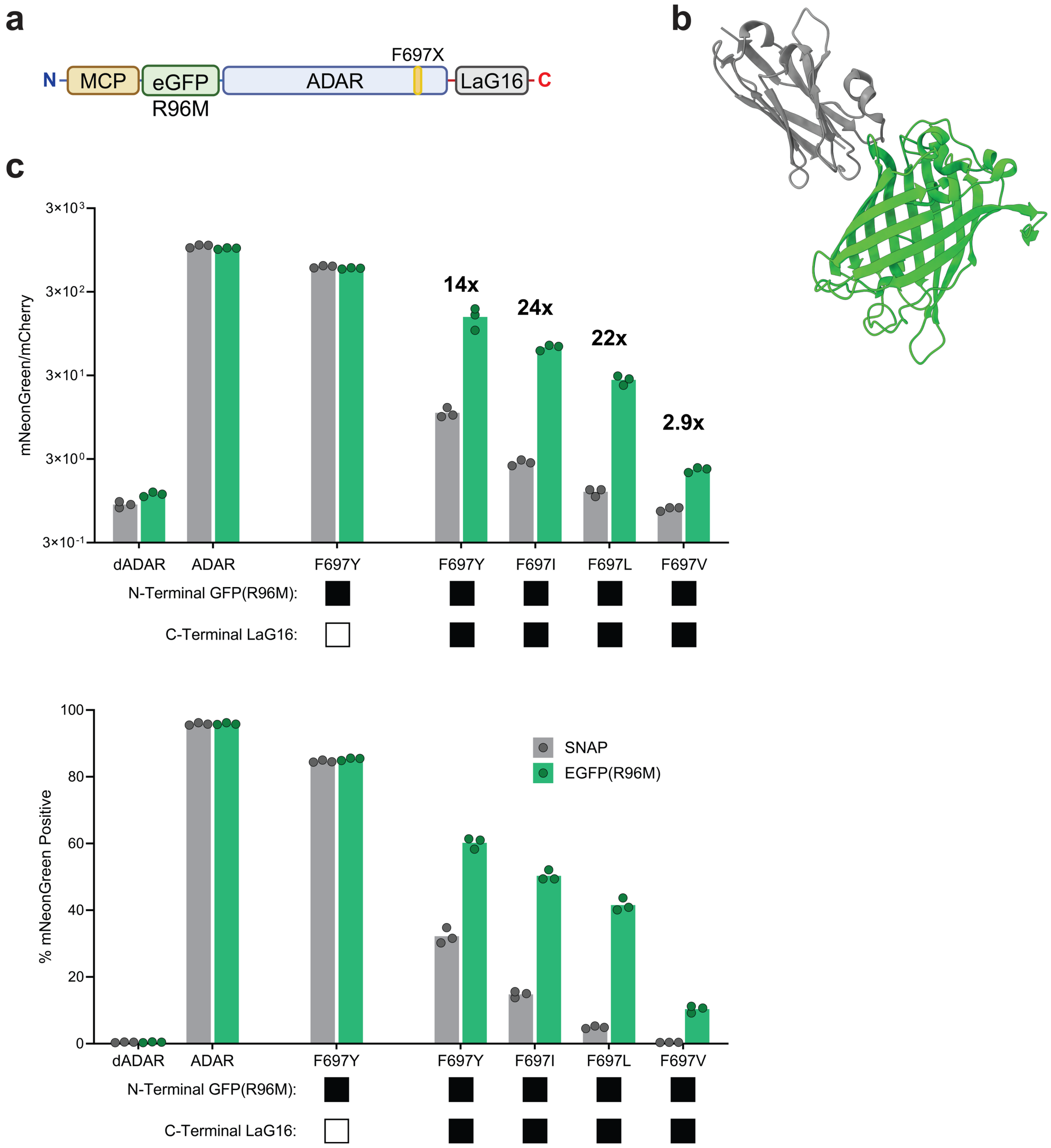


***Supplementary Figure 17 – Design and validation of antiDARs targeting a structured epitope based on GFP***

**a,** Schematic of antiDAR constructs used N-terminally with nfGFP and C-terminally with the LaG16 nanobody.
**b,** Structural model of LaG16 (gray) bound to GFP (green) (PDB: 6LR7[9]).
**c,** Flow cytometry analysis of HEK293FT cells co-transfected with constructs encoding the indicated GFP-antiDAR variants in combination with the ADAR-ON reporter, and either a SNAP domain control (gray) or a nfGFP-based antigen (green). Data were collected 48 h post-transfection and are shown as median mNeonGreen/mCherry fluorescence (top) and percentage of mNeonGreen$^+$ cells (bottom). Fold-change values relative to SNAP controls are indicated above bars. Bars, mean; points, replicates; error bars, SD; n = 3 independent transfections.

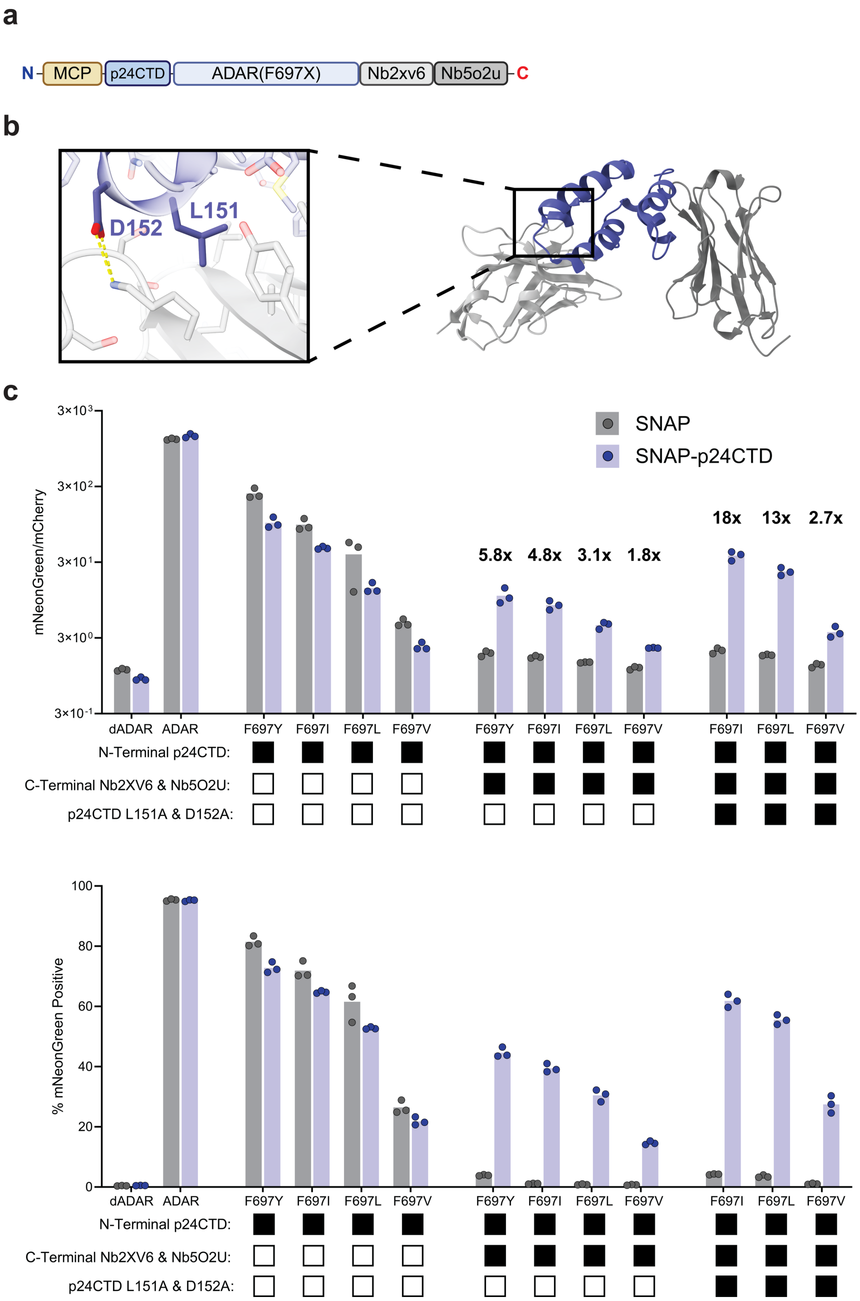


***Supplementary Figure 18 – Design and validation of p24CTD-antiDAR variants incorporating dual nanobody targeting domains***

**a,** Schematic of p24CTD-antiDAR constructs in which the ADAR2-DD is N-terminally fused with the HIV-1 p24 capsid protein C-terminal domain (p24CTD) and C-terminally linked to a dual Nbs recognizing distinct p24 epitopes (Nb2xv6 and Nb5o2u).

**b,** Structural models of p24CTD (blue) in complex with Nb5o2u (dark gray; PDB: 5O2U[10]) and Nb2xv6 (light gray; PDB: 2XV6[11]). The inset highlights L151 and D152 residues at the Nb2xv6 interface. Mutations L151A and D152A were introduced into p24CTD to weaken the p24CTD/Nb2xv6 interaction. For the mutated construct, the depicted hydrophobic and salt-bridge interactions are ablated upon alanine substitution.

**c,** Flow cytometry analysis of HEK293FT cells co-expressing the indicated p24CTD-antiDAR variants, an ADAR-ON reporter, and either a SNAP domain as a control (gray) or SNAP-p24CTD antigen fusion (blue). Data were collected 48 h post-transfection and are shown as median mNeonGreen/mCherry fluorescence (top) and percentage of mNeonGreen$^+$ cells (bottom). Fold-change values relative to SNAP controls are indicated above bars. Bars, mean; points, replicates; n = 3 independent transfections.

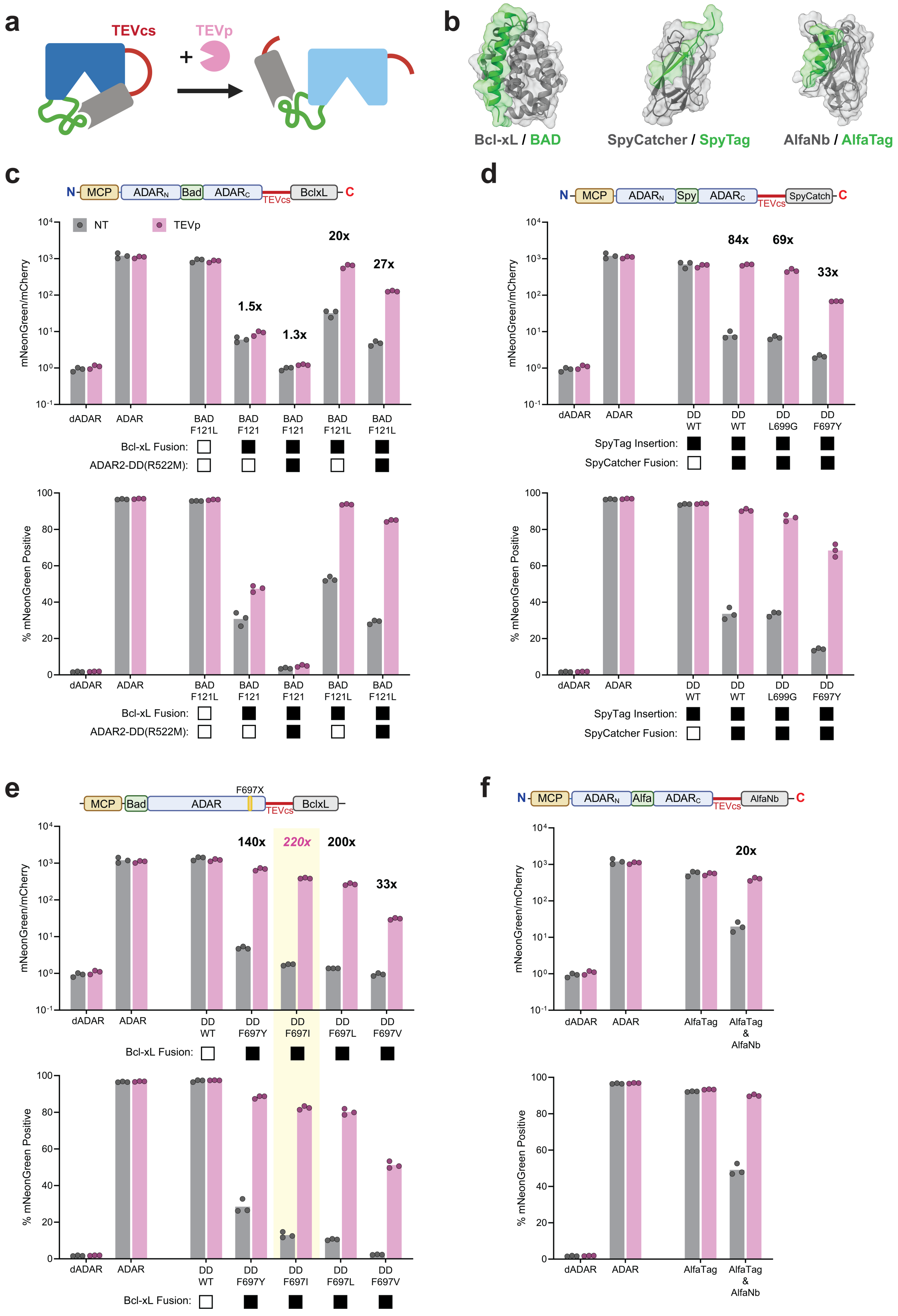


***Supplementary Figure 19 – Screening of lysiDAR constructs incorporating diverse autoinhibitory domain pairs and insertion sites***

**a,** Schematic depicting lysiDAR activation by TEV protease (TEVp) cleavage at a TEV cleavage site (TEVcs) inserted between the ADAR2-DD and C-terminally fused Bcl-xL domain.

**b,** Structural models of autoinhibitory domain pairs tested: BAD/Bcl-xL (PDB: 1G5J[12]), SpyTag/SpyCatcher (PDB: 4MLI[13]), and AlfaTag/AlfaNb (PDB: 6I2G[14]).
**c–f,** Flow cytometry analysis of HEK293FT cells co-expressing the indicated lysiDAR variants in combination with an ADAR-ON reporter, as co-transfected with either an empty vector (gray) or a TEVp-expressing plasmid (pink). Data were collected 48 h post-transfection and are shown as median mNeonGreen/mCherry fluorescence (top) and percentage of mNeonGreen$^+$ cells (bottom). Fold-change values between TEVp and control conditions are indicated above selected variants. Panels show: **c,** BAD-chemiDAR scaffold with TEVcs inserted within the RNA-binding loop; **d,** SpyTag-inserted ADAR variants inhibited by SpyCatcher fusion; **e,** N-terminal BAD-chemiDAR scaffold with TEVcs insertion; and **f,** AlfaTag/AlfaNb antiDAR with TEVcs insertion. Bars, mean; points, replicates; n = 3 independent transfections.

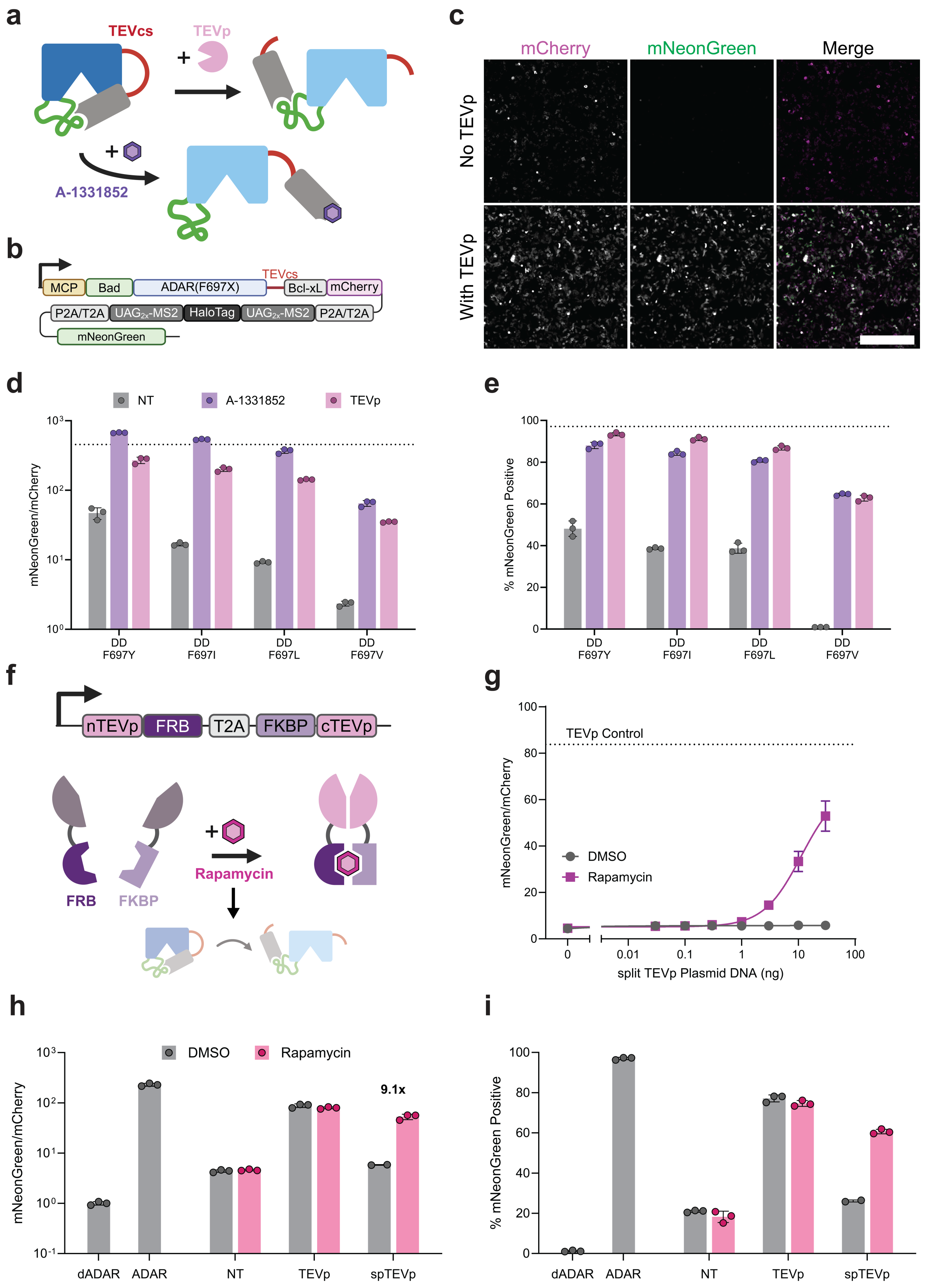


***Supplementary Figure 20 – Self-editing lysiDAR circuits facilitate protease-dependent RNA editing***

**a,** Schematic illustrating orthogonal activation modes for lysiDAR constructs via either proteolytic digestion of a TEV protease cut site (TEVcs) by TEV protease (TEVp, pink; top/right), or by ligand-induced competitive displacement of BAD and Bcl-xL through treatment with A-1331852 (purple; bottom).

**b,** Sequence organization of a lysiDAR-encoding self-editing reporter construct, in which lysiDAR activation leads to editing-mediated mNeonGreen translation.
**c,** Representative fluorescence microscopy images of HEK293FT cells transfected with self-editing construct from (b) encoding a lysiDAR-mCherry fusion bearing the F697I ADAR2-DD mutation. Cells were co-transfected alongside either empty vector DNA (top) or 3 ng of a plasmid encoding TEVp as expressed from the CMV promoter (bottom). Images were acquired 48 h post-transfection. Scale bar, 500 μm.
**d–e,** Flow cytometry analysis of HEK293FT expressing the self-editing construct from (c). The expressed lysiDAR was activated via either ligand-mediated displacement with 1 μM A-1331852 (purple bars) with or via co-expression with TEVp (pink bars). The tested lysiDARs contained the indicated F697X mutations within the ADAR2-DD. Data represent median mNeonGreen/mCherry ratios (**d**) and percentage of mNeonGreen$^+$ cells (**e**). Conditions tested: no treatment (NT, gray bars), treatment with 1 μM A-1331852 at time of transfection (purple bars), or co-transfection with 3 ng plasmid encoding TEVp (pink bars). Data are shown as median mNeonGreen/mCherry fluorescence (top) and percentage of mNeonGreen$^+$ cells (bottom). Dotted line indicates values measured from MCP-ADAR positive control. Bars, mean; points, replicates; n = 3 independent transfections.
**f,** Schematic depicting the construct used to co-express separate split-TEVp fragments as fused to FKBP and FRB (top) and cartoon showing lysiDAR activation via rapamycin-induced split-TEVp reconstitution via FKBP-FRB dimerization[15,16].
**g,** Dose-dependent lysiDAR activation measurements of cells co-expressing the construct from (**f**) following treatment with the indicated rapamycin concentration to induce split-TEVp reconstitution. HEK293FT cells were co-transfected using increasing amounts of the split-TEVp plasmid and treated with either DMSO (gray) or rapamycin at a dose of 500 nM (purple). The self-editing construct encoded a lysiDAR bearing the F697L ADAR2-DD mutation. Points and error bars, mean ± SD; n = 3 independent transfections for all conditions, except for DMSO treated cells transfected with 30 ng of plasmid, which was n = 2).
**h–i,** Control analyses showing the effect of rapamycin treatment on editing-mediated mNG translation. Flow cytometry of HEK293FT cells co-transfected with self-editing constructs encoding control MCP-ADAR or MCP-dADAR proteins (left, gray), or a lysiDAR bearing the F697L ADAR2-DD mutation (right). LysiDAR activities were evaluated under co-expression with a SNAP domain control (NT), full-length TEVp, or split-TEVp (spTEVp). Data are shown as median mNeonGreen/mCherry fluorescence ratios (**h**) and percentage of mNeonGreen$^+$ cells (**i**). LysiDAR-expressing cells were treated with DMSO (gray) or 500 nM rapamycin (hot pink) at the time of transfection. Data shown as median mNeonGreen/mCherry fluorescence ratios (top) and percentage of mNeonGreen$^+$ cells (bottom). Bars, mean; points, replicates; n = 2 or 3 independent transfections, as indicated.

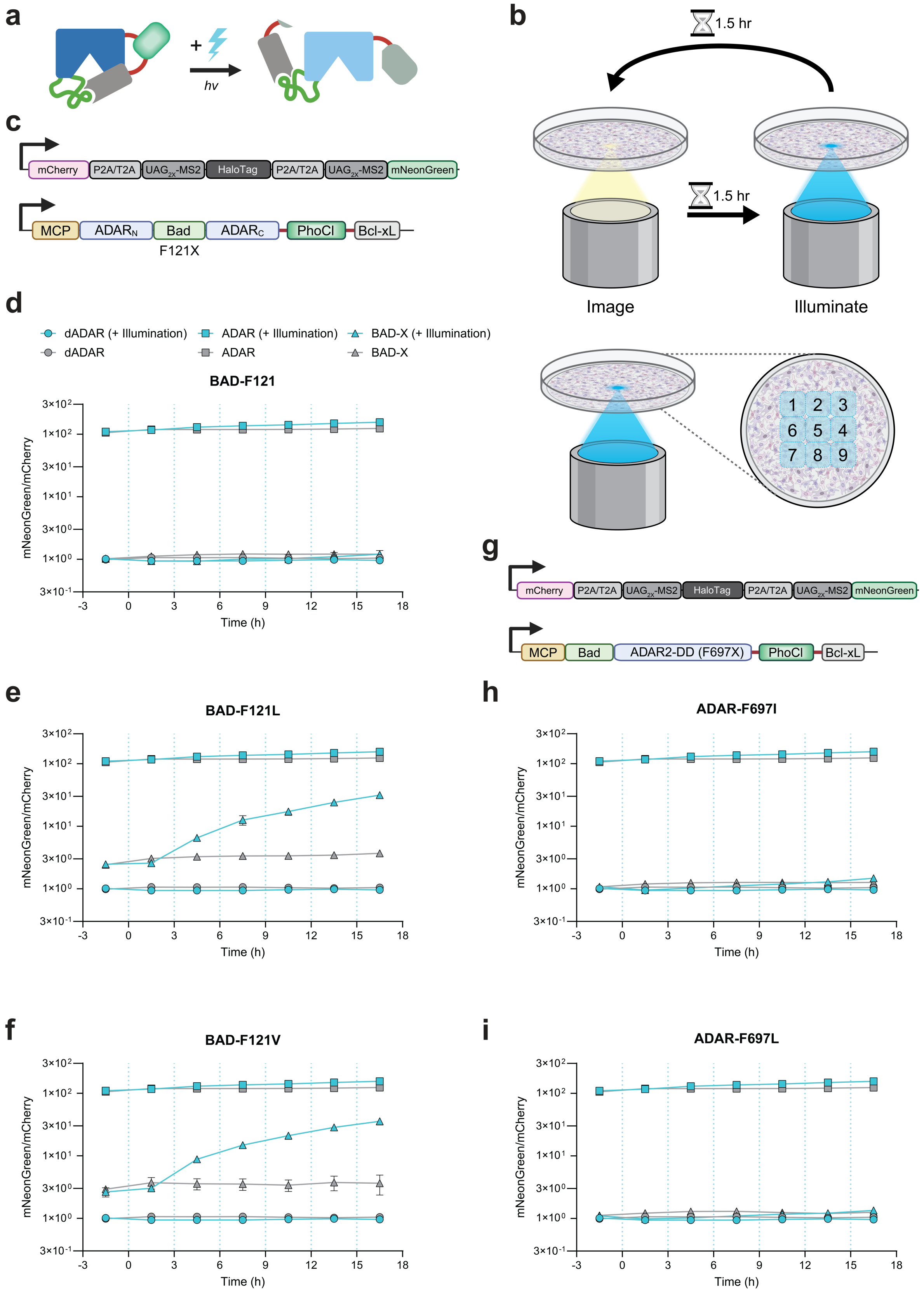


***Supplementary Figure 21 – Schematics and screening of optically activated iDAR (optiDAR) constructs***
**a,** Schematic showing the activation of optiDAR by violet light. Illumination induces photocleavage of PhoCl leading to complex dissociation, thereby relieving autoinhibition and restoring editing activity.

**b,** Experimental setup for live-cell illumination assays. At 24 h post-transfection, HEK293FT cells were transferred to a heated microscope stage. Initial mCherry and mNeonGreen emission were recorded from each well; 1.5 h later, cells were subjected to violet light illumination, as applied to the specimen through an EBFP excitation filter (ET405/20x; see Materials and Methods). Illumination was performed across a 3 × 3 tiled field of view (~2 × 2 mm) using 10 s exposure times per tile. Emissions from mCherry and mNeonGreen were recorded again at 1.5 h post-illumination. Illumination and image capture steps were carried out in series over a ~17 h duration.
**c,** Sequence schematics of separate reporter (top) and optiDAR constructs (bottom). The optiDAR construct contained autoinhibitory units based on BAD-F121X and Bcl-xL in combination with a F697X ADAR2-DD mutation.
**d–f,** Time-course analyses of BAD(F121X)-based optiDAR variants BAD-F121 (d), BAD-F121L (e), and BAD-F121V (f). Illumination occurred at 0 h and every 3 h thereafter, with intervening image capture, as outlined in (**b**). Median mNeonGreen and mCherry emissions were quantified using an ImageJ macro (see Materials and Methods), with normalized to dADAR controls (value of 1 = baseline). Gray, non-illuminated; blue, illuminated. Error bars, SD; n = 3 independent transfections. Values represent mNeonGreen/mCherry ratios.
**g-i,** Similar analyses were conducted as in (c-f) but using alternatively configured optiDAR constructs.
**g,** Sequence schematics of separate reporter (top) and optiDAR constructs (bottom). The optiDAR construct used in (**h-i**) is based on a distinct conformation (as depicted, bottom) with the addition of an F697X ADAR2-DD mutation.
**h–i,** Time-course analyses of alternatively configured optiDAR variants from (**g**) bearing the indicated F697X mutations (F697I in **h**; F697L in **i**). Cells were illuminated, imaged, and analyzed as described in (**d-f**). Symbols: circles, constitutive dADAR control; squares, constitutive ADAR control; triangles, optiDAR variants. Gray, non-illuminated; blue, illuminated. Error bars, SD; n = 3 independent transfections.

### SUPPLEMENTARY NOTES

#### Supplementary Note 1 – Custom ImageJ macro for fluorescence in situ hybridization quantification

```
// clear the log
print("\\Clear");

// to write new code for things you would like script to do: Macros>Record...

// function
function action(input, output, filename) {

    //open the image
    comp=input+filename;
    open(comp);

    //split the channels
    selectWindow(filename);
    run("Split Channels");

    //Channel C1 - TagBFP_ADAR
    //Channel C2 - GFPd2
    //Channel C3 - dTomato
    //Channel C4 - HCR - AF647
    //Channel C5 - Brightfield

    //set measurements
    run("Set Measurements...", "mean standard modal min median display redirect=None decimal=9");

    //subtract the background out of the individual images (manually measure background via ROI for 1 image to get values)
    selectWindow("C1-"+filename);
    rename("BFP");
        mode = "" + getValue("Mode");
        run("Subtract...", "value="+mode); //subtract background
    selectWindow("C2-"+filename);
    rename("GFP");
        mode = "" + getValue("Mode");
        run("Subtract...", "value="+mode); //subtract background
    selectWindow("C3-"+filename);
    rename("dT");
        mode = "" + getValue("Mode");
        run("Subtract...", "value="+mode); //subtract background
    selectWindow("C4-"+filename);
    rename("FISH");
        mode = "" + getValue("Mode");
        run("Subtract...", "value="+mode); //subtract background

    close("C5-"+filename); //close transmitted light image

    //set a threshold based upon the dTomato image
    thresh=500;
    selectWindow("dT");
    setThreshold(thresh, 65535, "raw");  //the lower bound will be the proper number to change based on images
    run("Create Mask");
```

```
run("Fill Holes");
rename("dT_mask");

//set a threshold based upon the BFP image
thresh=148;
selectWindow("BFP");
setThreshold(thresh, 65535, "raw");  //the lower bound will be the proper number to change based on images
run("Create Mask");
rename("BFP_mask");

//create AND-gate for BFP and dT positive cells
imageCalculator("AND create", "BFP_mask","dT_mask");
rename("BFP_dT_mask");

//create ROIs for individual cells that you want to measure
run("Analyze Particles...", "size=20-Infinity clear include overlay add");

//select raw HCR data and measure based on ROIs
selectWindow("FISH");
roiManager("Measure");

//save the FISH measurements to an excel file
run("Read and Write Excel", "file=["+output+"/int2_Analysis.xlsx] sheet="+filename+" dataset_label=FISH-raw");
close("Results");

//create a ratio of the FISH to BFP images
imageCalculator("Divide create 32-bit", "FISH","BFP");
selectWindow("Result of FISH");
rename("FISH-to-BFP");

//select ratios and measure based on ROIs
selectWindow("FISH-to-BFP");
roiManager("Measure");

//save the Ratio measurements to an excel file
run("Read and Write Excel", "file=["+output+"/int2_Analysis.xlsx] sheet="+filename+" dataset_label=FISH-to-BFP");
close("Results");

//create a ratio of the FISH to dT images
imageCalculator("Divide create 32-bit", "FISH","dT");
selectWindow("Result of FISH");
rename("FISH-to-dT");

//select ratios and measure based on ROIs
selectWindow("FISH-to-dT");
roiManager("Measure");

//save the Ratio measurements to an excel file
run("Read and Write Excel", "file=["+output+"/int2_Analysis.xlsx] sheet="+filename+" dataset_label=FISH-to-dT");
close("Results");

//create a ratio of the (FISH/dt) to BFP images
imageCalculator("Divide create 32-bit", "FISH-to-dT","BFP");
selectWindow("Result of FISH-to-dT");
rename("FISH/dT-to-BFP");

//select ratios and measure based on ROIs
```

```
    selectWindow("FISH/dT-to-BFP");
    roiManager("Measure");

    //save the Ratio measurements to an excel file
    run("Read and Write Excel", "file=["+output+"/int2_Analysis.xlsx] sheet="+filename+" dataset_label=FISH/dT-to-BFP");
    close("Results");

    //create a ratio of the GFP to dTomato images
    imageCalculator("Divide create 32-bit", "GFP","dT");
    selectWindow("Result of GFP");
    rename("GFP-to-dT");

    //select ratios and measure based on ROIs
    selectWindow("GFP-to-dT");
    roiManager("Measure");

    //save the Ratio measurements to an excel file
    run("Read and Write Excel", "file=["+output+"/int2_Analysis.xlsx] sheet="+filename+" dataset_label=GFP-to-dT");
    close("Results");


    //close all of the images to process next czi image
    run("Close All");


};

//'input' is the folder which contains all of the czi images
//'output' is the folder you would like to save your new images/composites

input=getDirectory("Select a Folder for Input");
output=getDirectory("Select a Folder for Output");

//for loop of 'action' function through input folder to generate 'filename' variable
setBatchMode(true);
list = getFileList(input);
for (i = 0; i < list.length; i++){
    print(list[i]);
    action(input, output, list[i]);
};
setBatchMode(false);
```

**Supplementary Note 2 – Custom ImageJ macro for optiDAR image analysis**

```
// clear the log
print("\\Clear");

// to write new code for things you would like script to do: Macros>Record...

// function
function action(input, output, filename) {

    //open the image
    comp=input+filename;
    open(comp);

    // 2. THIS IS THE FIX: Ask ImageJ "What is the name of the window you just opened?"
    var id = getTitle();
    print("ImageJ is using this window name: " + id);

    //split the channels
    selectWindow(id);
    rename(filename);
    run("Split Channels");

    //Channel C1 - mNeonGreen1
    //Channel C2 - mNeonGreen2
    //Channel C3 - mNeonGreen3
    //Channel C4 - mCherry1
    //Channel C5 - mCherry2
    //Channel C6 - mCherry3
    //Channel C7 - Brightfield


    //subtract the background out of the individual images (manually measure background via ROI for 1 image to get values)
    selectWindow("C1-"+filename);
    rename("mNG1");
        mode = "" + getValue("Mode");
        run("Subtract...", "value="+mode); //subtract background
    selectWindow("C2-"+filename);
    rename("mNG2");
        mode = "" + getValue("Mode");
        run("Subtract...", "value="+mode); //subtract background
    selectWindow("C3-"+filename);
    rename("mNG3");
        mode = "" + getValue("Mode");
        run("Subtract...", "value="+mode); //subtract background
    selectWindow("C4-"+filename);
    rename("mCh1");
        mode = "" + getValue("Mode");
        run("Subtract...", "value="+mode); //subtract background
    selectWindow("C5-"+filename);
    rename("mCh2");
        mode = "" + getValue("Mode");
        run("Subtract...", "value="+mode); //subtract background
    selectWindow("C6-"+filename);
    rename("mCh3");
```

```
    mode = "" + getValue("Mode");
    run("Subtract...", "value="+mode); //subtract background

close("C7-"+filename); //close transmitted light image

//set a threshold based upon the mCherry image
thresh=50;
selectWindow("mCh1");
setThreshold(thresh, 65535, "raw");  //the lower bound will be the proper number to change based on images
run("Create Mask");
run("Fill Holes");
run("Watershed");

//create ROIs for individual cells that you want to measure
run("Analyze Particles...", "size=100-Infinity circularity=0.1-1.00 clear include overlay add");

//create a ratio of the green and red images
imageCalculator("Divide create 32-bit", "mNG1","mCh1");
selectWindow("Result of mNG1");
rename("mNG1-to-mCh1");

//select ratios and measure based on ROIs
selectWindow("mNG1-to-mCh1");
roiManager("Measure");

//save the Ratio measurements to an excel file
run("Read and Write Excel", "file=["+output+"/"+filename+"_analysis.xlsx] dataset_label=mNG1-to-mCh1");
close("Results");

//create a ratio of the green and red images
imageCalculator("Divide create 32-bit", "mNG2","mCh1");
selectWindow("Result of mNG2");
rename("mNG2-to-mCh1");

//select ratios and measure based on ROIs
selectWindow("mNG2-to-mCh1");
roiManager("Measure");

//save the Ratio measurements to an excel file
run("Read and Write Excel", "file=["+output+"/"+filename+"_analysis.xlsx] dataset_label=mNG2-to-mCh1");
close("Results");


//create a ratio of the green and red images
imageCalculator("Divide create 32-bit", "mNG3","mCh1");
selectWindow("Result of mNG3");
rename("mNG3-to-mCh1");

//select ratios and measure based on ROIs
selectWindow("mNG3-to-mCh1");
roiManager("Measure");

//save the Ratio measurements to an excel file
run("Read and Write Excel", "file=["+output+"/"+filename+"_analysis.xlsx] dataset_label=mNG3-to-mCh1");
close("Results");

////////////////////////////////////////////////////////////////////////////
```

```
//create a ratio of the green and red images
imageCalculator("Divide create 32-bit", "mNG1","mCh2");
selectWindow("Result of mNG1");
rename("mNG1-to-mCh2");

//select ratios and measure based on ROIs
selectWindow("mNG1-to-mCh2");
roiManager("Measure");

//save the Ratio measurements to an excel file
run("Read and Write Excel", "file=["+output+"/"+filename+"_analysis.xlsx] dataset_label=mNG1-to-mCh2");
close("Results");

//create a ratio of the green and red images
imageCalculator("Divide create 32-bit", "mNG2","mCh2");
selectWindow("Result of mNG2");
rename("mNG2-to-mCh2");

//select ratios and measure based on ROIs
selectWindow("mNG2-to-mCh2");
roiManager("Measure");

//save the Ratio measurements to an excel file
run("Read and Write Excel", "file=["+output+"/"+filename+"_analysis.xlsx] dataset_label=mNG2-to-mCh2");
close("Results");


//create a ratio of the green and red images
imageCalculator("Divide create 32-bit", "mNG3","mCh2");
selectWindow("Result of mNG3");
rename("mNG3-to-mCh2");

//select ratios and measure based on ROIs
selectWindow("mNG3-to-mCh2");
roiManager("Measure");

//save the Ratio measurements to an excel file
run("Read and Write Excel", "file=["+output+"/"+filename+"_analysis.xlsx] dataset_label=mNG3-to-mCh2");
close("Results");


/////////////////////////////////////////////////////////////////////////

//create a ratio of the green and red images
imageCalculator("Divide create 32-bit", "mNG1","mCh3");
selectWindow("Result of mNG1");
rename("mNG1-to-mCh3");

//select ratios and measure based on ROIs
selectWindow("mNG1-to-mCh3");
roiManager("Measure");

//save the Ratio measurements to an excel file
run("Read and Write Excel", "file=["+output+"/"+filename+"_analysis.xlsx] dataset_label=mNG1-to-mCh3");
close("Results");
```

```
    //create a ratio of the green and red images
    imageCalculator("Divide create 32-bit", "mNG2","mCh3");
    selectWindow("Result of mNG2");
    rename("mNG2-to-mCh3");

    //select ratios and measure based on ROIs
    selectWindow("mNG2-to-mCh3");
    roiManager("Measure");

    //save the Ratio measurements to an excel file
    run("Read and Write Excel", "file=["+output+"/"+filename+"_analysis.xlsx] dataset_label=mNG2-to-mCh3");
    close("Results");


    //create a ratio of the green and red images
    imageCalculator("Divide create 32-bit", "mNG3","mCh3");
    selectWindow("Result of mNG3");
    rename("mNG3-to-mCh3");

    //select ratios and measure based on ROIs
    selectWindow("mNG3-to-mCh3");
    roiManager("Measure");

    //save the Ratio measurements to an excel file
    run("Read and Write Excel", "file=["+output+"/"+filename+"_analysis.xlsx] dataset_label=mNG3-to-mCh3");
    close("Results");


    //close all of the images to process next czi image
    run("Close All");


};

//'input' is the folder which contains all of the czi images
//'outpout' is the folder you would like to save your new images/composites

input = getDirectory("Select a Folder for Input");
output = getDirectory("Select a Folder for Output");

setBatchMode(true);

// 1. Get the list of items in the main folder
folder_list = getFileList(input);

for (i = 0; i < folder_list.length; i++) {
    // Only proceed if it is a folder
    if (endsWith(folder_list[i], "/")) {

        // Construct the path and FIX THE SLASHES
        // replace() ensures every backslash becomes a forward slash
        current_subfolder = input + folder_list[i];
        current_subfolder = replace(current_subfolder, "\\", "/");

        // 2. Get the files INSIDE that specific subfolder
        file_list = getFileList(current_subfolder);
```

```
        for (j = 0; j < file_list.length; j++) {
            if (endsWith(toLowerCase(file_list[j]), ".czi")) {

                // Clean the filename path as well
                clean_filename = replace(file_list[j], "\\", "/");

                print("Processing: " + current_subfolder + clean_filename);

                // Run your action
                action(current_subfolder, output, clean_filename);
            }
        }
    }
}

setBatchMode(false);
print("Finished all subfolders.");
```
